# Bioengineering multifunctional extracellular vesicles for targeted delivery of biologics to T cells

**DOI:** 10.1101/2022.05.14.491879

**Authors:** Devin M. Stranford, Lacy M. Simons, Katherine E. Berman, Luyi Cheng, Julius B. Lucks, Judd F. Hultquist, Joshua N. Leonard

## Abstract

Genetically modifying T cells can enable applications ranging from cancer immunotherapy to HIV treatment, yet delivery of T cell-targeted therapeutics remains challenging. Extracellular vesicles (EVs) are nanoscale particles secreted by all cells that naturally encapsulate and transfer proteins and nucleic acids, making them an attractive and clinically-relevant platform for engineering biocompatible delivery vehicles. We report a suite of technologies for genetically engineering cells to produce multifunctional EV vehicles—without employing chemical modifications that complicate biomanufacturing. We display high affinity targeting domains on the EV surface to achieve specific, efficient binding to T cells, identify a protein tag to confer active cargo loading into EVs, and display fusogenic glycoproteins to increase EV uptake and fusion with recipient cells. We demonstrate integration of these technologies by delivering Cas9-sgRNA complexes to edit primary human T cells. These approaches could enable targeting vesicles to a range of cells for the efficient delivery of cargo.

## Main text

CRISPR-Cas9 mediated genome engineering of human T cells is an area of active investigation for the development of therapeutics to treat cancer, autoimmunity, and infectious disease.^1^ Delivery of the programmable nuclease Cas9 with a single guide RNA (sgRNA) complementary to a target sequence results in the introduction of double-stranded breaks that can introduce frameshift mutations in coding genes and the ablation of protein expression. Alternately, co-delivery of a homology-directed repair template can insert specified mutations, insertions, or deletions into the genomes of target cells. While this technology has multiple applications, translation of this strategy remains difficult due to the challenges associated with *in vivo* delivery of Cas9. One approach that leverages foundational gene therapy advances is adeno-associated virus (AAV) vehicles, although safety and efficacy are often limited by anti-vector immunity and limited tissue tropism.^2-6^ Virus-like particles (VLPs) can also deliver Cas9 nucleases or base editors,^7-9^ although it remains unclear whether the immunogenicity of viral proteins will likewise limit these approaches.^10^ Synthetic nanoparticle-nucleic acid (e.g., mRNA) delivery is an alternative to viral vectors and has been successfully used for *in vivo* delivery of mRNA to confer sustained expression of chimeric antigen receptors in murine T cells.^11^ However, achieving efficient and specific T cell targeting in a manner that confers the transient expression of Cas9 needed to avoid off-target effects remains challenging.^12,13^ These general difficulties are uniquely compounded by the challenge of delivering any cargo to T cells, which exhibit low rates of endocytosis.^14^ Altogether, there exists substantial opportunity to improve delivery systems that could enable delivery of biologics to T cells inside a patient.

A promising emerging strategy is the use of extracellular vesicles (EVs) to deliver biomolecular cargo. EVs are nanoscale, membrane-enclosed particles secreted by all cells and naturally encapsulate proteins and nucleic acids during biogenesis. EVs mediate intercellular communication, delivering their contents to recipient cells to affect cellular function.^15,16^ Intrinsic properties such as non-toxicity and non-immunogenicity,^17,18^ as well as the ability to engineer surface and luminal cargo loading, make EVs an attractive platform for delivering a wide range of therapeutics. Cargo can be incorporated into vesicles either by overexpressing the cargo in the producer cells such that it is loaded during EV biogenesis or by physically or chemically modifying vesicles post-harvest.^17,19^ Cells that are genetically engineered to produce functionalized EVs may even be implanted to continuously generate such particles *in situ*.^20^ While modification of EVs post-harvest may confer cargo loading flexibility, this approach requires more extensive purification and introduces challenges from a manufacturing and regulatory standpoint.

Several recent studies have investigated the use of EVs to deliver Cas9 for treatment of cancer, hepatitis B, and genetic diseases, highlighting the promise of this method for achieving intracellular Cas9 delivery.^21-23^ However, many exploratory studies have employed EV engineering methods known to introduce artifacts in downstream experiments, which obscures how functional effects may be attributable to EVs. Of particular concern are methods that rely on transfecting EV producer cells with lipoplexes, loading EVs with electroporation methods known to result in cargo aggregation, or isolating EVs with commercial kits not intended for functional delivery applications, which have all been shown to introduce artifacts.^24-26^

Here, we address this need by developing an integrated bioengineering strategy for genetically engineering cells to direct the self-assembly and production of multifunctional EVs. As a motivating application, we systematically evaluate, compare, and generate techniques enabling EV targeting, active loading of protein cargo into EVs, and EV fusion to achieve functional cargo delivery to T cells. This exploration identifies key limitations and drivers of functional EV-mediated delivery, including a potential mechanism of receptor binding-mediated delivery enhancement to T cells. We validate these technologies by demonstrating a therapeutically relevant capability—delivering Cas9 ribonucleoprotein complex (RNP) to ablate the gene encoding the HIV co-receptor CXCR4^27,28^ in primary human CD4^+^ T cells.

## RESULTS

### Strategy for engineering multifunctional EVs for achieving delivery to T cells

Our overall approach for developing technologies toward the goal of enabling targeted delivery of biomolecules to T cells is to address each limiting step in the process (**Fig. 1**)—cargo loading into EVs during biogenesis, binding of EVs to specific target cells, uptake, and fusion of the EV with a recipient cell to release cargo into the cytoplasm. Our approach relies entirely upon genetically-encodable functions, and we term this strategy GEMINI—*G*enetically *E*ncoded *M*ultifunctional *I*ntegrated *N*anoves*i*cles.

**Fig. 1:**
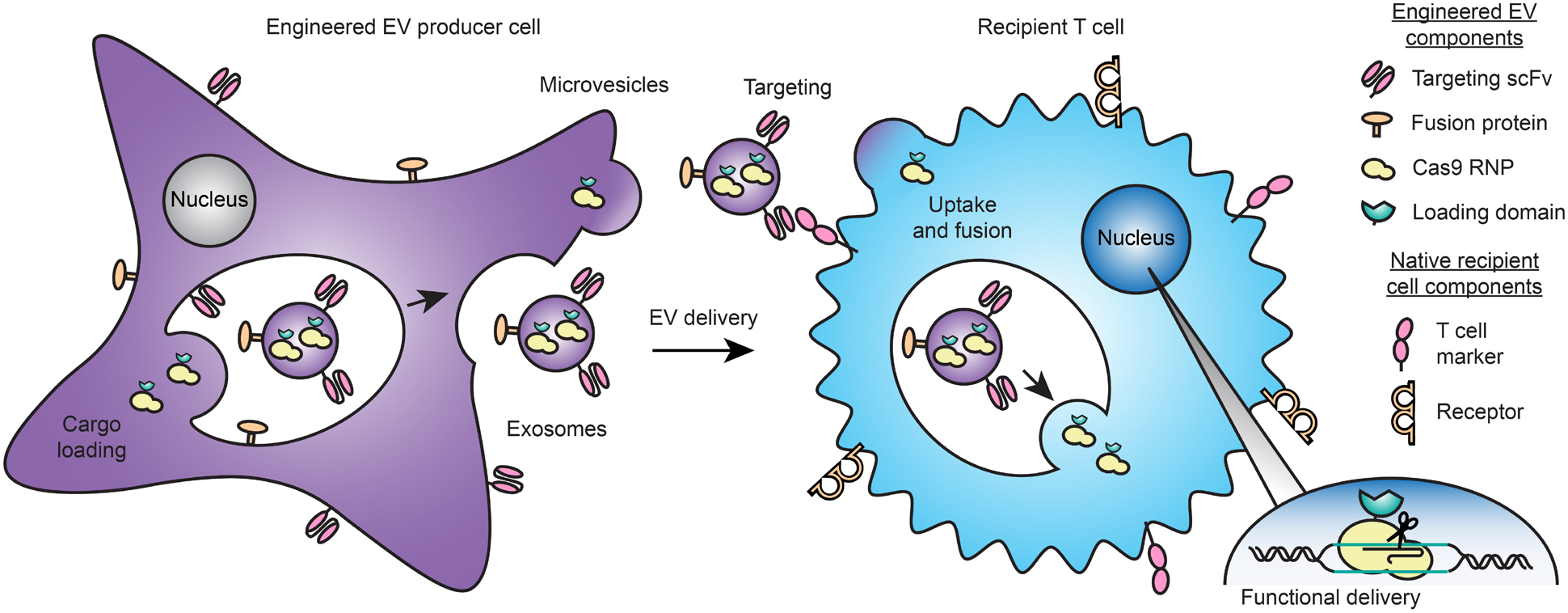
Overview of the GEMINI strategy for genetically engineering multifunctional EVs. EV cargo proteins are expressed in producer cells to facilitate incorporation into multiple vesicle populations: microvesicles, which bud from the cell surface, or exosomes, which are produced by endosomal invaginations into multivesicular bodies. Surface-displayed targeting and fusion proteins aid in binding to and uptake by recipient cells and subsequent cargo release via cell surface fusion or endosomal escape. In the proof-of-principle application explored in this study, the objective is to deliver a Cas9-sgRNA complex to T cells in order to knock out a gene, as described in subsequent sections.

### Engineered membrane scaffolds display scFvs on Evs

We first investigated strategies for conferring EV targeting. Displaying targeting moieties on the EV surface is one method to promote specific interactions between EVs and target cells and facilitate EV uptake. This strategy was pioneered using display of small peptides,^17,19^ although we and others have demonstrated that these effects are modest and variable.^29^ Recently, display of high affinity targeting domains, including nanobodies and antibody single chain variable fragments (scFvs), conferred EV targeting to receptors such as EGFR and HER2.^30-32^ We investigated whether this approach could mediate EV targeting to T cells using an anti-CD2 scFv.^33^ We selected CD2 as ligand engagement triggers internalization,^34^ and we hypothesized that such a mechanism could enhance EV uptake upon receptor docking. This could be of particular utility for conferring delivery to T cells, which exhibit low rates of endocytosis and for which delivery of other vehicles is generally challenging.^14^ We also chose to avoid targets such as CD3 which could induce non-specific T cell activation. We selected a distinct display system based upon the platelet-derived growth factor receptor (PDGFR) transmembrane domain;^19,29^ we hypothesized that using this general strategy may confer display of targeting domains on multiple EV populations. Since extravesicular linker design may impact scFv trafficking, folding, and target binding, we investigated three candidates: an α-helix to provide structure,^33^ a 40 residue glycine-serine sequence to provide flexibility, or the hinge region of IgG4 used in chimeric antigen receptors to display scFvs on synthetic receptors.^35^ All three constructs were expressed at comparable levels in HEK293FT cells (**Supplementary Fig. 1a, b**). To test display on EVs, two vesicle populations were isolated using a previously validated differential centrifugation method.^36,37^ EVs are best defined by the separation method used for their isolation;^26^ for convenience, hereafter the fraction isolated at 15,000 x g is termed “microvesicles” (MV) and the fraction isolated at 120,416 x g is termed “exosomes” (exo). Vesicles were enriched in canonical markers such as CD9, CD81, and Alix and depleted in the endoplasmic reticulum protein calnexin (**Supplementary Fig. 2a**). Both populations comprised vesicles averaging ∼120-140 nm in diameter and exhibited the expected “cup shaped” morphology (**Supplementary Fig. 2b, c**). Importantly, all three scFv display constructs were substantially expressed in both vesicle populations (**Supplementary Fig. 1c**).

### Display of anti-CD2 scFvs enhances EV binding to Jurkat T cells

To evaluate targeting, we harvested vesicles from HEK293FT producer cells stably expressing our scFv constructs and a cytosolic dTomato fluorescent protein. EVs were incubated with Jurkat T cells, which express high levels of CD2 (**Supplementary Fig. 3**), for 2 h, and then cells were washed to removed unbound vesicles (**Supplementary Fig. 4**) and analyzed by flow cytometry. All three constructs enhanced both microvesicle and exosome binding to T cells (**Fig. 2a–c**). Display of scFvs on microvesicles enhanced delivery of dTomato to T cells more so than did display on exosomes, though some— but not all—of this effect is attributable to greater dTomato incorporation in microvesicles vs exosomes (**Supplementary Fig. 5a, b**). Since the helical linker consistently conferred the greatest targeting effect, this design was carried forward for subsequent work.

**Fig. 2:**
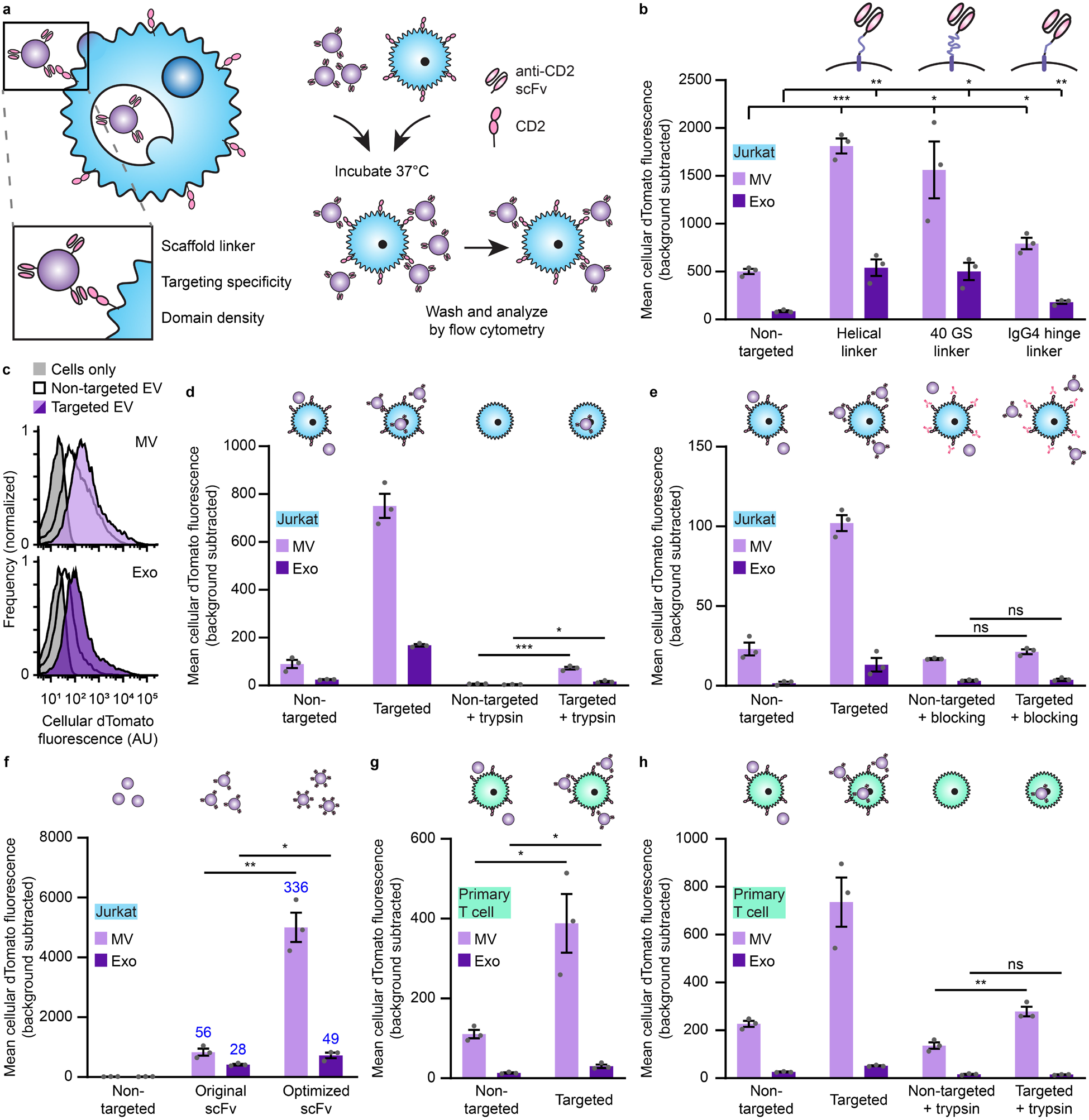
Display of scFvs on EVs mediates specific, targeted binding and uptake to T cells. **a**, Strategy for targeting EVs to T cells (left) and illustration of EV binding experiments (right). **b**, Targeted EVs binding to Jurkats (2 h incubation). To evaluate potential differences in dTomato loading, average EV fluorescence was analyzed separately (**Supplementary Fig. 5**). **c**, Representative histograms depicting distributions of helical linker EV-mediated fluorescence in recipient cells analyzed in **b. d**, Distinguishing binding and internalization for EVs targeted to Jurkats. Trypsinization was used to remove bound, non-internalized EVs following a 6 h incubation. **e**, Specificity of EV targeting to CD2. Pre-incubation with anti-CD2 antibodies ablated EV targeting to Jurkats. **f**, Enhancement of targeting by codon-optimized expression of scFv constructs. Fold increases over the non-targeted control are reported in blue. **g**, Binding of targeted EVs to primary human CD4^+^ T cells (2 h incubation). **h**, Distinguishing binding and internalization for EVs targeted to primary human CD4^+^ T cells. All experiments were performed in biological triplicate, and error bars indicate standard error of the mean. Statistical tests comprise two-tailed Student’s t-tests using the Benjamini-Hochberg method to reduce the false discovery rate. (*p < 0.05, **p < 0.01, ***p < 0.001). Exact p-values are reported in **Supplementary Table 1**. EV dTomato loading evaluations are in **Supplementary Fig. 5**.

### CD2-scFv binding mediates uptake by recipient cells

We next evaluated whether CD2 engagement by EVs triggers internalization (as noted, ligand binding naturally triggers CD2 internalization^34^). To distinguish EV binding from uptake, cells were treated with trypsin after incubation with EVs to remove non-internalized vesicles. Cells receiving targeted vesicles displayed a modest increase in fluorescence over the non-targeted control (**Fig. 2d**), indicating that CD2 targeting can mediate EV uptake.

### CD2-scFv-mediated EV targeting is specific

To determine whether the observed vesicle and T cell interactions resulted from specific receptor binding, we pre-incubated recipient Jurkat cells with an antibody binding the same T11_1_ epitope on CD2 as does our scFv to block potential binding sites. Antibody pre-treatment ablated scFv-enhanced EV binding (**Fig. 2e**), demonstrating that our targeting is specific for CD2. In contrast, pre-incubation with non-targeted EVs (a potential non-specific competitor) did not substantially reduce either background or scFv-enhanced binding (**Supplementary Fig. 6**). Together, these data indicate that the scFv mediates specific binding of EVs to CD2.

### Optimization of scFv expression increases targeting

Increasing the avidity of interactions between binders (e.g., targeted therapeutics) and their receptors is a generally useful strategy for enhancing delivery and function *in vivo*.^38^ To potentially capitalize upon this mechanism, we sought to increase the expression of our scFv constructs and therefore loading into vesicles through mass action. By optimizing the coding sequence of our scFv display construct for expression in human cells using a sliding window algorithm,^39^ we enhanced cellular expression of our scFv (**Supplementary Fig. 7a, b**) and increased scFv loading onto vesicles without affecting vesicle size or morphology (**Supplementary Fig. 7c-e**). EVs generated from cells stably expressing optimized scFv constructs exhibited enhanced specific binding to target cells (**Fig. 2f**). At the end of this limited optimization, targeted EV binding to CD2^+^ Jurkat T cells exceeded a 100-fold increase over non-targeted EVs. This optimized targeting system also conferred enhanced EV binding and modest EV internalization in primary human CD4^+^ T cells (**Fig. 2g, h**), which express high levels of CD2 (**Supplementary Fig. 8**) and was carried forward for the rest of this study.

### CD2-scFv display scaffold influences loading and specificity

Previous reports have achieved scFv display on EVs by fusion to the C1C2 lactadherin domain, which binds to phosphatidylserine on the outer membrane leaflet of some vesicles.^30,31,40^ To compare our PDGFR-based display strategy to other state-of-the-art EV scFv display systems, we fused our optimized anti-CD2 scFv to the C1C2 lactadherin domain scaffold (**Supplementary Fig. 9a**).^30,31,40^ We observed similar expression of both constructs in cells, but higher loading of C1C2 scFv constructs (as compared to PDGFR constructs) into vesicles (**Supplementary Fig. 9b, c**). Both systems conferred similar microvesicle binding to Jurkat cells (**Supplementary Fig. 9d**). C1C2 display appeared to confer some enhancement in exosome binding to T cells (compared to PDGFR display), but C1C2 display targeting was uneven, with only a subset of Jurkat recipient cells bound strongly to C1C2 display EVs, whereas PDGFR display targeting generally mediates delivery to the entire population of T cells (**Supplementary Fig. 9e**). Since this pattern might provide evidence of CD2-independent EV binding (which would comprise an artifact), we investigated whether C1C2 display targeting was specific. Pre-incubation of EVs with an anti-CD2 antibody mediated only a partial reduction in C1C2 display targeted EV binding (in contrast to PDGFR display targeting), suggesting the existence of substantial non-target-specific mechanisms for C1C2 display targeting of EVs using this scFv (**Supplementary Fig. 9f, g**). Given these observations, we opted to proceed with the validated and efficient PDGFR display of scFvs to achieve EV targeting.

### Abscisic acid-inducible dimerization domains enable an active EV cargo loading system

We next sought to engineer our scFv-containing EVs to load a therapeutic cargo of interest. Overexpression of cytosolic cargo in EV producer cells results in passive loading into vesicles during biogenesis via mass action.^41^ Increasing cargo content in EVs would potentially produce a more potent delivery vehicle. In order to both enhance cargo protein loading and increase the likelihood that a given vesicle will incorporate both a cytosolic cargo protein and our membrane-bound scFv, we designed a small molecule dimerization-based loading system (**Fig. 3a**). Systems using light or small molecules (e.g., rapamycin) as inducers have been reported to aid EV cargo loading,^42,43^ but light is difficult to scale to large volumes and rapamycin-induced dimerization is so tight that it is functionally irreversible.^44^ Therefore, we explored a new strategy based upon the plant hormone abscisic acid (ABA)-inducible interaction between truncated versions of the abscisic acid insensitive 1 (ABI) and pyrabactin resistance-like (PYL) proteins.^45^ This “ABA” system confers several advantages: association is rapid; the dimerization is reversible, presumably allowing for cargo release in recipient cells; ABA is inexpensive and non-toxic; and small molecule-regulated loading is more readily applicable to biomanufacturing than is control by light. We first investigated fusing the ABI and PYL domains to the luminal side of our scFv construct and to either the 5’ or 3’ end of a cytosolic or nuclear-localized EYFP cargo protein to determine effects on protein expression and function. Fusion with the PYL domain reduced expression (or destabilized) EYFP (**Supplementary Fig. 10a**), while the scFv was tolerant to fusions with either ABI or PYL domains (**Supplementary Fig. 10b, c**). Thus, we moved forward with the scFv-PYL and EYFP-ABI (3’ fusion) constructs. ABA-induced dimerization of ABI and PYL in this setup was readily evident by microscopy (**Fig 3b** and **Supplementary Fig. 11**).

**Fig. 3:**
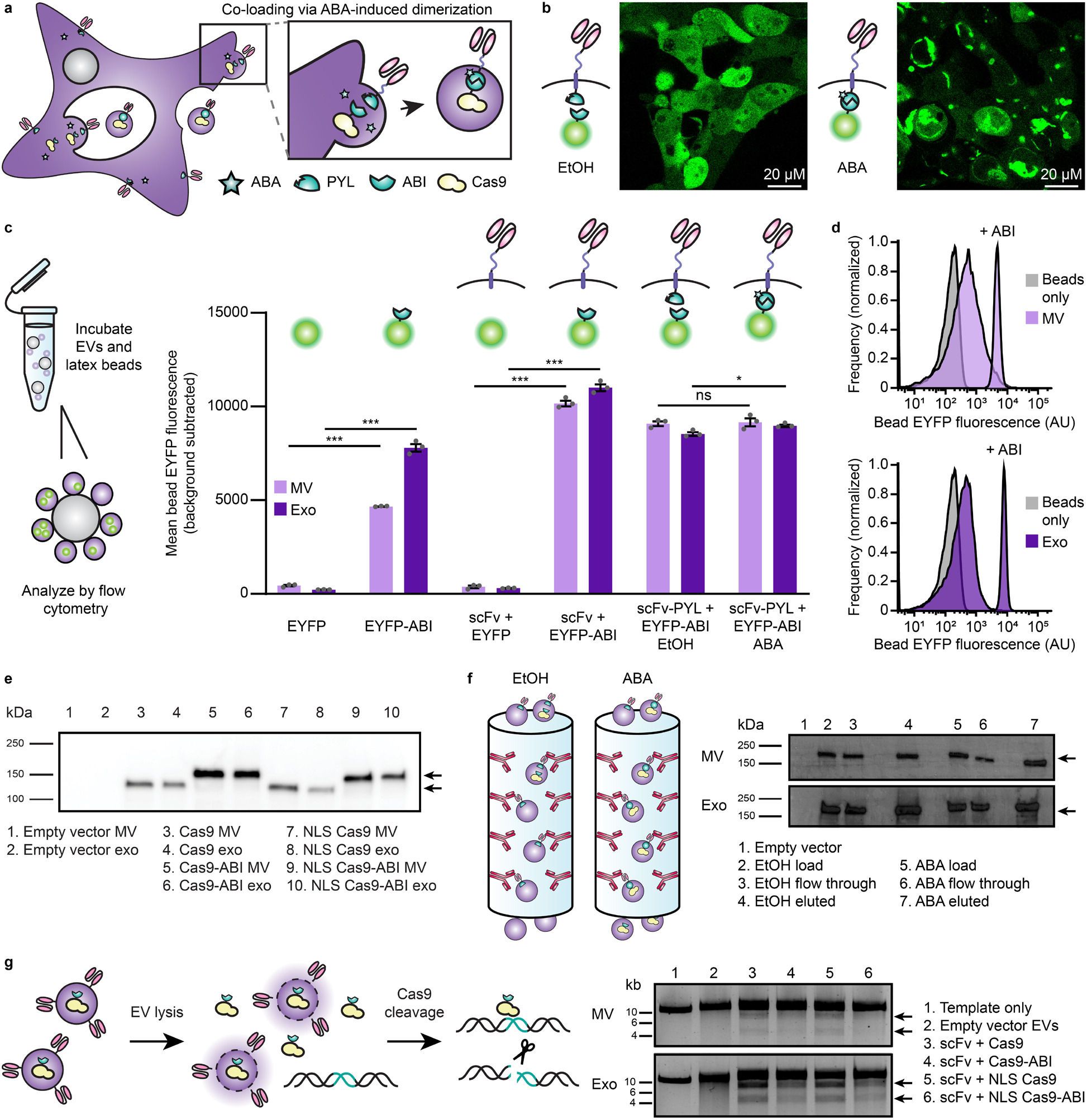
Cargo protein is actively loaded into EVs via tagging with the ABI domain of the abscisic acid dimerization system. **a**, Illustration of abscisic acid-based dimerization of EV cargo proteins and subsequent loading into vesicles. **b**, ABA-induced dimerization between PYL and ABI domains. Illustrative microscopy showing anti-CD2 scFv-PYL (membrane bound) and EYFP-ABI (cytosolic) association in the presence of ABA. Full images are in **Supplementary Fig. 11. c**, ABI-induced cargo loading into EVs. EVs generated under conditions indicated were adsorbed to aldehyde/sulfate latex beads and analyzed by flow cytometry to determine bulk average fluorescence. Experiments were performed in biological triplicate, and error bars indicate standard error of the mean. Statistical tests comprise two-tailed Student’s t tests using the Benjamini-Hochberg method to reduce the false discovery rate. (*p < 0.05, **p < 0.01, ***p < 0.001). Exact p-values are reported in **Supplementary Table 1. d**, Representative histograms of EYFP +/-ABI conditions in **c. e**, Active loading of Cas9-ABI with and without an NLS into EVs. 6.0×10^8^ EVs were loaded per lane. Expected band sizes (∼160 or 195 kDa, arrows) correspond to Cas9 -/+ the ABI domain. The full blot is provided in **Supplementary Fig. 13d. f**, Analysis of ABA-dependent Cas9-ABI loading into EVs enriched for anti-CD2 scFv-PYL via affinity chromatography. 1.3×10^7^ MVs or 2.0×10^7^ exos were loaded per lane. Expected band size: 195 kDa (arrows). Full blots are provided in **Supplementary Fig. 14b. g**, Bioactivity of EV-associated Cas9. Vesicles were lysed and incubated with a linearized target plasmid for 1 h at 37°C in Cas9 nuclease reaction buffer. Expected cut band sizes: 7.6 and 4.6 kb (arrows).

### The ABI domain alone drives protein incorporation into EVs

To investigate cargo protein loading, vesicles were adsorbed to latex beads and analyzed by flow cytometry. Surprisingly, no increase in EV loading was observed with ABA treatment, and across all conditions, constructs containing the ABI domain demonstrated a higher degree of loading than did those lacking this domain (**Fig 3c, d**). This effect was not attributable to ABI-dependent increases of protein expression in producer cells (**Supplementary Fig. 12a**). ABI-enhanced loading was evident when paired with the scFv alone or the scFv-PYL construct, indicating that intrinsic ABI-enhanced loading is independent of ABI-PYL interactions (**Fig. 3c**). The presence of the scFv conferred an added benefit in protein loading over an EYFP-ABI only control, for unknown reasons (**Supplementary Fig. 12b**). In order to investigate the role of subcellular localization on the EV loading process, we introduced a nuclear localization sequence (NLS) to EYFP-ABI and compared loading to the purely cytosolic construct. ABA-induced dimerization again had a negligible effect on cargo loading, and addition of an NLS to EYFP-ABI did not diminish loading into EVs (**Supplementary Fig. 12c**). Altogether, these data support the serendipitous discovery that ABI comprises a novel, potent EV cargo protein loading tag.

### The ABI domain mediates Cas9 loading into EVs

We next investigated whether ABI can be used to load EVs with functional cargo. For this, we selected *S. pyogenes* Cas9, as Cas9 can be synthesized in producer cells (and is thus consistent with the GEMINI strategy) and because Cas9 must travel to the nucleus of recipient cells to act on genomic targets. ABI was fused to the N- or C-terminus of Cas9, and, in general, expression patterns matched those observed for EYFP with improved expression of C-terminally tagged Cas9 (**Supplementary Fig. 13a**). Thus, we moved forward with the Cas9-ABI (3’ fusion) constructs. We also investigated whether addition of an NLS or ABI domain impacted Cas9 function. When expressed via transfection (along with a cognate sgRNA targeting a reporter construct) in reporter Jurkat T cells, Cas9 fusion constructs exhibited similar nuclease activity to Cas9 alone (**Supplementary Fig. 13b, c**). When Cas9 constructs were expressed in EV producer cells, the NLS minimally influenced Cas9 loading into EVs, while the ABI domain noticeably increased Cas9 loading (**Fig. 3e** and **Supplementary Fig. 13d**) but not overall expression in producer cells (**Supplementary Fig. 13e**). These trends are consistent with those observed with EYFP and demonstrate the utility of the ABI loading tag across multiple cargo proteins.

### Membrane scFvs and ABI-fused Cas9 co-load into EVs

An important, but largely unexplored, factor to consider in engineering EV-based therapeutics is the extent to which multiple cargo types localize to the same vesicles in a population. Although ABI (alone) successfully loads protein into EVs, it remained unknown whether dimerization of cargo and display proteins could enhance co-loading into EVs (i.e., co-loading of both the scFv and Cas9 into individual vesicles). To evaluate this question, we generated vesicles from cells expressing scFv-PYL and Cas9-ABI treated with ABA or a vehicle control and isolated anti-CD2 scFv-displaying vesicles via the 3x FLAG tag located on the N-terminus of the scFvs by affinity chromatography (**Supplementary Fig. 14a**).^46^ High levels of Cas9 were found in scFv-enriched vesicles, independent of ABA treatment, indicating that ABI-tagging of cargo is sufficient to achieve substantial scFv and Cas9 co-localization in EVs (**Fig. 3f** and **Supplementary Fig. 14b**).

### EV-loaded Cas9 exhibits nuclease function

To evaluate whether EV-encapsulated Cas9 RNPs are functional, we devised a direct *in vitro* assay. EVs from Cas9 and sgRNA-expressing cells were lysed and incubated with a plasmid encoding the sgRNA target sequence (**Fig. 3g**). Plasmids treated with lysed RNP-containing EVs showed the expected specific cleavage products under all conditions tested. The presence or absence of an NLS did not impact cleavage efficiency in this assay, but Cas9 fused to the ABI domain exhibited some reduced cleavage for both vesicle populations. Since it is not clear whether this partial effect (e.g., a potential reduction in Cas9 turnover rate) is meaningful in a cellular delivery context (further consideration in **Discussion**), both ABI+ and ABI-constructs were evaluated in subsequent experiments.

### Viral glycoprotein display increases EV uptake by T cells

To promote EV uptake and fusion, we investigated displaying viral glycoproteins on EVs. We first investigated vesicular stomatitis glycoprotein (VSV-G), which is commonly used in lentiviral pseudotyping and has been reported to confer EV fusion with recipient cells.^47,48^ VSV-G was transiently expressed in dTomato-expressing producer cells, and the resulting EVs were incubated with recipient T cells for 16 h prior to trypsinization (to remove non-internalized vesicles) and analysis by flow cytometry. VSV-G enhanced EV uptake in both Jurkat T cells (**Fig. 4a, b**) and primary human CD4^+^ T cells (**Fig. 4c**), establishing the utility in of viral fusion proteins for delivering EVs to T cells.

**Fig. 4:**
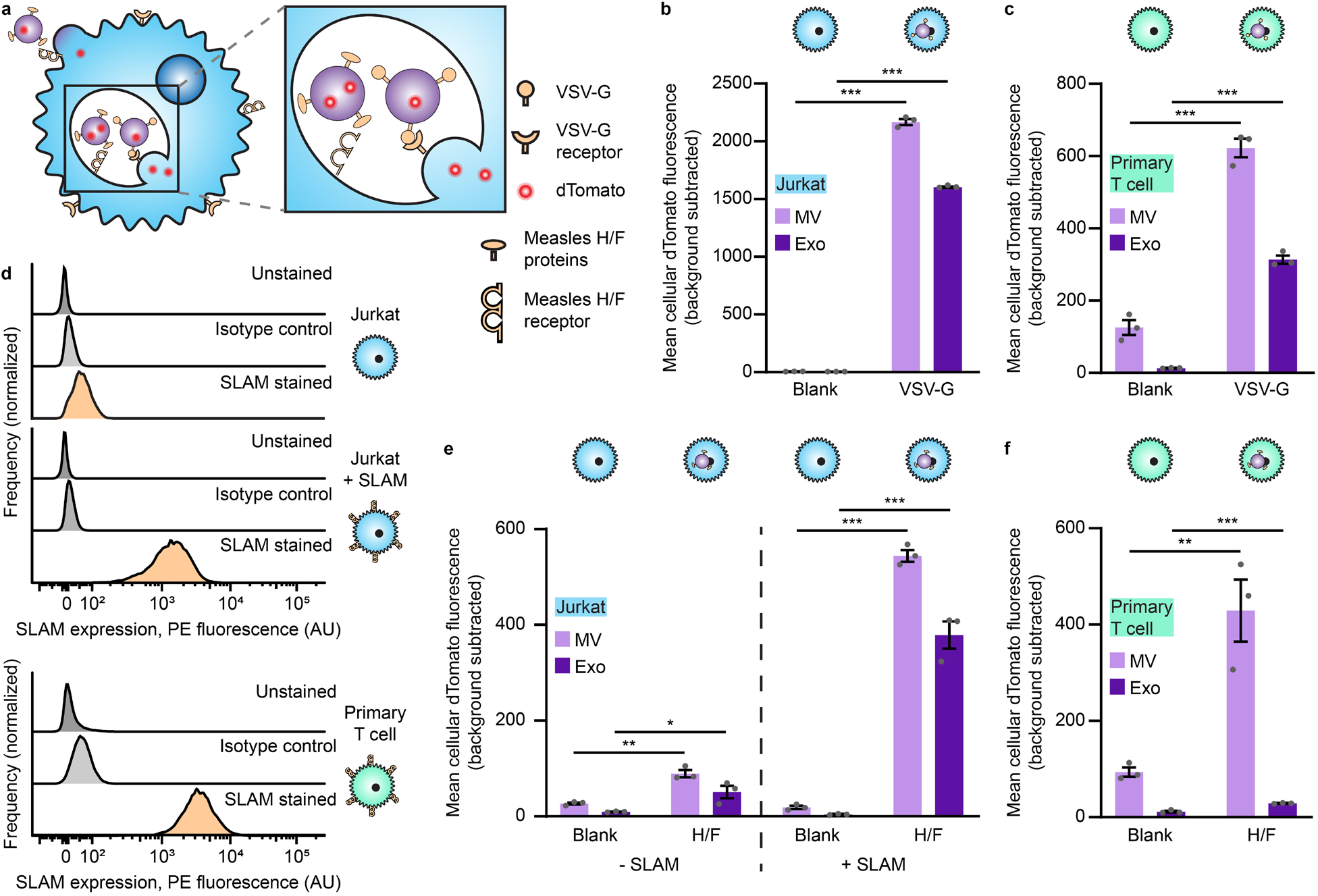
Viral glycoprotein display on EVs mediates uptake by recipient T cells. **a**, Illustration of viral glycoproteins facilitating EV uptake and fusion at either the plasma membrane or in the endosome. **b**, Uptake of dTomato-labeled VSV-G EVs by Jurkat T cells. **c**, Uptake of dTomato-labeled VSV-G EVs by primary human CD4^+^ T cells. **d**, Surface expression of SLAM on T cells. Unmodified Jurkats, Jurkats expressing transgenic SLAM, or primary human CD4^+^ T cells were evaluated for SLAM surface expression by flow cytometry. **e**, Uptake of dTomato-labeled measles viral glycoproteins H/F EVs by Jurkats (+/-SLAM). **f**, Uptake of dTomato-labeled measles virus glycoproteins H/F EVs by primary human CD4^+^ T cells. In all cases, EVs were incubated with cells for 16 h and trypsinized to remove surface-bound vesicles. Experiments were performed in biological triplicate, and error bars indicate standard error of the mean. Statistical tests comprise two-tailed Student’s t tests using the Benjamini-Hochberg method to reduce the false discovery rate. (*p < 0.05, **p < 0.01, ***p < 0.001). Exact p-values are shown in **Supplementary Table 1**. EV dTomato loading evaluations are in **Supplementary Fig. 15**.

To develop an EV fusion system that is more specific to T cells (since VSV-G mediates fusion to most cell types),^49^ we investigated the use of truncated versions of the measles virus glycoproteins H and F, which have previously been used to aid lentiviral delivery to T cells.^50,51^ These proteins bind signaling lymphocyte activation molecule F1 (SLAM) and/or the complement regulator CD46, both of which are expressed on diverse T cells.^52^ H/F proteins are classically believed to mediate viral fusion at the cell surface,^53^ although it has also been reported that viral endocytosis can be mediated by SLAM.^51,54^ In the same fluorescent EV uptake assay described above, we investigated EV delivery to Jurkats (which minimally express SLAM), SLAM-transgenic Jurkats, or primary human T cells that express SLAM (**Fig. 4d**). H/F proteins conferred modest EV uptake to parental Jurkats (SLAM-), but these proteins substantially enhanced EV uptake by SLAM-transgenic Jurkats and primary human CD4^+^ T cells (**Fig. 4e, f**). We also explored an alternative, non-viral protein-based strategy reported to promote functional transfer by overexpressing constitutively active Cx43, a connexin protein involved in the formation of gap junctions, on EV producer cells.^20,55^ Cx43 did not confer increased EV internalization by T cells in this application, so this approach was not further investigated (**Supplementary Fig. 16**). Altogether, these results support the use of the measles H/F glycoproteins as a method for enhancing EV uptake by SLAM+ T cells.

### EVs mediate functional delivery of Cas9 to primary T cells

Evaluating functional delivery of Cas9 to recipient T cells requires effective cargo loading, T cell binding and fusion, and subsequent release of active Cas9 RNPs, and having validated each of these steps individually, we proceeded to evaluate the combined technologies—the first combined test of the GEMINI strategy. Specifically, we investigated the use of Cas9 to target the CXCR4 locus in primary T cells using a previously validated sgRNA; CXCR4 is a clinically-relevant target for the treatment of HIV.^56-58^ Since viral glycoprotein expression is cytotoxic, at this point we pivoted to biomanufacturing EVs using a Lenti-X HEK293T cell line that is well-suited to this challenge; this line was selected for its ability to produce high lentiviral titers. EVs containing the anti-CD2 scFv, NLS Cas9-ABI with the appropriate sgRNA, and either VSV-G or measles virus glycoproteins H/ F were incubated with primary human CD4^+^ T cells for 6 d before harvesting genomic DNA for high throughput sequencing (HTS) to quantify and characterize targeted edits in a region of 64 nucleotides centered around the expected cleavage site. Excitingly, indels were identified at the predicted Cas9 cut site for all vesicle treatments containing Cas9 RNPs (**Fig. 5** and **Supplementary Fig. 17**). VSV-G display on EVs conferred higher editing efficiencies than did measles H and F proteins, and exosome treatments conferred more edits than did microvesicle treatments for matched designs. The majority of edits were classified as deletions with a smaller number of insertion events or edits consisting of both an insertion and a deletion. This overall pattern is consistent with prior reports of Cas9 RNP editing at this locus,^56^ in that edits comprise mostly small deletions and insertions centered around the cleavage locus, indicating that EV-mediated delivery of Cas9 using GEMINI yields effects that are qualitatively comparable to electroporation of recombinant Cas9 RNPs. In order to evaluate the role of ABI-mediated active loading in functional delivery, we generated EVs with Cas9 +/-ABI and evaluated editing efficiencies in primary T cells. The two Cas9 variants performed comparably well in this context, despite previously noted tradeoffs in loading and specific cleavage activity (**Supplementary Fig. 18**).

**Fig. 5:**
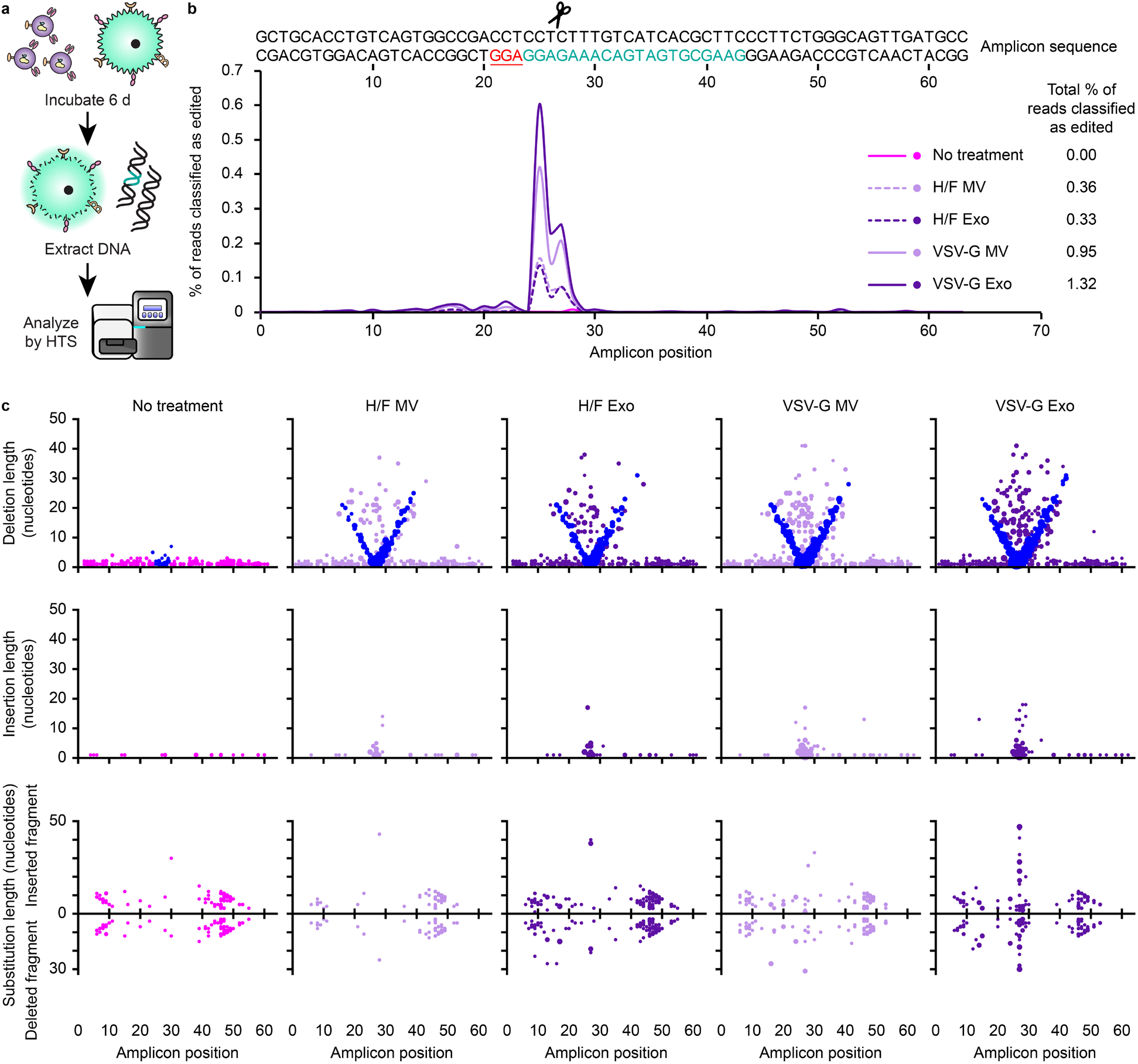
EVs mediate functional delivery of Cas9 in primary human T cells. **a**, Illustration of function delivery evaluation. 2.0×10^10^ EVs were incubated with 5.0×10^4^ CD4^+^ T cells for 6 d prior to genomic DNA extraction and HTS analysis. **b**, Frequency of indels detected at the Cas9-targeted CXCR4 locus. The sgRNA recognition site (green), PAM sequence (underlined, red), and predicted cut site (amplicon position 26, scissors) are shown. Total percentage of HTS reads classified as “edited” represents the area under the histogram trace shown for each sample. **c**, Distributions of EV-Cas9-mediated edits, by type. DNA amplicon position is plotted on the abscissa and length of the edit observed is plotted on the ordinate, while the size of each dot scales with the number of edits that meet that description. Each read is uniquely classified as a deletion, insertion, or substitution such that no one read contributes to more than one dot in this panel. In the case of substitutions, the positive ordinate reports the insertion portion of the edit, and the negative ordinate reports the deletion portion of the edit such that each edit is represented by two dots. In this panel, deletions are reported by placing a dot at the midpoint of the deleted segment. To help explain the apparent “V” pattern, dots are colored blue to indicate cases where one end of the deleted segment corresponds to the predicted cut region, presumably corresponding to a subset of the DNA repair outcomes observed. Sample dot coloring is as in **b**.

Having achieved functional delivery with our multifunctional EVs, this enabled us to next interrogate the specific contributions of each engineered EV feature. In particular, we sought to evaluate the unique contribution of the anti-CD2 scFv, since it can confer some degree of binding and uptake *in vitro*. To ascertain the requirement of EV scFv-CD2 engagement for functional delivery, we pretreated and cultured cells with an anti-CD2 antibody prior to EV addition to block receptors on recipient cells. Surprisingly, we found that pretreatment with the anti-CD2 antibody increased editing rates across vesicle types and viral glycoprotein systems (**Fig. 6a, b**). To explain this observation, we hypothesized that engagement of CD2 might result in higher levels of T cell activation, making cells more susceptible to EV uptake and editing; this would be a novel consequence of CD2 engagement if confirmed. To investigate this possibility, we incubated T cells with either EV scFvs or anti-CD2 antibodies and analyzed surface expression of CD25 2 d post-treatment. CD25 expression was minimally impacted by any treatment, indicating that T cell activation cannot explain the observed increase in editing upon CD2 engagement (**Supplementary Fig. 19**). To investigate how editing efficiency scales with practical considerations such as EV dose, and to probe how CD2 engagement may contribute to this process, we evaluated EV delivery to T cells from two distinct donors using only a single EV dose or repeating EV administration every day for the 6 d incubation (**Fig. 6c, d**). As expected, repeat EV administration increased editing efficiency in all cases, indicating that redosing is a useful handle for boosting editing. In general, scFv-CD2 engagement enhanced editing mediated by exosomes, although this effect was not evident for microvesicle-mediated editing. Finally, in order to evaluate which trends hold across experiments, we performed a combined analysis (normalizing to control for variables hypothesized to contribute to variation in editing efficiency, such as donor T cell batch-specific susceptibility to Cas9 RNP-mediated editing^57^) (**Fig. 6e, f**). Overall, these combined trends support the key conclusions noted above.

**Fig. 6:**
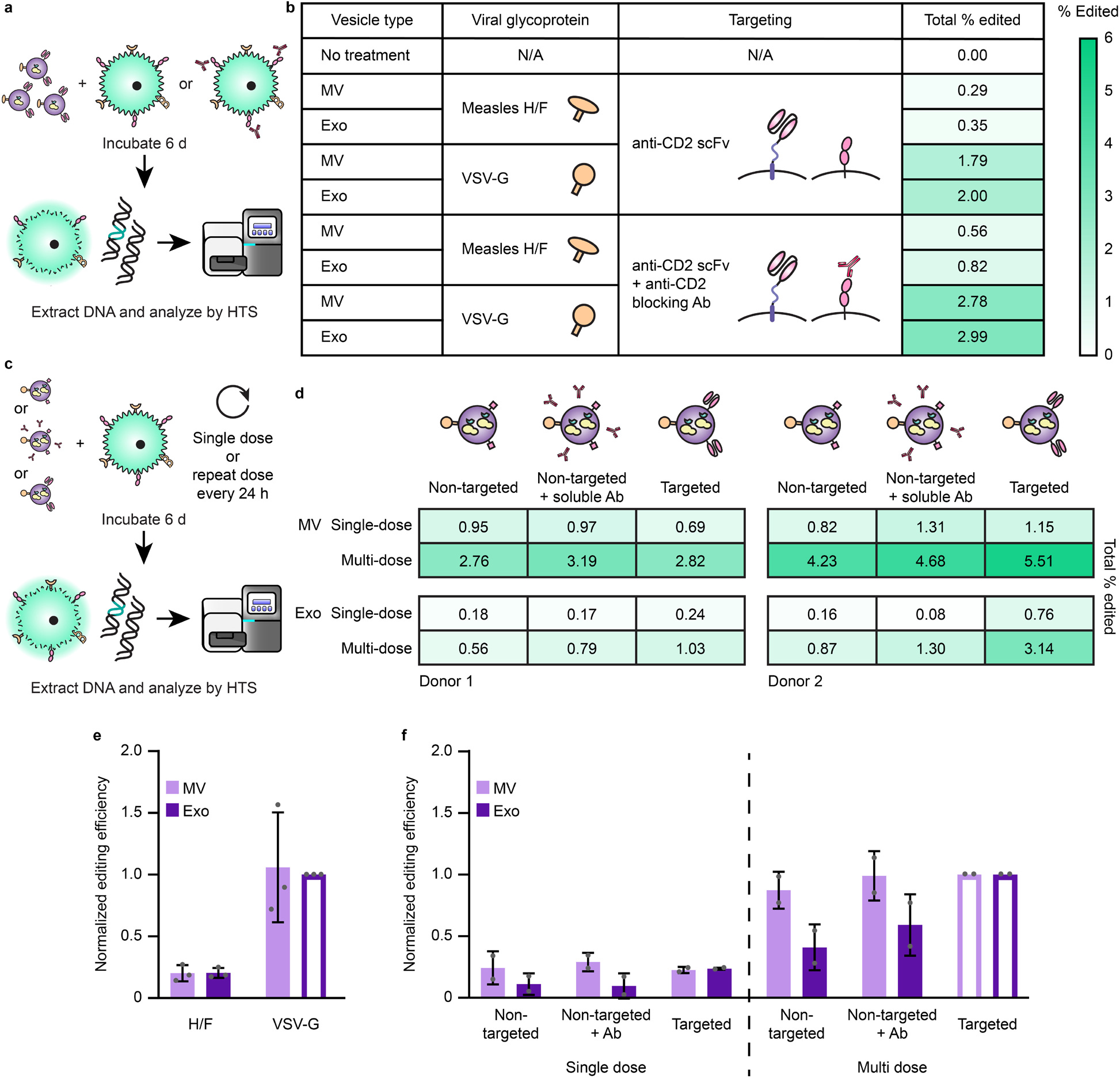
CD2 engagement and repeat dosing enhance EV-mediated functional cargo delivery and vary with vesicle subpopulation. **a**, Illustration of strategy for probing the requirement of scFv-CD2 engagement by blocking CD2. **b**, Blocking CD2 on recipient cells prior to EV addition increases total editing for all vesicle types. 8.0×10^9^ EVs were incubated per 4.0×10^4^ CD4^+^ T cells for 6 d prior to genomic DNA extraction and HTS analysis. Heat map coloring scales from 0-6% total Cas9-mediated editing. **c**,**d**, Illustration (**c**) and evaluation (**d**) of experiments probing Cas9-mediated editing after repeat EV administration and various modes of CD2 engagement. Two independent experiments using different donor cells and EV preparations are shown. EV dosing was: Donor 1—1.25×10^10^ MVs or 5.50×10^9^ exos per 5×10^4^ cells; Donor 2—1.50×10^10^ MVs or 7.50×10^9^ exos per 5×10^4^ cells. Heat map coloring is as in **b. e**, EV-mediated Cas9 functional delivery shows consistent trends across 3 donors and EV batches. Editing efficiency was normalized to the sample receiving VSV-G exosomes (open bar) for each of three independent experiments. **f**, Combined analysis of experiments presented in **d**. Within each vesicle population, editing efficiencies were normalized to the sample receiving multiple doses of VSV-G EVs (open bars); this normalization strategy is designed to control for expected sources of greatest variation (i.e., intrinsic donor/T cell batch-specific susceptibility to EVs and editing). Error bars represent one standard deviation.

## DISCUSSION

In this study, we developed the GEMINI strategy of combining genetically-encoded, general platform approaches for targeting EVs to recipient cells with surface-displayed scFvs, actively loading EVs with protein cargo via tagging with vesicle-localizing domains, and promoting uptake and fusion with recipient cells by displaying viral glycoproteins. The motivating application of achieving Cas9 delivery to T cells—a challenging objective—proved useful for refining and validating technologies that can be combined to achieve this goal.

An exciting aspect of EV-mediated delivery is the potential to target vesicles to cells and receptors of interest through engineered interactions. Prior reports have demonstrated non-targeted EV-mediated transfer to T cells, with cargo including EV-encapsulated AAV8^59^ or zinc finger-fused methyltransferases.^60^ EVs that bind T cells have also been described as a method of crosslinking T cells and other cellular targets by displaying linked anti-CD3 and anti-EGFR scFvs on the PDGFR transmembrane domain.^61^ To our knowledge, this study is the first demonstrating integration of EV targeting and uptake by T cells. We anticipate that the modularity of our targeting construct will be useful for directing EVs to other cell types and receptors.

One technology reported here involved the serendipitous discovery that the ABI domain (from the ABA dimerization system) facilitates EV cytosolic cargo protein loading even without with ABA. The mechanism of this effect is unknown. ABI is not predicted by WoLF PSORT (https://www.genscript.com/wolf-psort.html) to localize to the cell membrane or endosomal pathways. An advantage of this system is that ABI-mediated loading is easier to implement than multi-domain dimerization systems (using light,^42^ rapamycin,^43^ or Dmr domains^62^) or tags that require overexpression of helper proteins to facilitate trafficking into vesicles, such as the WW domain and Ndfip1.^63^ Other active loading tags have recently been explored by Codiak Biosciences,^64^ in this case deriving a tag from a membrane-associating protein, though the reversibility of such interactions has yet to be established. Although increased Cas9 loading did not confer additional DNA cleavage in our *in vitro* assay, potentially because this particular Cas9 fusion strategy reduced Cas9 turnover rate (**Supplementary Fig. 13c**), higher cargo loading is likely beneficial in cell delivery contexts where EVs must overcome additional barriers of uptake, fusion, cytosolic release, and intracellular trafficking. In such contexts, the advantage of delivering more Cas9-sgRNA cargo may outweigh slower reaction rates. It is also possible that the ABI fusion strategy may be refined in future work to mitigate any effects on Cas9 activity.

The eventual fate of EVs in recipient cells is often degradation in the endosomal/lysosomal pathway,^41^ and thus developing methods to achieve vesicle fusion in recipient cells is critical for achieving (or enhancing) functional cargo delivery (i.e., to the cytoplasm). Here, we demonstrated the use of VSV-G and measles virus glycoproteins H/F to achieve efficient internalization of EVs by both Jurkat and primary T cells for VSV-G and in cells expressing the lymphocyte receptor SLAM for H/F. An important translational consideration is that mutant versions of the H/F proteins have been developed to evade neutralizing host antibodies, such as those induced by the measles vaccine.^65^ However, in functional Cas9 delivery studies, we observed greater Cas9 editing efficiencies in primary T cells treated with VSV-G vesicles as compared to H/F, likely because of increased fusion of VSV-G in acidic endosomal environments.^66^ Our observed conversion efficiencies, although modest at the doses used in this exploration, meet or exceed comparable reports in the literature. Perhaps the most rigorous and compelling comparator study reported that 12 repeat, high-dose (∼1×10^11^ EVs as compared to our ∼1×10^10^ EVs) administrations of vesicles derived from MDA-MB-231 breast cancer cells loaded with an sgRNA were required to achieve conversion efficiencies on the order of 0.1% in HEK293T reporter cells that constitutively express Cas9 (a cell type to which delivery of viral vectors and various biomolecules is fairly efficient compared to T cells).^67^ We observed substantially greater conversion rates in our system, and conversion increased with repeat administration for both EV types. In the specific HIV application contemplated, conversion of even a limited pool of T cells to resist HIV infection could confer therapeutic benefits.^68^ EVs have been explored for potential utility in HIV treatment through approaches such as Cas9-mediated excision of proviruses in microglial cells,^62^ repressing viral replication with zinc finger-fused methyltransferases,^60^ or killing of infected cells using HIV Env-targeted vesicles,^69^ but these preliminary demonstrations have not yet been developed into methods for achieving specific delivery and treatment of T cells using a clinically translatable approach. Another important finding is that while exact editing efficiencies varied across donors and EV doses (a pattern observed with Cas9 RNP delivery by other methods^57^), the overall trends we observed were highly conserved when controlling for these effects, demonstrating repeatability. These results are particularly exciting when noting that the quantified efficiencies are limited by Cas9-mediated cleavage and DNA repair rates, such that we are certainly underestimating the number of functional delivery events, and other cargo types and mechanisms might confer even greater rates of functional delivery.

A surprising finding is that CD2 engagement, by either recombinant antibody or EV-displayed antibody, enhanced functional exosome-mediated delivery *in vitro*, though no comparable benefit for microvesicle-mediated delivery was observed. This combination of effects is not explained by known features of CD2/T cell biology, although it could be related to findings that ligand engagement triggers CD2 internalization^34^. While scFv-displaying vesicles of both types specifically bound CD2 and were internalized to some degree, there may exist a difference in intracellular trafficking and fusion between the two vesicle populations. For example, CD2-binding-mediated trafficking might favor fusion over native uptake pathways in a way that differentially favors exosomes. This phenomenon warrants further study to elucidate underlying mechanisms.

A key feature of this study is the selection of methods that avoid artifacts found in EV studies. One general and often overlooked artifact with EV functional delivery experiments is the risk of transfer of residual producer cell transfection reagent; particles from cells transfected with lipoplexes can mediate functional effects erroneously attributed to EVs.^24^ We minimized such risks by employing a transfection method that is unlikely to transfer plasmids to T cells. Key comparative observations (e.g., differences in functional delivery by viral glycoprotein choice) support our interpretation that we quantified true EV-mediated delivery.

The technologies employed here are generalizable and amenable to large scale production and biomanufacturing. Our strategy of genetically programming the self-assembly of multifunctional particles avoids the need for post-harvest chemical modification that necessitates further purification, lower EV yields, and may incur regulatory challenges. Although we used transient transfections for some transgenes (e.g., viral glycoproteins that cannot be constitutively expressed due to toxicity), such genes are regularly expressed from inducible promoters for production of biologics.^70,71^ We anticipate that the integrated tools developed here for EV targeting, cargo loading, and vesicle fusion will be widely applicable for a range of applications and targets, providing a flexible platform for engineering EV therapeutics.

## METHODS

### Plasmid construction

Plasmids were constructed using standard molecular biology techniques. Codon optimization was performed using the GeneArt gene synthesis tool (Thermo Fisher). PCR was performed using Phusion DNA polymerase (New England Biolabs, NEB), and plasmid assembly was performed via restriction enzyme cloning. Plasmids were transformed into TOP10 competent *E. coli* (Thermo Fisher) and grown at 37°C.

### Plasmid backbones

A modified pcDNA3.1 (Thermo Fisher V87020), was used to generate a general expression vector. Briefly, the hygromycin resistance gene and SV40 promoter were removed, leaving the SV40 origin of replication and poly(A) signal intact. The BsaI sites in the AmpR gene and 5’-UTR and the BpiI site in the bGH poly(A) signal were mutated. The lentiviral vector pGIPZ (Open Biosystems) was obtained through the Northwestern High Throughput Analysis Laboratory. plentiCRISPRv2 was a gift from Feng Zhang^72^ (Addgene plasmid No. 52961).

### Plasmid source vectors

Fluorescent proteins enhanced blue fluorescent protein 2 (EBFP2), enhanced yellow fluorescent protein (EYFP), and dimeric tomato (dTomato) were sourced from Addgene vectors (plasmid Nos. 14893, 58855, and 18917, respectively) gifted by Robert Campbell,^73^ Joshua Leonard,^74^ and Scott Sternson.^75^ dsRedExpress2 was purchased from Clontech/Takara. Monomeric teal fluorescent protein 1 (TFP1) was synthesized by Thermo Fisher. psPAX2 and pMD2.G were gifts from William Miller. The anti-CD2 scFv was synthesized from a previously published scFv sequence derived from monoclonal antibody 9.6,^33^ and the PDGFR transmembrane domain was sourced from a pDisplay system vector (Addgene plasmid No. 61556, gifted by Robert Campbell).^76^ The C1C2 domain sequence was provided by Natalie Tigue^30^ and synthesized by Thermo Fisher. Constitutively active Cx43 and SLAM were synthesized by Thermo Fisher from Uniprot sequences P17302 CXA1_HUMAN and Q13291-1 SLAF1_HUMAN isoform 1, respectively. Plasmids encoding the measles virus glycoproteins were gifts from Isabelle Clerc, Thomas Hope, and Richard D’Aquila.^50^ pX330 encoding Cas9 was gifted by Erik Sontheimer (UMass), originally sourced from Addgene plasmid No. 42230 gifted by Feng Zhang.^77^ The CXCR4 sgRNA sequence was provided by Judd Hultquist and is as follows: GAAGCGTGATGACAAAGAGG.^57^ ABI and PYL domains^45^ were codon optimized and synthesized by Thermo Fisher and IDT, respectively.

### Plasmid preparation

Bacteria were grown overnight in 100 mL LB + Amp cultures for 12-14 h. Cultures were spun at 3,000 g for 10 min to pellet the bacteria, and pellets were resuspended and incubated for 30 min in 4 mL of 25 mM Tris pH 8.0, 10 mM EDTA, 15% sucrose, and 5 mg/mL lysozyme. Bacteria were lysed for 15 min in 8 mL of 0.2 M NaOH and 1% SDS, followed by a 15 min neutralization in 5 mL of 3 M sodium acetate (pH 5.2). The precipitate was pelleted at 9,000 g for 20 min, and supernatant was filtered through cheese cloth and incubated for 1-3 h at 37°C with 3 µL of 10 mg/mL RNAse A (Thermo Fisher). Samples were extracted with 5 mL phenol chloroform, and the aqueous layer was recovered after centrifugation at 7,500 g for 20 min. A second phenol chloroform extraction was performed with 7 mL solvent. 0.7 volumes isopropanol was added to the recovered supernatant, and samples were inverted and incubated at room temperature for 10 min prior to centrifugation at 9,000 g for 20 min to pellet the DNA mixture. Pellets were briefly dried and resuspended in 1 mL of 6.5% PEG 20,000 and 0.4 M NaCl. DNA was incubated on ice overnight and pelleted at 21,000 g for 20 min. The supernatant was removed, and pellets were washed in cold absolute ethanol and dried at 37°C before resuspension in TE buffer (10mM Tris, 1 mM EDTA, pH 8.0). DNA was diluted to 1 µg/µL using a Nanodrop 2000 (Thermo Fisher).

### Cell culture

HEK293FT cells (Thermo Fisher R70007) were cultured in Dulbecco’s Modified Eagle Medium (DMEM, Gibco 31600-091) supplemented with 10% FBS (Gibco 16140-071), 1% penicillin-streptomycin (Gibco 15140-122), and 4 mM additional L-glutamine (Gibco 25030-081). Jurkat T cells (ATCC TIB-152) were cultured in Roswell Park Memorial Institute Medium (RPMI 1640, Gibco 31800-105) supplemented with 10% FBS, 1% pen-strep, and 4 mM L-glutamine. Sublines generated from these cell lines were cultured in the same way. Cells were subcultured at a 1:5 or 1:10 ratio every 2-3 d, using Trypsin-EDTA (Gibco 25300-054) to remove adherent cells from the plate. Lenti-X cells (Takara) were cultured the same way with additional 1 mM sodium pyruvate (Thermo Fisher 11360-070). Primary human CD4^+^ T cells were cultured in RPMI supplemented with 10% FBS, 1% pen-strep, 5 mM HEPES, 5 mM sodium pyruvate, and 20 U/mL IL-2 (added fresh at time of use). Cells were maintained at 37°C at 5% CO_2_. HEK293FT and Jurkat cells tested negative for mycoplasma with the MycoAlert Mycoplasma Detection Kit (Lonza LT07-318).

### Transfection

For transfection of HEK293FT cells and derived cell lines in 15 cm dishes for EV packaging, cells were plated at a density of 18×10^6^ cells/dish (1×10^6^ cells/mL) 6-12 h prior to transfection. Cells were transfected with 30 µg DNA plus 1 µg of a fluorescent transfection control via the calcium phosphate method. Plasmid DNA was mixed with 2 M CaCl_2_ (final concentration 0.3 M) and added to a 2x HEPES-buffered saline solution (280 mM NaCl, 0.5 M HEPES, 1.5 mM Na_2_HPO_4_) dropwise in a 1:1 ratio and mixed seven times by pipetting. The transfection solution w-as incubated for 3 min, mixed eight times by pipetting, and added gently to the side of the plate. For transfection of HEK293FT cells in 10 cm dishes for EV packaging, cells were plated at a density of 5×10^6^ cells/ dish (6.25×10^5^ cells/mL) and transfected with 20 µg DNA plus 1 µg transfection control as described above, adding transfection mixture dropwise to the dish. Lenti-X cells were transfected in 10 cm dishes in the same manner, though were plated 24 h prior to transfection as per the manufacturer recommendation (Takara). For transfection of HEK293FT cells in 24 well plates, cells were plated at a density of 1.7×10^5^ cells/well (3.4×10^5^ cells/mL) and transfected with 200 µg DNA as described above, adding transfection mixture dropwise to the well. Medium was changed 12-16 h later. Jurkat lipofectamine transfections were performed according to the manufacturer’s protocol.

### Cell line generation

To generate lentivirus, HEK293FT or Lenti-X cells were plated in 10 cm dishes at a density of 5×10^6^ cells/dish (6.25×10^5^ cells/mL). 6-12 h later for HEK293FT or 24 h later for Lenti-X, cells were transfected with 10 µg of viral vector, 8 µg psPAX2, and 3 µg pMD2.G via calcium phosphate transfection as described above. Medium was changed 12-16 h later. 28 h post media change, lentivirus was harvested from the conditioned medium. Medium was centrifuged at 500 g for 2 min to clear cells, and the supernatant was filtered through a 0.45 µm pore filter (VWR). Lentivirus was concentrated from the filtered supernatant by ultracentrifugation in Ultra Clear tubes (Beckman Coulter 344059) at 100,420 g at 4°C in a Beckman Coulter Optima L-80 XP ultracentrifuge using an SW41Ti rotor. Supernatant was aspirated, leaving virus in ∼100 µL final volume, and concentrated lentivirus was left on ice for at least 30 min prior to resuspension, then used to transduce ∼1×10^5^ cells, either plated at the time of transduction or the day before. When appropriate, drug selection began 2 d post transduction, using antibiotic concentrations of 1 µg/mL puromycin (Invitrogen ant-pr) and 10 µg/mL blasticidin (Alfa Aesar J61883) on HEK293FT cells or 0.2 µg/mL puromycin and 2 µg/mL blasticidin on Jurkat cells. Cells were kept in antibiotics for at least two weeks with subculturing every one to two days.

### Sorting of Cas9 reporter lines

Cells were prepared for fluorescence-activated cell sorting (FACS) by resuspending in either DMEM or RPMI, as appropriate, supplemented with 10% FBS, 25 mM HEPES, and 100 µg/mL gentamycin (Amresco 0304) at a concentration of 1×10^7^ cells/mL. Cells were sorted for the highest mTFP1 expressors (top 10% or less) lacking any dTomato expression on a BD FacsAria Ilu using a 488 nm laser (530/30 filter) and a 562 nm laser (582/15 filter). Cells were collected in DMEM or RPMI, as appropriate, supplemented with 20% FBS, 25 mM HEPES, and 100 µg/mL gentamycin. Cells were spun down and resuspended in normal growth medium with 100 µg/mL gentamycin for recovery.

### EV production, isolation, and characterization

EV producer cell lines were plated in 10 or 15 cm dishes and transfected the same day by the calcium phosphate method where appropriate. Medium was changed to EV-depleted medium the following morning. EV-depleted medium was made by supplementing DMEM with 10% exosome depleted FBS (Gibco A27208-01), 1% pen-strep, and 4 mM L-glutamine. EVs were harvested from the conditioned medium 24-36 h post medium change by differential centrifugation as previously described.^36,37^ Briefly, conditioned medium was cleared of debris by centrifugation at 300 g for 10 min to remove cells followed by centrifugation at 2,000 g for 20 min to remove dead cells and apoptotic bodies. Supernatant was centrifuged at 15,000 g for 30 min in a Beckman Coulter Avanti J-26XP centrifuge with a J-LITE JLA 16.25 rotor to pellet microvesicles. Supernatant was collected and exosomes pelleted by ultracentrifugation at 120,416 g for 135 min in a Beckman Coulter Optima L-80 XP ultracentrifuge with an SW41 Ti rotor, using polypropylene ultracentrifuge tubes (Beckman Coulter 331372). All centrifugation steps were performed at 4°C. EV pellets were left in ∼100-200 µL of conditioned media and incubated on ice for at least 30 min after supernatant removal before resuspension. EV concentration was determined by NanoSight analysis. Samples were diluted in PBS to concentrations on the order of 10^8^ particles/mL for analysis. NanoSight analysis was performed on an NS300 (Malvern), software version 3.4. Three 30 s videos were acquired per sample using a 642 nm laser on a camera level of 14, an infusion rate of 30, and a detection threshold of 7. Default settings were used for the blur, minimum track length, and minimum expected particle size. EV concentrations were defined as the mean of the concentrations calculated from each video. Size distributions were generated by the software. For TEM, samples were fixed for 10 min in Eppendorf tubes by adding 65 µL of 4% PFA to 200uL of EVs. 15 µL of fixed suspension was pipetted onto a plasma cleaned (PELCO easiGlow), formvar/carbon coated grid (EMS 300 mesh). After 10 min, the solution was removed by wicking with a wedge of filter paper, then washed by inverting the grid onto a drop of buffer for 30 seconds twice, followed with diH_2_O once. A 2% uranyl acetate (Ted Pella) stain was applied twice and wicked after 30 s. Grids were allowed dry before storing in a grid box until use. Grids were imaged in a JEOL JEM 1230 TEM (JEOL USA) with a 100 KV accelerating voltage. Data was acquired with a Orius SC1000 CCD camera (Gatan). EVs were stored on ice and used within 10 days or stored at -80°C for long term preservation.

### Immunoblotting

For western blot analysis, cells were lysed in RIPA buffer (150 mM NaCl, 50 mM Tris-HCl pH 8.0, 1% Triton X-100, 05% sodium deoxycholate, 0.1% SDS, and one protease inhibitor cocktail tablet (Pierce PIA32953) per 10 mL) and incubated on ice for 30 min. Lysates were cleared by centrifugation at 12,000 g for 20 min at 4°C, and supernatant was harvested. Protein concentration was determined by BCA assay (Pierce) according to the manufacturer’s instructions. Samples were normalized by protein content ranging from 1 to 2 µg (for cell lysates) or by vesicle count ranging from 1×10^7^ to 6×10^8^ (for EVs). Samples were heated in Laemmli buffer (60 mM Tris-HCl pH 6.8, 10% glycerol, 2% SDS, 100 mM dithiothreitol, 0.01% bromophenol blue) at 70°C (for membrane-bound scFv and calnexin) or 98°C (for Cas9, CD9, CD81, and Alix) for 10 min. Samples were loaded onto 4-15% polyacrylamide gradient Mini-PROTEAN TGX precast protein gels (Bio-Rad) and run at 50 V for 10 min followed by 100 V for 1 h. Protein was transferred to a PVDF membrane (Bio-Rad) at 100 V for 45 min. For anti-FLAG blots, membranes were blocked in 3% milk in TBS (50 mM Tris, 138 mM NaCl, 2.7 mM KCl, pH 8.0) for 30 min. Membranes were washed once in TBS for 5 min, then incubated in primary anti-FLAG antibody (Sigma F1804) diluted 1:1000 in 3% milk in TBS overnight at 4°C. Membranes were washed once for 5 min in TBS and twice in TBST 1 (50 mM Tris, 138 mM NaCl, 2.7 mM KCl, 0.05% Tween 20, pH 8.0) for 5 min each prior to secondary antibody staining. For all other blots, membranes were blocked in 5% milk in TBST 2 (50 mM Tris, 150 mM NaCl, 0.1% Tween 20, pH 7.6) for 1 h. Membranes were incubated in primary antibody diluted in 5% milk in TBST 2 overnight at 4°C. Primary antibodies include anti-HA (Cell Signaling Technology 377245 C29F4, 1:1000), anti-CD9 (Santa Cruz Biotechnology sc-13118, 1:500), anti-CD81 (Santa Cruz Biotechnology sc-23962, 1:500, run in non-reducing conditions), anti-Alix (Abcam Ab117600, 1:500), and anti-calnexin (Abcam Ab22595, 1:1000). Membranes were washed three times in TBST 2 for 5 min each prior to secondary antibody staining. HRP-conjugated anti-mouse (Cell Signaling Technology 7076) and anti-rabbit (Invitrogen 32460) secondary antibodies were diluted 1:3000 in 5% milk in TBST 2. Membranes were incubated in secondary antibody at room temperature for 1 h, then washed three times in TBST 2 (5 min washes). Membranes were probed with Clarity Western ECL Substrate (Bio-Rad) and either exposed to film, which was developed and scanned, or imaged using an Azure c280 imager. Images were cropped using Adobe Illustrator. No other image processing was employed.

### Surface immunoblotting

Cells were transferred to FACS tubes (adherent cells were harvested using FACS buffer (PBS pH 7.4 with 0.05% BSA and 2 mM EDTA) prior to staining) with 1 mL of FACS buffer and centrifuged at 150 g for 5 min. Supernatant was decanted, and cells were resuspended in 50 µL of FACS buffer. 10 µL of human IgG (Thermo Fisher 027102) was added, cells were flicked to mix, and were incubated at 4°C for 5 min. Conjugated primary antibody was then added at the manufacturer’s recommended dilution, cells were flicked to mix and incubated at 4°C for 30 min. Cells were then washed three times with 1 mL of FACS buffer, centrifuging at 150 g for 5 min and decanting the supernatant after each wash. Cells were resuspended in two drops of FACS buffer prior to flow cytometry. For Miltenyi Biotec antibodies, cells were stained at 4°C for 15 min without blocking and were washed once prior to flow cytometry, as per manufacturer protocol. Antibodies used in this study were as follows: Anti-FLAG-APC (Abcam ab72569), anti-CD2-APC (R&D Systems FAB18561A), anti-CD25-PE (Miltenyi REA945, 130-115-628), anti-SLAM-PE (Miltenyi REA151, 130-123-970), and anti-mouse IgG1-APC (R&D Systems IC002A) or anti-human IgG1-PE (Miltenyi REA293, 130-113-438) were used as isotype controls where appropriate.

### EV binding and uptake experiments

Jurkat T cells or primary human CD4^+^ T cells were incubated with EVs at an EV to cell ratio of 100,000:1 (typically 1×10^10^ EVs per 1×10^5^ cells) unless otherwise indicated. For Jurkats, cells were plated in a 48 well plate with 300 µL total volume. For primary T cells, cells were plated in a 96 well plate with 200 µL total volume. Cells were plated at the time of EV addition, and wells were brought to the appropriate volume with RPMI. For binding experiments, cells were incubated for 2 h at 37°C unless otherwise indicated, then washed three times in FACS buffer, centrifuging at 150 g for 5 min for Jurkat cells or 400 g for 3 min for primary T cells. Cells were resuspended in one drop of FACS buffer prior to flow cytometry. To adsorb EVs to aldehyde/sulfate latex beads (Thermo Fisher), EVs were mixed with beads at a ratio of 1×10^9^ EVs per 2 µL beads diluted 1:10 in PBS. Volumes were normalized across samples with PBS, and beads and EVs were incubated for 15 min at room temperature. Samples were then brought to 200 µL with PBS and allowed to incubate for 2 h at room temperature while rocking. Cells were blocked with an anti-CD2 antibody binding the same epitope as the scFv (Beckman Coulter A60794) or with blank EVs for 1 h at 37°C prior or EV incubation where indicated. For EV uptake experiments with viral glycoproteins, cells were incubated with EVs for 16 h at 37°C. To prepare for analysis, cells were washed twice in PBS and incubated with two drops of trypsin-EDTA for 5 min at 37°C to remove surface bound vesicles. Cells were washed with RPMI to quench the trypsin, then washed twice more with FACS buffer prior to analysis.

### Analytical flow cytometry and analysis

Flow cytometry was performed on a BD LSR Fortessa Special Order Research Product using the 562 nm laser for dTomato (582/15 filter), the 488 nm laser for EYFP (530/30 filter), and the 488 nm and 405 nm lasers for mTFP1 (530/30 filter and 525/50 filter, respectively). Approximately 10,000 live cells were collected per sample for analysis. Data were analyzed using FlowJo v10 (FlowJo, LLC). Briefly, cells were identified using an FSC-A vs SSC-A plot and gated for singlets using an FSC-A vs FSC-H plot (**Supplementary Fig. 20**). Fluorescence data were compensated for spectral bleed-through where appropriate. Mean fluorescence intensity (MFI) of single-cell samples was exported and averaged across three biological replicates. Autofluorescence from untreated cells was subtracted from other samples. Standard error of the mean was propagated through calculations. Where indicated, 9 peak Ultra Rainbow Calibration Particles (Spherotech URCP-100-2H) were used to generate a calibration curve to convert fluorescence into absolute fluorescence units.

### Confocal microscopy

Cells were transfected via the calcium phosphate method on poly L-lysine coated glass coverslips and mounted on glass slides for imaging. Microscopy images were taken on Leica SP5 II laser scanning confocal microscope using a 100x oil-immersion objective. Bright-field images were acquired at a PMT setting of 443.0 V. A 514 nm laser at 20% intensity and 94% smart gain was used for fluorescence excitation. Emission spectra were captured from 520-540 nm using an HyD sensor. Images were captured at 512 × 512 resolution at scanning speed of 400 Hz. Pseudocolored fluorescence images were contrast-adjusted in ImageJ such that 4% of pixels were saturated.

### Affinity chromatography

Affinity chromatography isolation was performed as previously reported.^46^ Briefly, an anti-FLAG affinity column was prepared by loading anti-FLAG M2 affinity gel (Sigma A2220-1ML) in a 4 mL 1 × 5 cm glass column (Bio-Rad) and drained via gravity flow. The column was washed with 5 mL TBS (50 mM Tris-HCl, 150 mM NaCl, pH 7.5) and equilibrated with three sequential 1 mL washes with regeneration buffer (0.1 M glycine-HCl, pH 3.5), followed by a 5 mL wash of TBS. Concentrated EVs were loaded onto the top of the column and chased with 1-2 mL of TBS. The column was incubated with EVs for 5 min before continuing. The flow through was then re-loaded onto the column such that the EV-containing medium passed through the matrix five times. The column was washed with 10 mL TBS prior to elution. EVs were eluted with 2.5 mL elution buffer (100 µg/mL 3x FLAG peptide (Sigma F4799-4MG) in TBS), which was incubated on the column for 5-10 min after the void fraction was drained (∼1 mL). Five fractions of EVs were collected in 0.5 mL fractions (approximately 8 drops off the column per fraction). The column was regenerated with three sequential 1 mL washes with regeneration buffer and stored at 4°C in storage buffer (50% glycerol, 0.02% sodium azide in TBS).

### Cas9 in vitro cleavage assays

EVs were produced as described above with components transiently transfected in 10 cm dishes with the following DNA ratios: 6 µg anti-CD2 scFv, 9 µg Cas9 vector, 5 µg sgRNA vector, and 1 µg mTFP1 transfection control. EVs were lysed by incubation with mammalian protein extraction reagent (MPER, Thermo Fisher) for 10 min at room temperature (20-23°C) with gentle agitation. 200 ng of linearized target plasmid template was added to vesicles with Cas9 reagent buffer (IDT, Alt-R CRISPR-Cas9 System), and samples were incubated at 37°C for 1 h. Proteinase K (Thermo Fisher) was added to samples at 1 µL per 10 µL of reaction mixture and incubated at 55°C for 10 min. Samples were run on a 1% agarose gel stained with SYBR safe (Thermo Fisher) and imaged using a BioDoc-It imaging system (VWR).

### Primary CD4^+^ T cell isolation, culture, and activation

Peripheral blood mononuclear cells (PBMCs) were isolated by density gradient centrifugation using Ficoll-Paque Plus (GE Health Care, #17-1440-02). PBMCs were washed with PBS three times to remove platelets and suspended at a final concentration of 5×10^8^ cells/mL in PBS, 0.5% BSA, 2 mM EDTA. Bulk CD4^+^

T cells were subsequently isolated from PBMCs by magnetic negative selection using an EasySep Human CD4^+^ T Cell Isolation Kit (STEMCELL, per manufacturer’s instructions). Isolated CD4^+^ T cells were suspended in RPMI-1640 (Gibco) supplemented with 5 mM 4-(2-hydroxyethyl)-1-piperazineethanesulfonic acid (HEPES, Corning), 50 μg/mL penicillin/streptomycin (P/S, Corning), 5 mM sodium pyruvate (Corning), and 10% FBS (Gibco). Media was supplemented with 20 IU/mL IL-2 (Miltenyi) immediately before use. For activation, bulk CD4^+^ T cells were immediately plated on anti-CD3 coated plates [coated for 12 h at 4°C with 20 µg/mL anti-CD3 (UCHT1, Tonbo Biosciences)] in the presence of 5 µg/mL soluble anti-CD28 (CD28.2, Tonbo Biosciences). Cells were stimulated for 72 h at 37°C and 5% CO_2_. After stimulation, cell purity and activation were verified by CD4/CD25 immunostaining and flow cytometry as previously described.^57^

### EV functional delivery experiments

EVs were produced as described above with components transiently transfected in 10 cm dishes with the following DNA ratios: 6 µg anti-CD2 scFv, 9 µg dual Cas9 and sgRNA vector, either 2.5 µg each of measles virus glycoproteins H/ F or 3 µg VSV-G with 2 µg filler promoterless pcDNA, and 1 µg mTFP1 transfection control. For generation of vesicles lacking the scFv, a PDGFR-bound 3x FLAG tag construct in the same vector backbone was transfected at the same plasmid copy number in place of the scFv. EVs were delivered to primary human CD4^+^ T cells as described above. Cells were cultured in the presence of EVs for 6 days, adding fresh supplemental RPMI and IL-2 every 2-3 days. For repeat dose administration, 100 µL of media were carefully removed from the top of each well and replaced with 100 µL fresh EVs and media. Cells were harvested on day 6 and washed with PBS by centrifugation at 400 g for 3 min a 4°C to pellet. Cells were resuspended in 100 µL QuickExtract DNA Extract Solution (Lucigen QE9050), and genomic DNA was harvested according to the manufacturer’s protocol. Briefly, samples were vortexed for 15 s, heated at 65°C for 6 min, vortexed for 15 s, and heated at 98°C for 2 min. DNA was stored at -80°C.

### High Throughput Sequencing (HTS) library generation

Approximately 100 ng genomic DNA was used as a template in the first round PCR amplification. The CXCR4 region of interest was amplified with the following primers: F1: 5’ ACA CTC TTT CCC TAC ACG CTC TTC CGA TCT NNN NNG AGA AGC ATG ACG GAC AAG TAC AG 3’ R1: 5’ GTG ACT GGA GTT CAG ACG TGT GCT CTT CCG ATC TNN NNN TCC CAA AGT ACC AGT TTG CCA C 3’ The PCR protocol was as follows: 98°C 3 min, (98°C 15 s, 65°C 30 s, 72°C 3 s) x 15, 72°C 5 min, 4°C 5 min. PCR products were purified using MagJET beads (Thermo Fisher K2821) and used as templates in a second round PCR amplification with the following primers: F2: 5’ AAT GAT ACG GCG ACC GAG ATC TAC ACT CTT TCC CTA CAC GAC GCT CTT CCG ATC T 3’ R2: 5’ CAA GCA GAA GAC GGC ATA CGA GAT-Index-GTG ACT GGA GTT CAG ACG TGT GCT C 3’ The PCR cycles were as follows: 98°C 3 min, (98°C 15 s, 69°C 30 s, 72°C 5 s) x 20, 72°C 5 min, 4°C 5 min. PCR products were again purified using MagJET beads prior to HTS. Both first and second round PCRs were run with primer concentrations of 200 nM and Phusion DNA polymerase.

### HTS

Genomic DNA sample concentrations were measured on a Qubit using an HS dsDNA kit and pooled in libraries with equimolar concentrations. Libraries were diluted to 4 nM in serial dilutions. Libraries and PhiX were denatured with NaOH according to the Illumina MiSeq guide and diluted to 14 pM. Reaction mixtures consisted of 8% PhiX and 92% library. Samples were run on an Illumina MiSeq using a MiSeq Reagent Kit v3, collecting paired-end reads. Data were analyzed using custom code developed by 496code (see **Data and code availability**). The overall strategy for analyzing these data is summarized in **Supplementary Note 1**.

### Statistical analysis

Statistical details are described in the figure legends. Unless otherwise stated, three independent biological replicates (cells) or technical replicates (beads) were analyzed per condition, and the mean fluorescence intensity of approximately 10,000 live single cells or beads were analyzed per sample. Unless otherwise indicated, error bars represent the standard error of the mean. Pairwise comparisons were made using two-tailed Student’s t-tests in Excel with the null hypothesis that the two samples were equal. The significance threshold was set to 0.05. Tests were followed by a Benjamini-Hochberg procedure applied within each panel of a given figure to decrease the false discovery rate.

### Reporting summary

Further information on research design is available in the Nature Research Reporting Summary linked to this article.

## Data and code availability

All reported experimental data are included as **Source Data**. The raw datasets generated during and/or analyzed during the current study are available from the corresponding author on reasonable request. Plasmid maps for all plasmids reported in this study are provided as annotated GenBank files in **Source Data**. Key plasmids used in this study are deposited with and distributed by Addgene, including complete and annotated GenBank files, at https://www.addgene.org/Joshua_Leonard/. Code for analyzing HTS data will be provided at https://github.com/leonardlab/GEMINI-HTS under an open-source license along with the final published version of this manuscript.

## Acknowledgements

We thank Dr. Richard D’Aquila for his support and guidance in starting this project. We thank Dr. Isabelle Clerc for her assistance with the measles virus glycoproteins. This work was supported by the Third Coast Center for AIDS Research (CFAR), an NIH funded center (P30 AI117943), NIH grants R01AI165236 and R01AI150998 (J.F.H.), National Science Foundation Award 1844219 (J.N.L. and Neha P. Kamat), and Kairos Ventures (gift). This work was also supported by NSF Graduate Research Fellowship (NSF GRFP) award DGE-1324585 (to D.M.S.). Sanger sequencing was performed through the NUSeq Core Facility of Northwestern’s Center for Genetic Medicine and a partnership with ACGT, Inc. NanoSight analysis was performed in the Analytical bioNanoTechnology Core Facility of the Simpson Querrey Institute at Northwestern University. ANTEC is currently supported by the Soft and Hybrid Nanotechnology Experimental (SHyNE) Resource (NSFECCS-1542205). We thank Charlene Wilke for her assistance with TEM. TEM was performed at the BioCryo facility of Northwestern University’s NU*ANCE* Center, which has received support from the Soft and Hybrid Nanotechnology Experimental (SHyNE) Resource (NSF ECCS-1542205); the MRSEC program (NSF DMR-1720139) at the Materials Research Center; the International Institute for Nanotechnology (IIN); and the State of Illinois, through the IIN. It also made use of the CryoCluster equipment, which has received support from the MRI program (NSF DMR-1229693). We thank Hailey Edelstein for her assistance with confocal microscopy. Microscopy was performed at as performed at the Biological Imaging Facility at Northwestern University (RRID:SCR_017767), graciously supported by the Chemistry for Life Processes Institute, the NU Office for Research, and the Department of Molecular Biosciences. We thank Paul Mehl for his assistance with FACS. Flow cytometry was performed at the Northwestern University RHLCCC Flow Cytometry Facility, which is supported by a Cancer Center Support Grant (NCI CA060553). We thank Jim Brink and Steve Hockema at 496code for their assistance with HTS data analysis.

## Author Contributions

D.M.S. and J.N.L. conceptualized the project and designed the experiments. D.M.S. performed the experiments. D.M.S. and J.N.L. analyzed the data. L.M.S. isolated and activated the primary T cells. K.E.B. and L.C. conducted the MiSeq runs. D.M.S. drafted the original manuscript and created the figures. J.N.L., J.F.H., and J.B.L. supervised the work. All authors reviewed, edited, and approved the final manuscript.

## ADDITIONAL INFORMATION

**Supplementary information** is available for this paper online.

**Correspondence** and requests for materials should be addressed to J.N.L.

