## Supplementary Information for "Bioengineering multifunctional extracellular vesicles for targeted delivery of biologics to T cells"

**for**

Bioengineering multifunctional extracellular vesicles

for targeted delivery of biologics to T cells

Devin M. Stranford[1,2], Lacy M. Simons[3,4], Katherine E. Berman[2,5], Luyi Cheng[2,5], Julius B. Lucks[1,2,5,6], Judd F. Hultquist[3,4], Joshua N. Leonard[1,2,5,6,7]*

1. Department of Chemical and Biological Engineering, Northwestern University, Evanston, IL, 60208, USA
2. Center for Synthetic Biology, Northwestern University, Evanston, IL, 60208, USA
3. Department of Medicine, Division of Infectious Diseases, Northwestern University Feinberg School of Medicine, Chicago, IL, 60611, USA
4. Center for Pathogen Genomics and Microbial Evolution, Northwestern University Havey Institute for Global Health, Chicago, IL, 60611, USA
5. Interdisciplinary Biological Sciences Training Program, Northwestern University, Evanston, IL, 60208, USA
6. Chemistry of Life Processes Institute, Northwestern University, Evanston, IL, 60208, USA
7. Member, Robert H. Lurie Comprehensive Cancer Center, Northwestern University, Evanston, IL, 60208, USA

Contents

**Supplementary Figures 1-20**

**Supplementary Note 1: Summary of HTS analysis code workflow**

**Supplementary Table 1: Statistical test p-values**

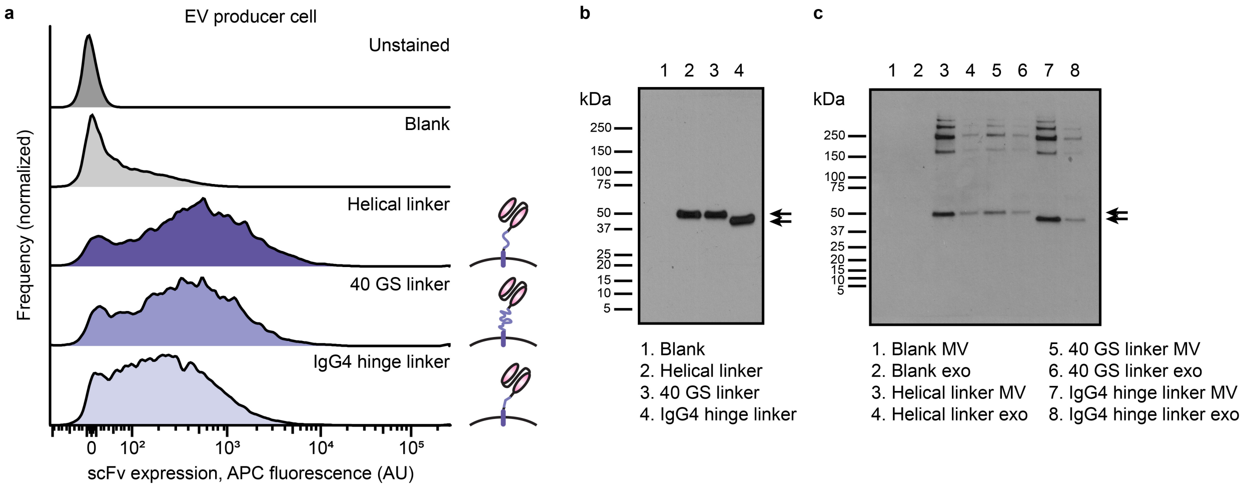


**Supplementary Fig. 1: EVs harvested from anti-CD2 scFv-expressing cell lines contain full length scFvs. a,** Surface stain (via 3x FLAG tag) of stable cell lines expressing anti-CD2 scFvs with different linkers between the binding and transmembrane domains. **b, c,** Expression of scFvs in stable HEK293FT producer cell lines (**b**) or EVs harvested from those cell lines (**c**). Expected band size: ~38-40 kDa (arrows). 0.5 µg cell lysate or 1.0x$10^8$ EVs were loaded per lane.

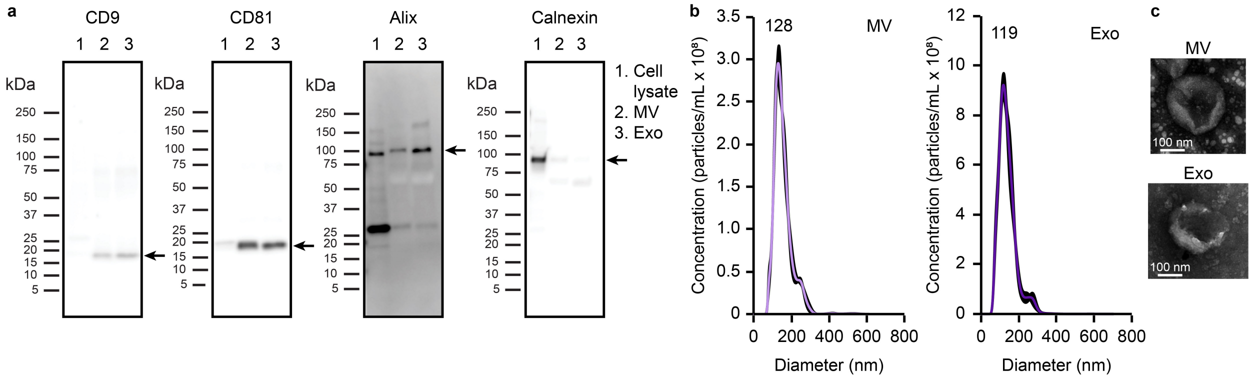


**Supplementary Fig. 2: EVs harvested via differential ultracentrifugation display characteristic surface markers, size distribution, and morphology. a,** Detection of CD9 (25 kDa), CD81 (26 kDa), and Alix (96 kDa) in both microvesicle (MV) and exosome (Exo) EV fractions. EV fractions contained minimal calnexin (~90 kDa). Expected band positions are indicated by arrows. 3 µg cell lysate or $4.5x10^8$ vesicles were loaded per lane. **b,** Representative NTA size distributions of EV subpopulations. Numbers above histograms refer to the mode size. Error bars (black) indicate standard error of the mean, calculated for each bin. **c,** Representative TEM of EV subpopulations.

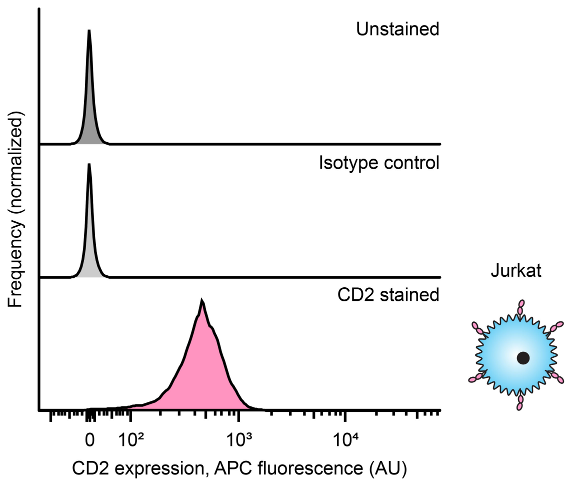


**Supplementary Fig. 3: Jurkat T cells express CD2 on the cell surface.** Cells were surface stained for CD2 expression and analyzed by flow cytometry.

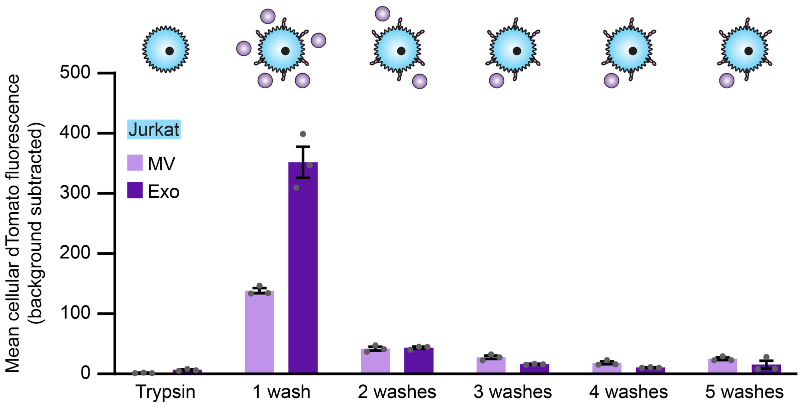


**Supplementary Fig. 4: Repeat washing removes non-specifically bound EVs from Jurkats.** EVs loaded with dTomato were incubated with Jurkat T cells for 2 h at 37°C and subjected to different numbers of washes to remove excess vesicles prior to analysis by flow cytometry. Cells treated with trypsin for 5 min after EV incubation were used as a reference (a proxy for complete EV removal from the cell surface). 3 washes were used in all subsequent binding assays.

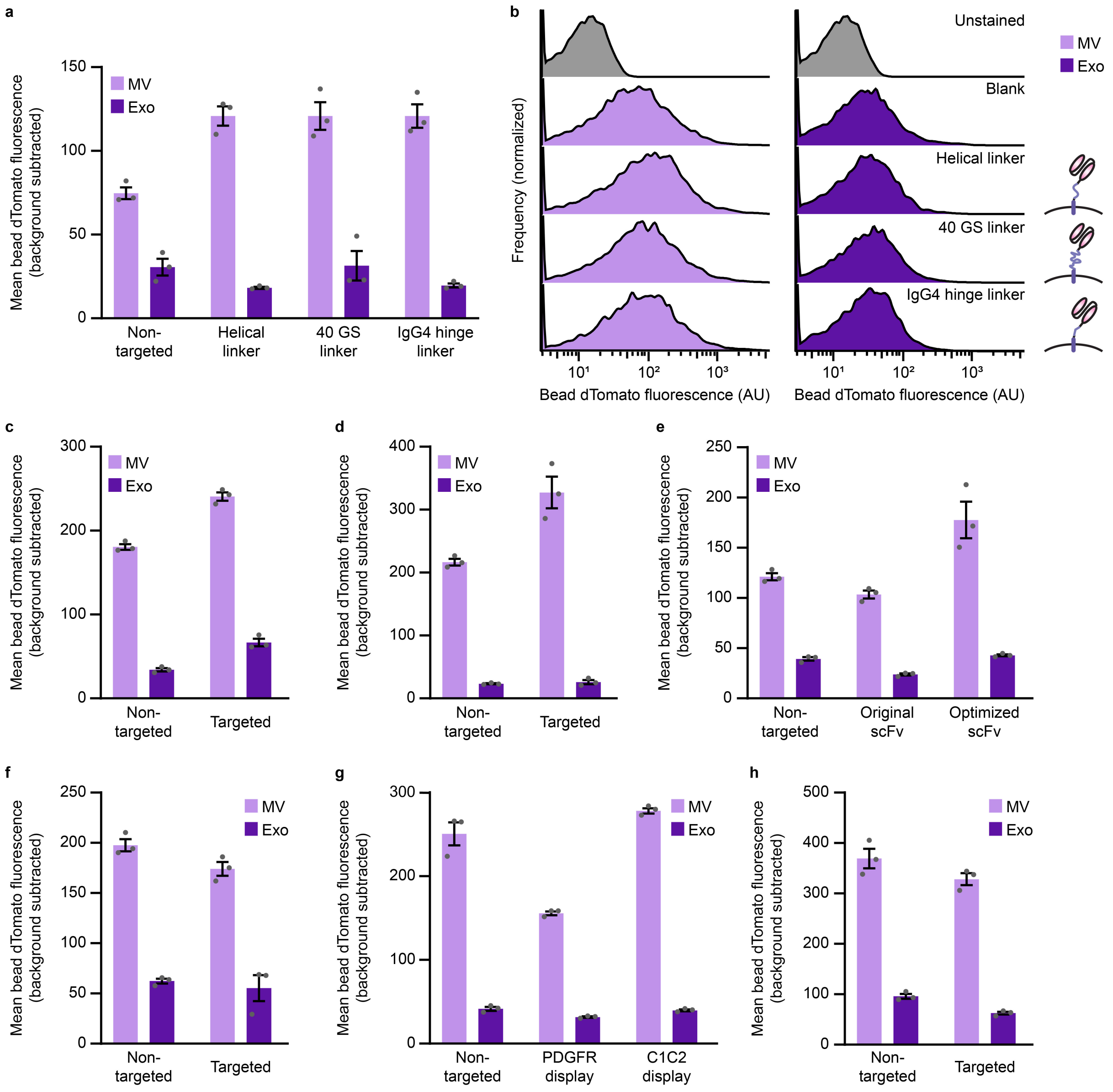


**Supplementary Fig. 5: EVs harvested from fluorescent cells have similar mean fluorescence within subsets. a,** EV dTomato loading evaluations for **Fig. 2b**. EVs were adsorbed to aldehyde/sulfate latex beads and analyzed by flow cytometry to determine a bulk population fluorescence. Experiments were performed in biological triplicate, and error bars indicate standard error of the mean. **b,** Representative histograms of EV-loaded bead fluorescence distributions represented in **a**. **c-h,** EV dTomato loading evaluations for **Fig. 2d** and **Supplementary Fig. 6** (**c**), **Fig. 2e** (**d**), **Fig. 2f** (**e**), **Figs. 2g** and **2h** (**f**), **Supplementary Fig. 9d** (**g**), and **Supplementary Fig. 9f** (**h**). In general, some differences in mean bead fluorescence were observed across samples of a given EV subtype, but these variations do not explain trends observed in the related cell delivery experiments.

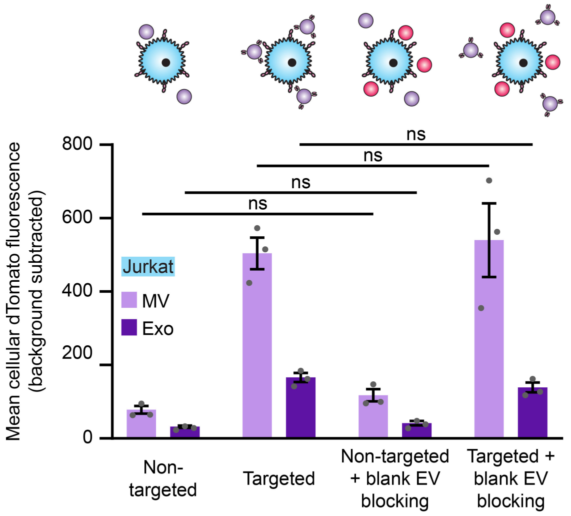


**Supplementary Fig. 6: Blocking EV recipient cells with blank EVs does not impact targeted or background binding.** Recipient Jurkat T cells were incubated for 1 h in the presence or absence of non-fluorescent, non-targeted EVs (red) to block scavenger receptors prior to a 2 h incubation with fluorescent, targeted vesicles (purple). Experiments were performed in biological triplicate, and error bars indicate standard error of the mean. Statistical tests comprise two-tailed Student's t-tests using the Benjamini-Hochberg method to reduce the false discovery rate. (*$p < 0.05$, **$p < 0.01$, ***$p < 0.001$). Exact p-values are reported in **Supplementary Table 1**. EV dTomato loading evaluations are presented in **Supplementary Fig. 5c**.

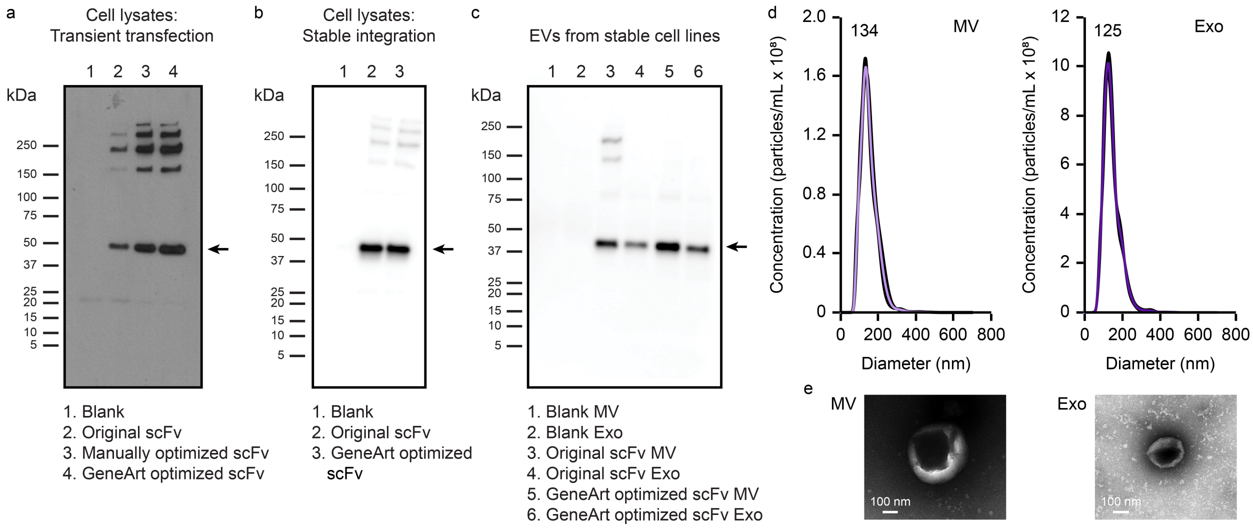


**Supplementary Fig. 7: Codon optimization increases scFv display on EVs without altering EV morphology. a,** Expression of scFv constructs in EV producer cell lysates with different levels of codon optimization. The low-expressing (non-optimized) construct was used in previous experiments, the medium-expressing construct was generated through manual codon optimization, and the high-expressing construct was optimized through Fisher GeneArt synthesis. The low- and high- expressing constructs were carried forward for further evaluation. 0.2 µg cell lysate was loaded per lane. Expected band size: ~40 kDa (arrows). A 3x FLAG tagged NanoLuc (~20 kDa) was used as a transfection control to evaluate whether possible differences in transfection efficiency may account for observed differences in protein expression. **b,** Expression of original and optimized scFv constructs in stable EV producer cell lines. 2 µg cell lysate was loaded per lane. **c,** scFv display in EVs harvested from cells in **b**. $4.5 \times 10^8$ EVs were loaded per lane. **d,** Representative NTA size distributions of optimized scFv-displaying EV subpopulations. Numbers above histograms refer to the mode size. Error bars (black) indicate standard error of the mean, calculated for each bin. **e,** Representative TEM of optimized scFv-displaying EVs.

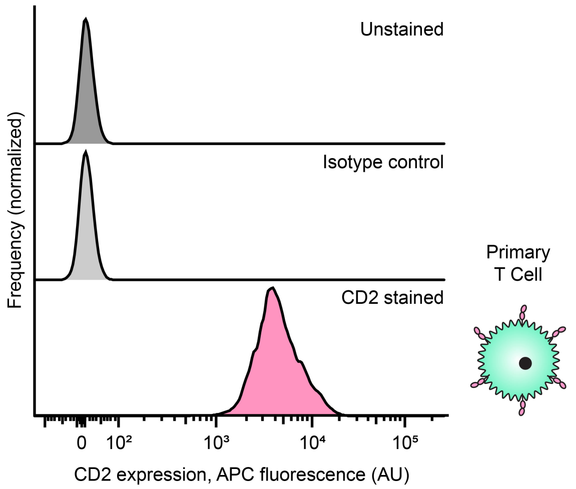


**Supplementary Fig. 8: Primary T cells express CD2 on the cell surface.** Cells were surface stained for CD2 expression and analyzed by flow cytometry.

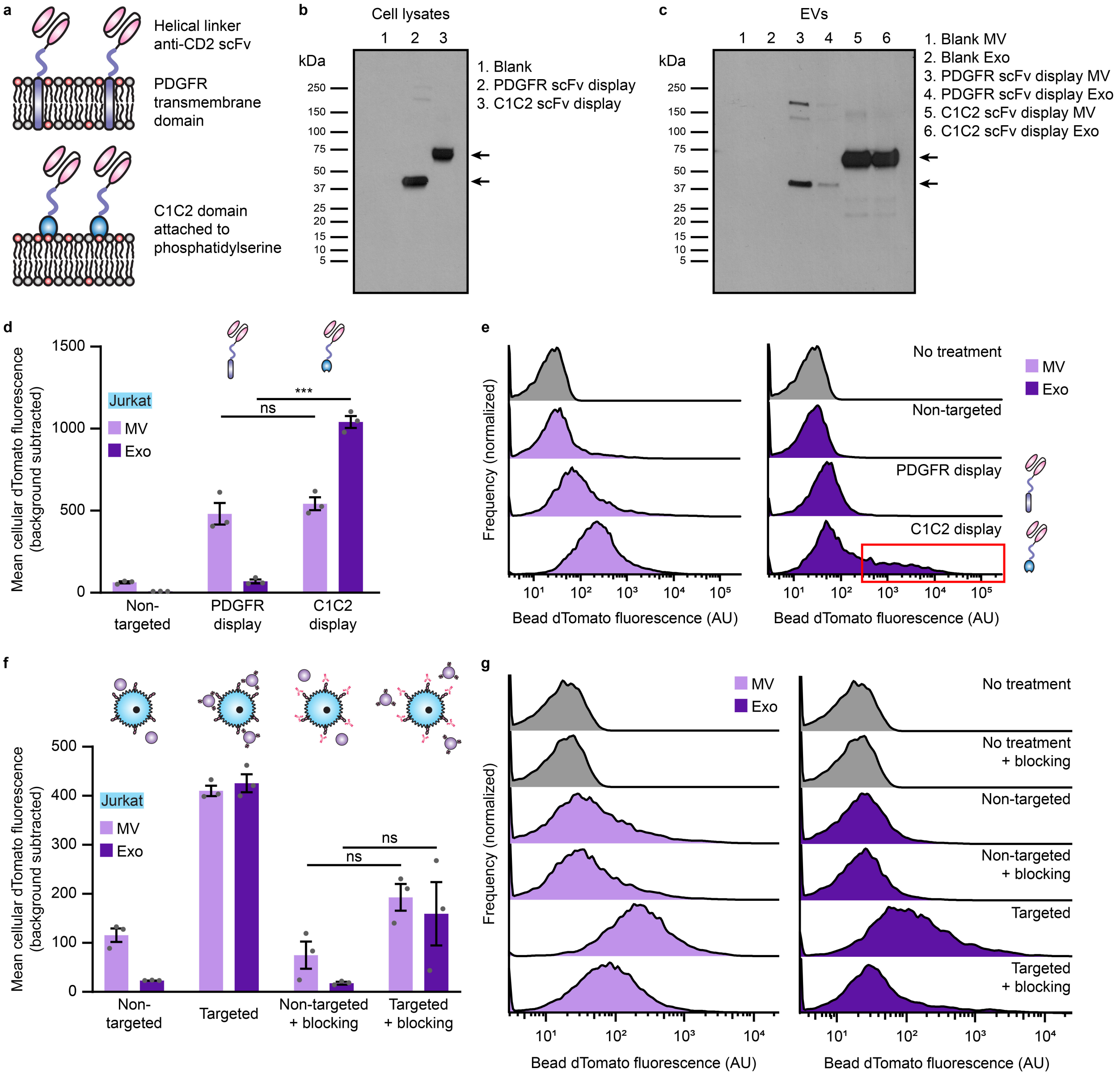


**Supplementary Fig. 9: Different scFv display techniques result in different EV targeting properties. a,** Cartoon highlighting the structures of the PDGFR transmembrane domain scFv display and lactadherin C1C2 domain anchoring to phosphatidylserine. **b,** Expression of scFv constructs in EV producer cell lysates. 1 µg cell lysate was loaded per lane. Expected band sizes: ~40 kDa and ~75 kDa (black arrows). **c,** Loading of scFv constructs into EVs generated from cell lines in **b**. $5.0 \times 10^8$ EVs were loaded per lane. **d,** Binding of targeted EVs to Jurkat T cells following a 2 h incubation. **e,** Representative histograms corresponding to the summary data reported in **d**. The subpopulation of cells showing a skewed, high degree of exosome binding is indicated by the red box. **f,** Recipient Jurkat T cells were incubated for 1 h in the presence or absence of anti-CD2 antibodies prior to a 2 h incubation with EVs. **g,** Representative histograms corresponding to the summary data reported in **f**. Flow cytometry experiments were performed in biological triplicate, and error bars (panels **d**, **f**) indicate standard error of the mean. EV dTomato loading evaluations are presented in **Supplementary Fig. 5**. Statistical tests comprise two-tailed Student's t-tests using the Benjamini-Hochberg method to reduce the false discovery rate. (*$p < 0.05$, **$p < 0.01$, ***$p < 0.001$). Exact p-values are reported in **Supplementary Table 1**.

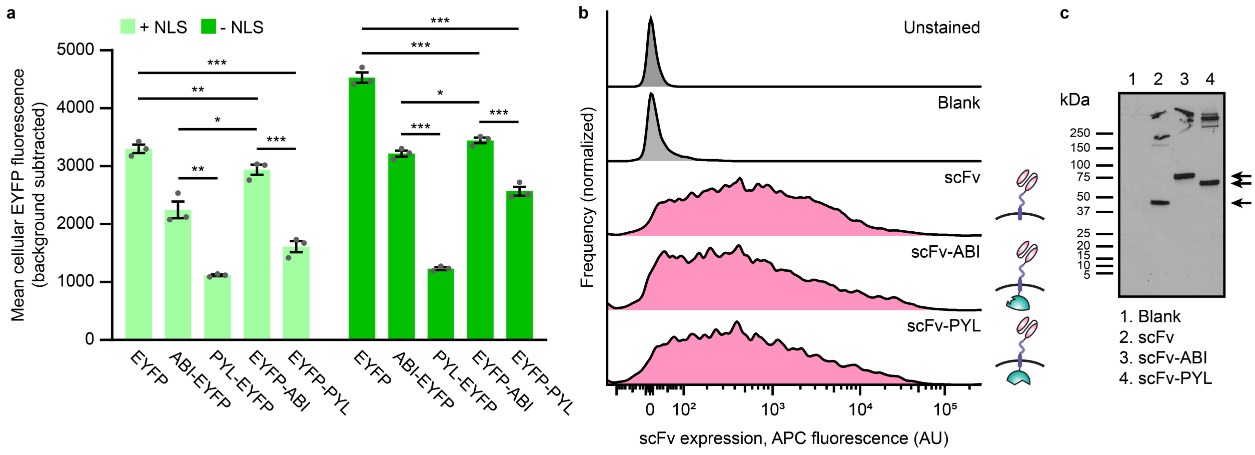


**Supplementary Fig. 10: ABA-binding domains can be incorporated into EV cargo proteins.** **a,** Expression of EYFP fused to the ABI and PYL ABA-binding domains with and without an NLS in transiently transfected HEK293FT cells analyzed by flow cytometry. Experiments were performed in biological triplicate, and error bars indicate standard error of the mean. Statistical tests comprise two-tailed Student's t-tests using the Benjamini-Hochberg method to reduce the false discovery rate. (*$p < 0.05$, **$p < 0.01$, ***$p < 0.001$). Exact p-values are reported in **Supplementary Table 1**. **b,** Surface stain (via 3x FLAG tag) of HEK293FTs transfected with anti-CD2 scFv constructs fused to ABI or PYL at the C-terminus. **c,** Expression of anti-CD2 scFv constructs from **b**. 2 µg cell lysate was loaded per lane. Expected band sizes: ~40, 62, and 75 kDa (arrows).

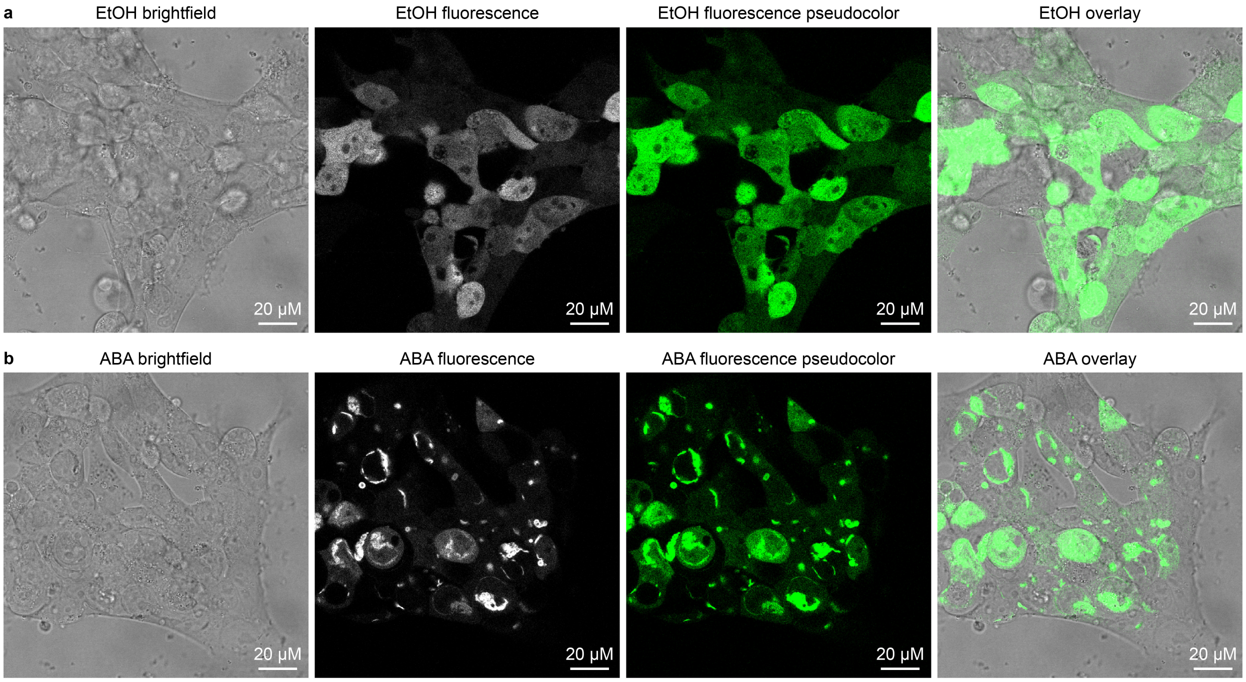


**Supplementary Fig. 11: ABA induces dimerization between the ABI and PYL domains. a**,**b**, HEK293FT cells transfected with anti-CD2 scFv-PYL and EYFP-ABI were treated with EtOH (**a**) or ABA (**b**) and imaged via confocal microscopy. Brightfield, fluorescence, contrast-adjusted and pseudo-colored fluorescence, and overlays are shown.

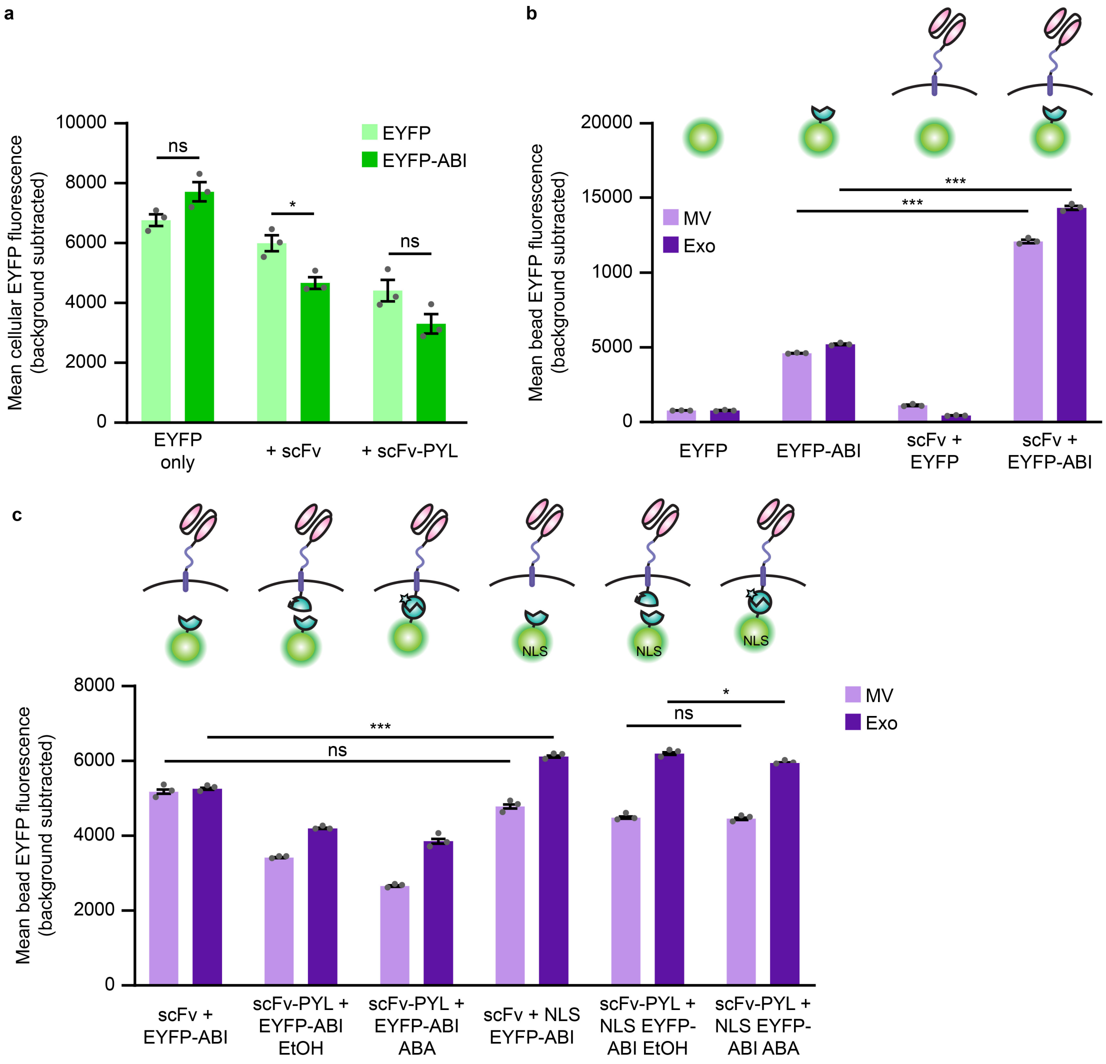


**Supplementary Fig. 12: The ABI domain increases EV cargo loading independent of total protein expression. a,** Expression of EYFP and EYFP-ABI in the presence of anti-CD2 targeting constructs in transiently transfected HEK293FT cells analyzed by flow cytometry. A key observation is that addition of the ABI domain does not increase overall cargo protein expression in producer cells. **b,** Repeat of EYFP-ABI EV loading trends in the presence of an scFv shown in **Fig. 3c**. **c,** Comparison of EYFP loading into EVs with and without an NLS with ABA-binding constructs and under ABA-induced dimerization conditions. Addition of an NLS did not substantially impact EYFP loading, nor did ABA-induced dimerization substantially impact loading of nuclear-localized cargo. Experiments were performed in biological triplicate, and error bars indicate standard error of the mean. Statistical tests comprise two-tailed Student's t-tests using the Benjamini-Hochberg method to reduce the false discovery rate. (*$p < 0.05$, **$p < 0.01$, ***$p < 0.001$). Exact p-values are reported in **Supplementary Table 1**.

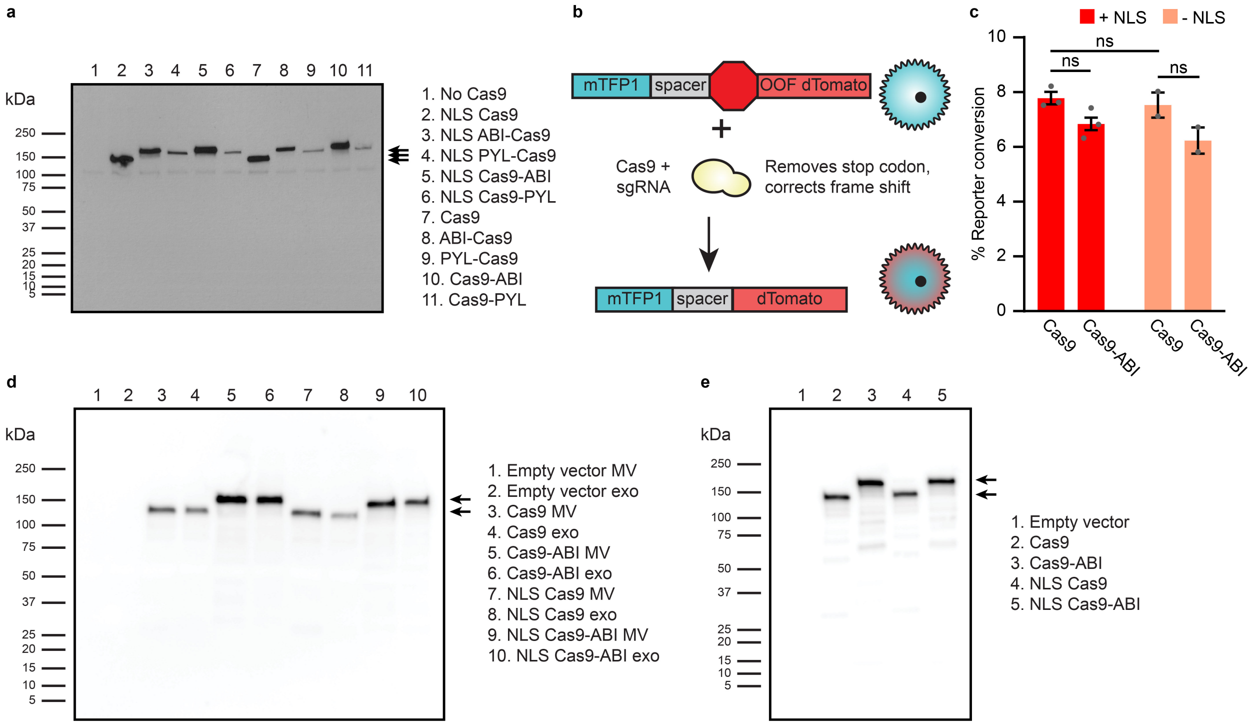


**Supplementary Fig. 13: The ABI domain increases Cas9 loading into EVs and Cas9-ABI retains function**. **a,** Expression of Cas9 fused to either the ABI or PYL domain in transiently transfected HEK293FT cells. 2 µg cell lysate was loaded per lane. Expected band sizes: ~160, 183, and 195 kDa (arrows). **b,** Cartoon illustrating the Cas9 reporter construct. Successful editing by Cas9 results in the deletion of a stop codon and (in some random fraction of cases) a repair-mediated frame shift induces express dTomato. **c,** Absence of an NLS or presence of the ABI domain does not meaningfully reduce Cas9 editing efficiency in transiently transfected Jurkat T cells. Cells were analyzed by flow cytometry 3 d post-transfection. Experiments were performed in biological triplicate, and error bars indicate standard error of the mean. Statistical tests comprise two-tailed Student's t-tests using the Benjamini-Hochberg method to reduce the false discovery rate. (*$p < 0.05$, **$p < 0.01$, ***$p < 0.001$). Exact p-values are reported in **Supplementary Table 1.** Samples with high cellular autofluorescence were excluded from analysis. **d,** Full blot of Cas9 EV active loading data presented in **Fig. 3e**. **e,** Cellular expression of Cas9 with and without the ABI domain or an NLS. 2 µg cell lysate was loaded per lane.

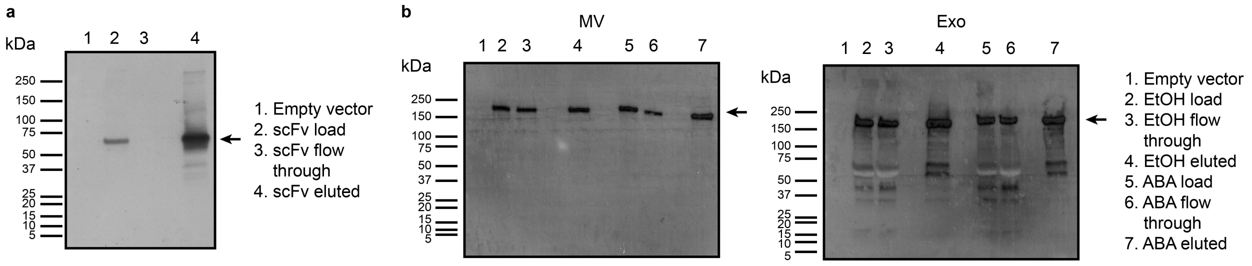


**Supplementary Fig. 14: EVs populations can be separated by affinity chromatography to analyze cargo loading patterns. a,** Validation of affinity chromatography technique. 3x FLAG tagged scFv containing vesicles were run through an anti-FLAG affinity matrix and analyzed for the FLAG tag to demonstrate enrichment in the eluted population. $1.5 \times 10^7$ EVs were loaded per lane. Expected band size: ~62 kDa (arrow). **b,** Full blots of affinity-isolated EV Cas9 content with and without ABA-induced dimerization presented in **Fig. 3f**.

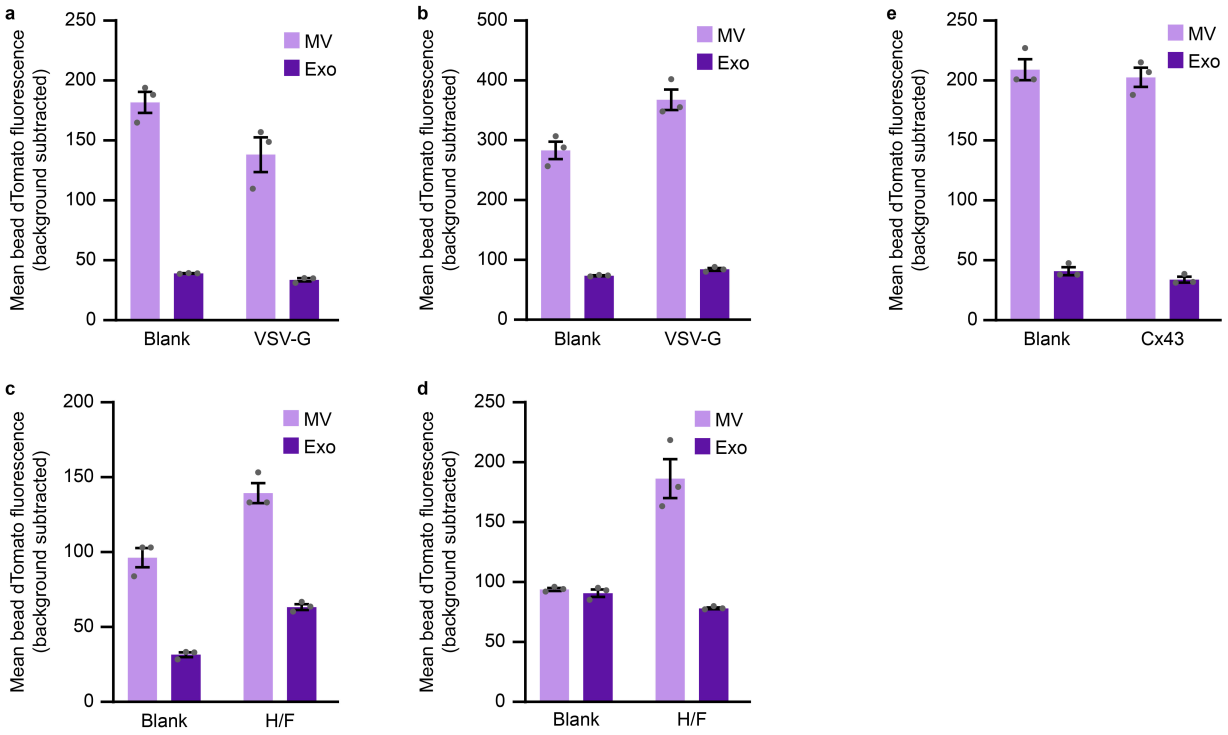


**Supplementary Fig. 15: EV dTomato loading evaluations for vesicle uptake and fusion experiments. a-e**, EV fluorescence controls for **Fig. 4b** (**a**), **Fig. 4c** (**b**), **Fig. 4e** (**c**), **Fig. 4f** (**d**), and **Supplementary Fig. 16** (**e**). EVs were adsorbed to aldehyde/sulfate latex beads and analyzed by flow cytometry to determine a bulk population fluorescence. Experiments were performed in biological triplicate, and error bars indicate standard error of the mean.

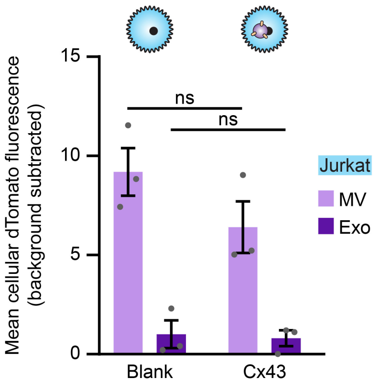


**Supplementary Fig. 16: Display of Cx43 on EVs does not lead to increased EV uptake by Jurkat T cells.** dTomato EVs were incubated with Jurkats for 16 h. Cells were trypsinized to remove surface-bound vesicles prior to analysis by flow cytometry. Experiments were performed in biological triplicate, and error bars indicate standard error of the mean. Statistical tests comprise two-tailed Student's t-tests using the Benjamini-Hochberg method to reduce the false discovery rate. (*$p < 0.05$, **$p < 0.01$, ***$p < 0.001$). Exact p-values are reported in **Supplementary Table 1**. EV dTomato loading evaluations can be found in **Supplementary Fig. 15e**.

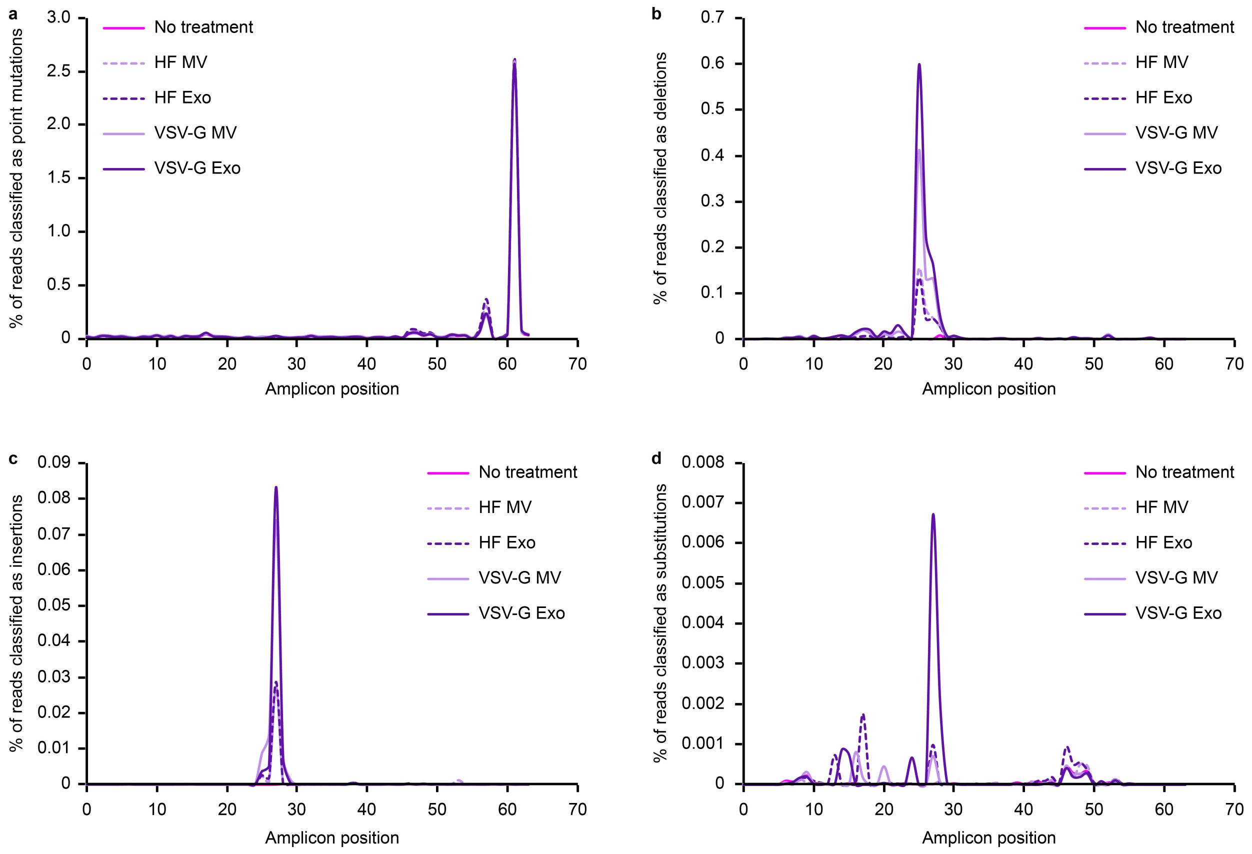


**Supplementary Fig. 17: EV-mediated Cas9 editing results in indels around the predicted CXCR4 cleavage site. a-d**, General point mutations (**a**), deletions (**b**), insertions (**c**) and substitutions (defined as edits that contain simultaneous insertions and deletions) (**d**) observed by HTS in primary human CD4$^{+}$ T cells as a function of percent of sequencing reads classified as edited. The general point mutations tallied in **a** did not differ with treatment and are thus not attributable to Cas9 or EVs. Each edit observed was classified uniquely into one of these four categories. Predicted Cas9 cut site was position 26 of the amplicon shown in **Figure 5**.

| Vesicle type | Viral glycoprotein | Active loading domain | Total % edited |
|---|---|---|---|
| No treatment | N/A | N/A | 0.00 |
| MV | Measles H/F | ABI | 0.03 |
| Exo | | | 0.03 |
| MV | VSV-G | | 0.25 |
| Exo | | | 0.16 |
| MV | Measles H/F | None | 0.02 |
| Exo | | | 0.01 |
| MV | VSV-G | | 0.14 |
| Exo | | | 0.21 |

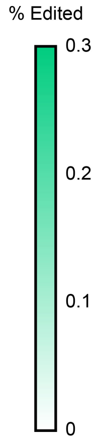


**Supplementary Fig. 18: Presence of the ABI active loading domain does not generally impact Cas9 editing efficiency.** 8.0x$10^9$ EVs were incubated per 4x$10^4$ $CD4^+$ T cells for 6 d prior to genomic DNA extraction and HTS analysis. Heat map coloring scales from 0-0.3% total Cas9 editing.

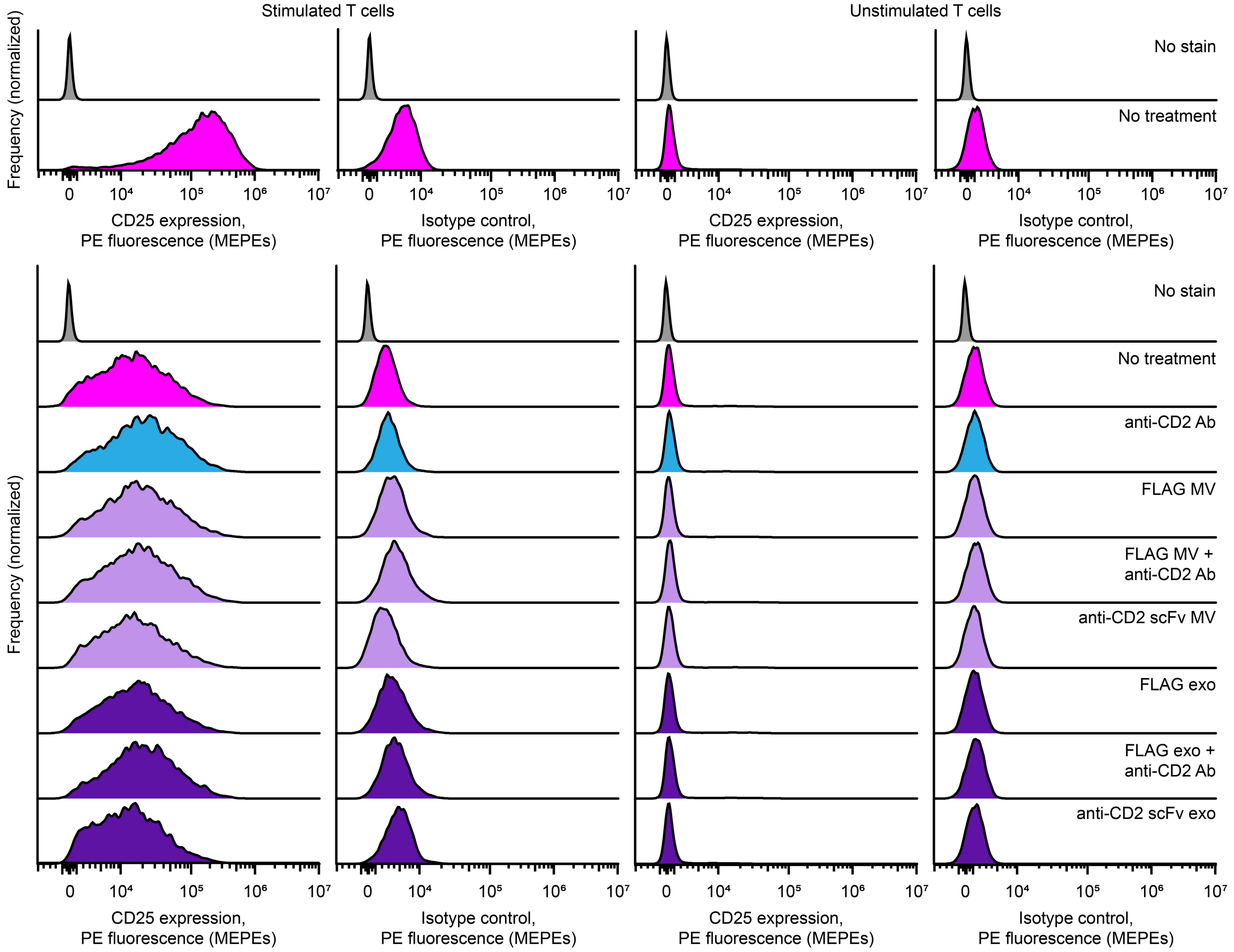


**Supplementary Fig. 19: CD2 engagement does not affect primary T cell activation state.** Primary human CD4$^+$ T cells were stained with anti-CD25 at the time of EV or anti-CD2 antibody treatment (upper) or 2 d post-treatment (lower). Unstimulated cells were used as a control for background anti-CD25 staining, and isotype controls were used to determine the impact of treatments on general cellular staining. Fluorescence was normalized using calibration beads to allow for signal comparison across days. MEPE: Molecules of equivalent PE.

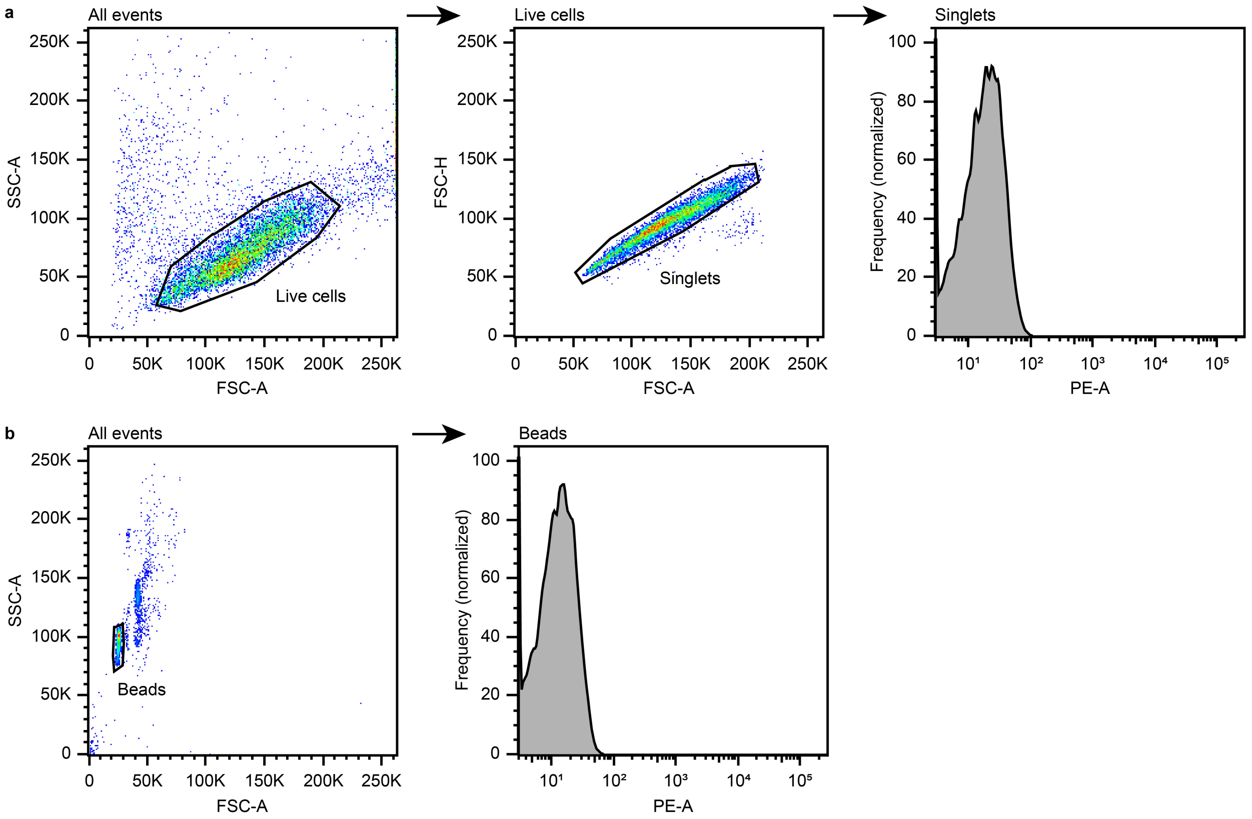


**Supplementary Fig. 20: Representative flow cytometry gating strategy for EV delivery experiments. a,** Live cells were identified based on their FSC-A vs SSC-A profile, and singlets were identified from live cells by their FSC-A vs FSC-H profile. Mean fluorescence intensity was quantified from singlets. **b,** Aldehyde/sulfate latex beads were identified based on their FSC-A vs SSC-A profile. Mean fluorescence intensity was quantified from beads.

**Supplementary Note 1: Summary of HTS analysis code workflow**

Code used for performing HTS analysis is deposited on GitHub as described in **Methods**. Here, we summarize the strategy used to parse HTS data and calculate and classify edits.

The raw data are provided as Read 1/Read 2 (R1/R2) pairs, which represent the 5' and 3' ends of a complete fragment. Analysis is based upon the assumption that the template structure is as follows from left to right: first, a 5-nt index segment; a forward primer; the 64-nt target sequence; a reverse primer; and finally, another 5-nt index segment. These are described in code and passed to the segment processor, which, for each R1/R2 pair, first determines the associated fragment and then uses pattern-matching techniques to determine the best possible match to the template, taking into account mutations and indels. Best fits are determined using standard string-edit-distance techniques. For each fragment that matches the template with sufficient accuracy, any errors (mutations or indels) are tabulated including the site of the error and associated lengths/sequences. These values are tabulated across each sample and are then used to generate the raw CSV data and plots as outputs.

**Supplementary Table 1: Statistical test p-values**

| Figure | Comparison | p value |
|---|---|---|
| Fig. 2b | Non-targeted vs helical linker scFv MV | 9.38E-05 |
| | Non-targeted vs 40 GS linker scFv MV | 2.36E-02 |
| | Non-targeted vs IgG4 hinge linker scFv MV | 1.09E-02 |
| | Non-targeted vs helical linker scFv exo | 6.32E-03 |
| | Non-targeted vs 40 GS linker scFv exo | 1.01E-02 |
| | Non-targeted vs IgG4 hinge linker scFv exo | 7.40E-03 |
| Fig. 2d | Trypsin cases: Non-targeted vs targeted MV | 3.74E-04 |
| | Trypsin cases: Non-targeted vs targeted exo | 1.82E-02 |
| Fig. 2e | Blocking cases: Non-targeted vs targeted MV | 3.20E-02 |
| | Blocking cases: Non-targeted vs targeted exo | 3.92E-01 |
| Fig. 2f | Original vs optimized MV | 1.18E-03 |
| | Original vs optimized exo | 3.28E-02 |
| Fig. 2g | Non-targeted vs targeted MV | 2.01E-02 |
| | Non-targeted vs targeted exo | 2.05E-02 |
| Fig. 2h | Trypsin cases: Non-targeted vs targeted MV | 3.95E-03 |
| | Trypsin cases: Non-targeted vs targeted exo | 5.28E-01 |
| Fig. 3c | EYFP vs EYFP ABI MV | 5.28E-08 |
| | EYFP vs EYFP ABI exo | 2.80E-06 |
| | scFv cases: EYFP vs EYFP-ABI MV | 4.96E-07 |
| | scFv cases: EYFP vs EYFP-ABI exo | 5.02E-07 |
| | scFv-PYL + EYFP-ABI cases: EtOH vs ABA MV | 7.94E-01 |
| | scFv-PYL + EYFP-ABI cases: EtOH vs ABA exo | 2.15E-02 |
| Fig. 4b | Blank vs VSV-G MV | 1.27E-07 |
| | Blank vs VSV-G exo | 1.92E-09 |
| Fig. 4c | Blank vs VSV-G MV | 1.10E-04 |
| | Blank vs VSV-G exo | 1.33E-05 |
| Fig. 4e | - SLAM cases: Blank vs H/F MV | 1.44E-03 |
| | - SLAM cases: Blank vs H/F exo | 3.27E-02 |
| | + SLAM cases: Blank vs H/F MV | 1.99E-06 |
| | + SLAM cases: Blank vs H/F exo | 1.89E-04 |
| Fig. 4f | Blank vs H/F MV | 6.82E-03 |
| | Blank vs H/F exo | 8.52E-04 |
| Supplementary Fig. 6 | Non-targeted vs non-targeted + blocking MV | 1.09E-01 |
| | Non-targeted vs non-targeted + blocking exo | 2.09E-01 |
| | Targeted vs targeted + blocking MV | 7.58E-01 |
| | Targeted vs targeted + blocking exo | 2.10E-01 |
| Supplementary Fig. 9d | PDGFR display vs C1C2 display MV | 4.71E-01 |
| | PDGFR display vs C1C2 display exo | 1.43E-05 |
| Supplementary Fig. 9f | Blocking cases: Non-targeted vs targeted MV | 3.85E-02 |
| | Blocking cases: Non-targeted vs targeted exo | 9.36E-02 |
| Supplementary Fig. 10a | NLS EYFP vs NLS EYFP-ABI | 3.49E-02 |

| | | |
|---|---|---|
| | NLS EYFP vs NLS EYFP-PYL | 1.47E-04 |
| | NLS ABI-EYFP vs NLS EYFP-ABI | 1.53E-02 |
| | NLS ABI-EYFP vs NLS PYL-EYFP | 1.51E-03 |
| | NLS EYFP-ABI vs NLS EYFP-PYL | 5.27E-04 |
| | EYFP vs EYFP-ABI | 4.41E-04 |
| | EYFP vs EYFP-PYL | 7.87E-05 |
| | ABI-EYFP vs EYFP-ABI | 3.27E-02 |
| | ABI-EYFP vs PYL-EYFP | 4.53E-06 |
| | EYFP-ABI vs EYFP-PYL | 6.40E-04 |
| Supplementary Fig. 12a | EYFP vs EYFP-ABI | 6.53E-02 |
| | scFv cases: EYFP vs EYFP-ABI | 1.62E-02 |
| | scFv-PYL cases: EYFP vs EYFP-ABI | 8.45E-02 |
| Supplementary Fig. 12b | EYFP-ABI vs scFv + EYFP-ABI MV | 3.67E-07 |
| | EYFP-ABI vs scFv + EYFP-ABI exo | 3.93E-07 |
| Supplementary Fig. 12c | scFv cases: EYFP-ABI vs NLS EYFP-ABI MV | 3.95E-02 |
| | scFv cases: EYFP-ABI vs NLS EYFP-ABI exo | 1.78E-04 |
| | scFv-PYL + NLS EYFP-ABI cases: EtOH vs ABA MV | 6.85E-01 |
| | scFv-PYL + NLS EYFP-ABI cases: EtOH vs ABA exo | 1.70E-02 |
| Supplementary Fig. 13c | NLS Cas9 vs Cas9 | 6.14E-01 |
| | NLS Cas9 vs NLS Cas9-ABI | 6.56E-02 |
| | Cas9 vs Cas9-ABI | 6.14E-01 |
| Supplementary Fig. 16 | Blank vs Cx43 MV | 1.85E-01 |
| | Blank vs Cx43 exo | 8.08E-01 |
