## Supplementary figures and images for "Bioengineering multifunctional extracellular vesicles for targeted delivery of biologics to T cells"

### Fig 3b ABA and Supplementary Fig 11b.tif

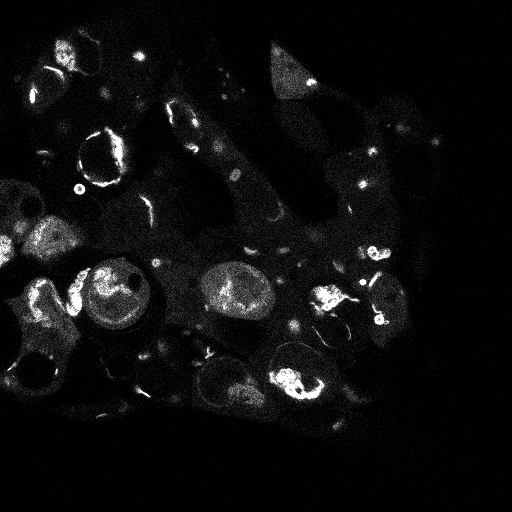

### Fig 3b EtOH and Supplemenatry Fig 11a.tif

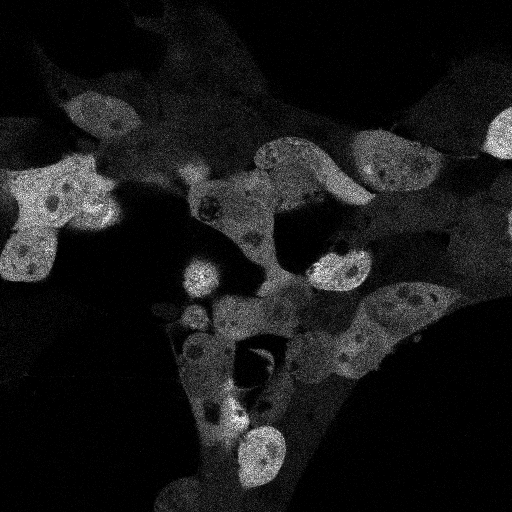

### Fig 3e and Supplementary Fig 13d.tif

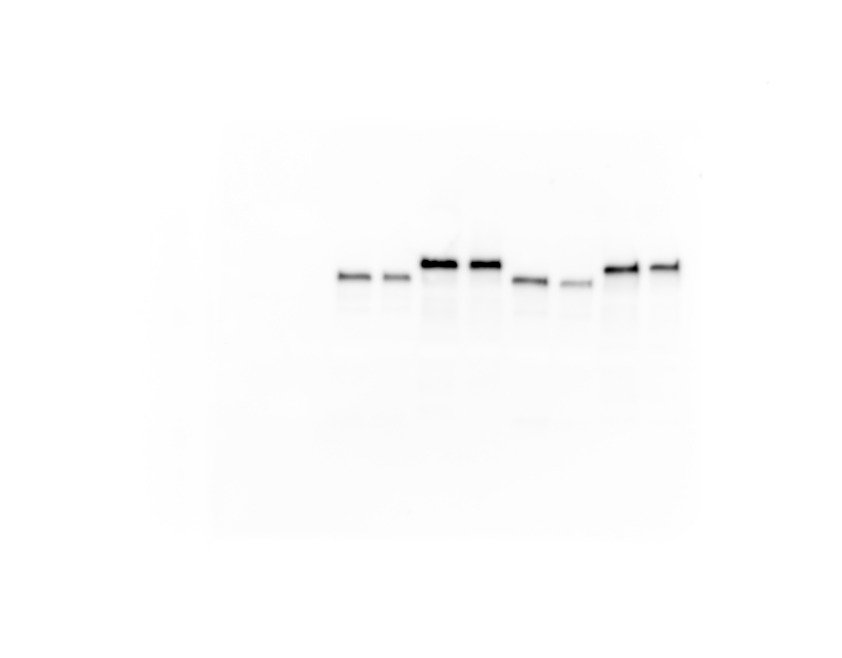

### Fig 3f and Supplementary Fig 14b.jpg

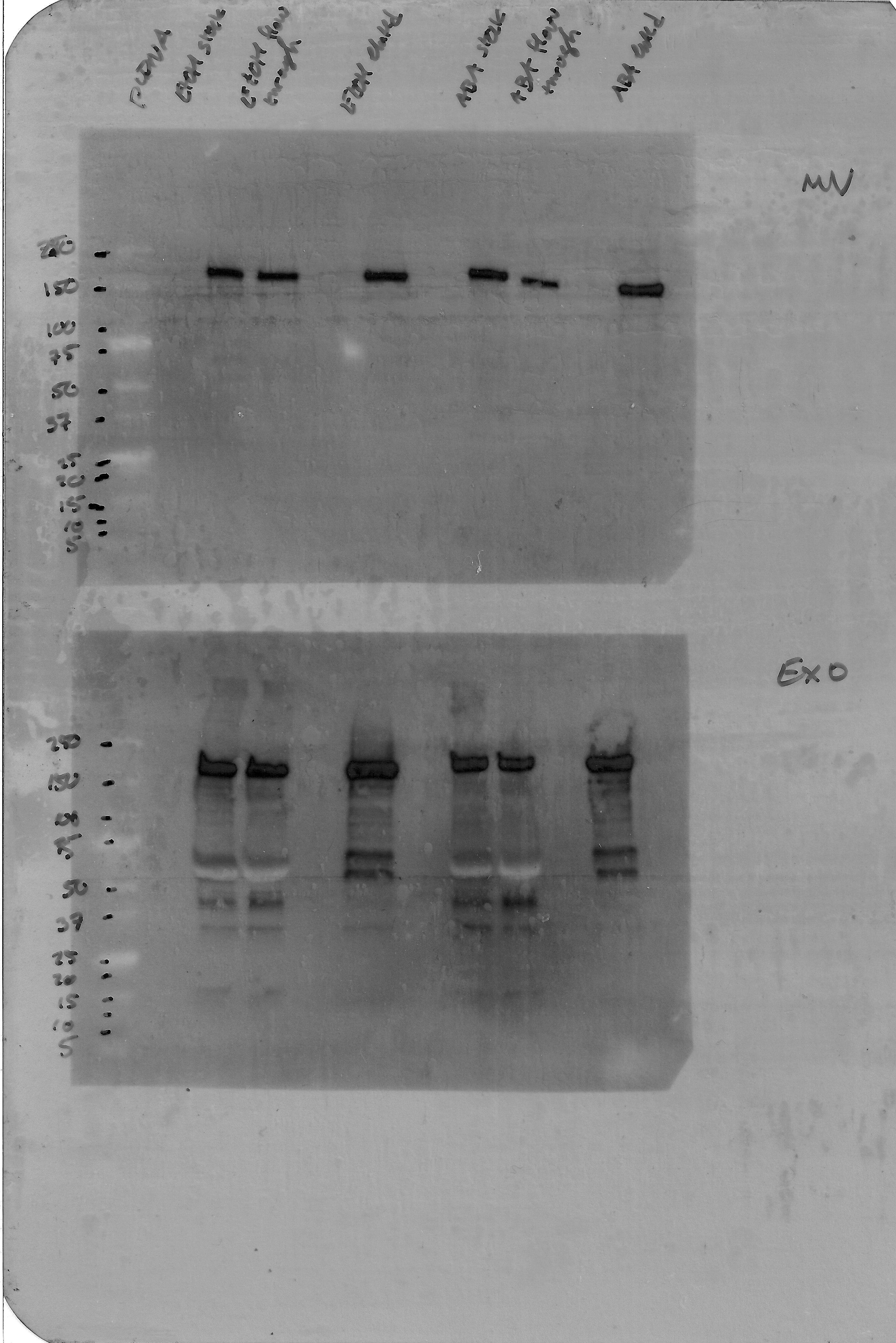

### Fig 3g exo.jpg

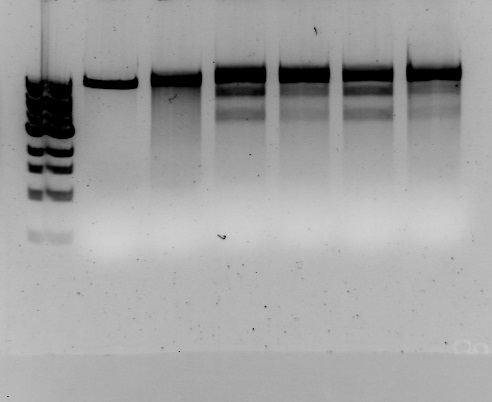

### Fig 3g MV.jpg

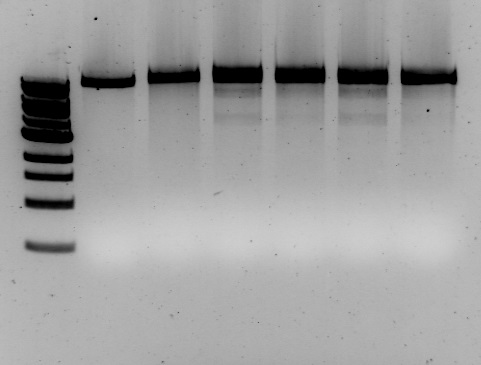

### Supplementary Fig 1b (right).jpg

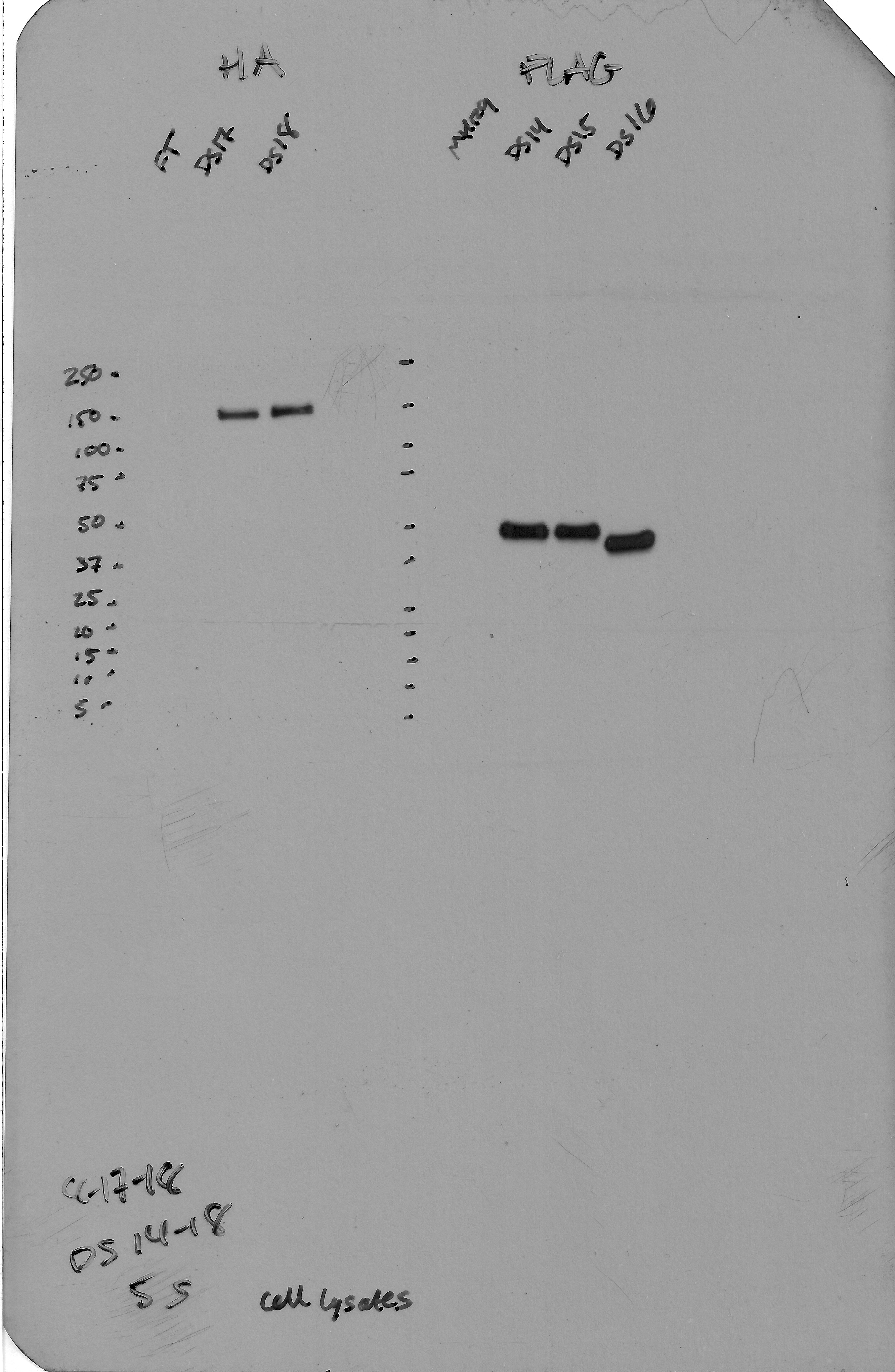

### Supplementary Fig 1c (top).jpg

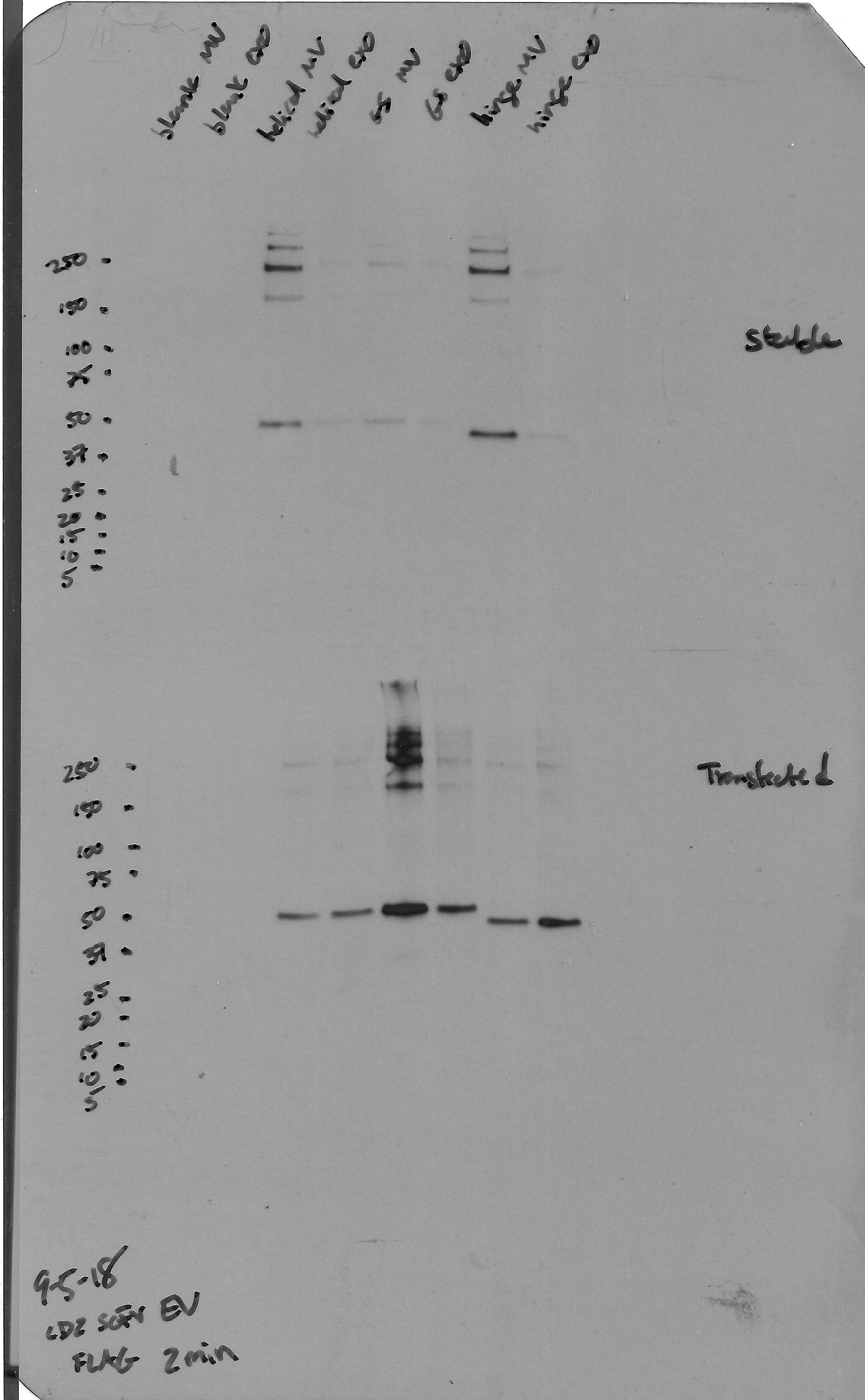

### Supplementary Fig 2a Alix.tif

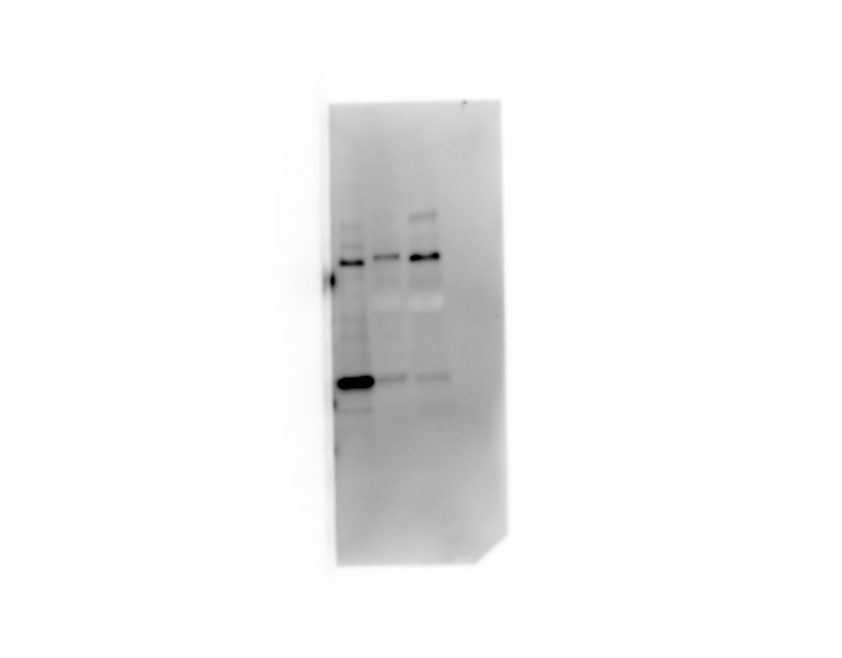

### Supplementary Fig 2a calnexin.tif

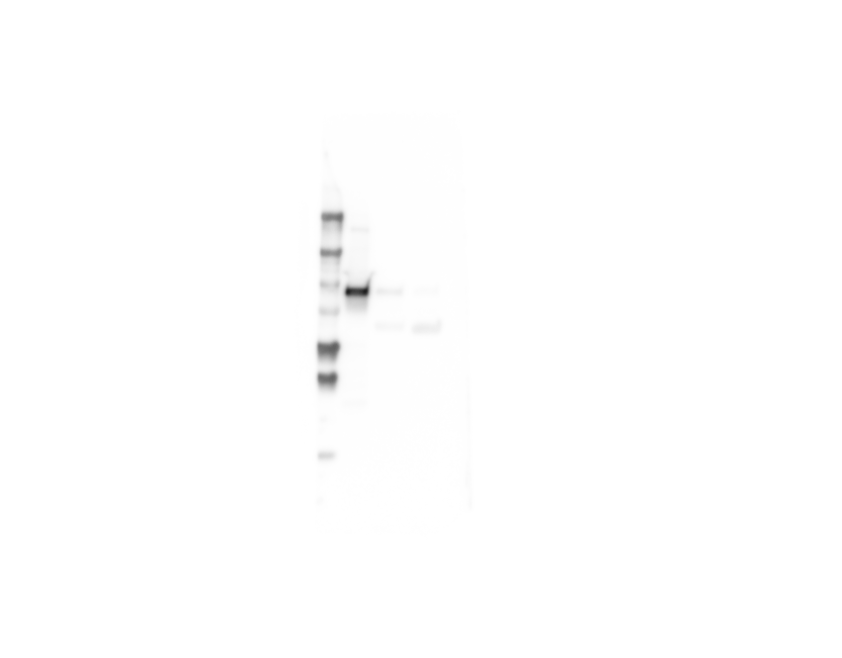

### Supplementary Fig 2a CD9.tif

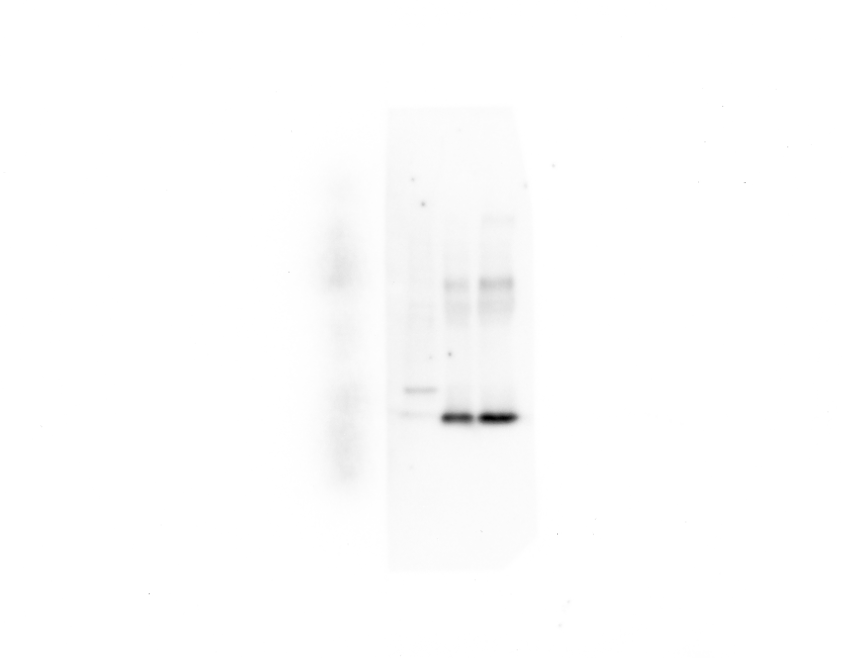

### Supplementary Fig 2a CD81.tif

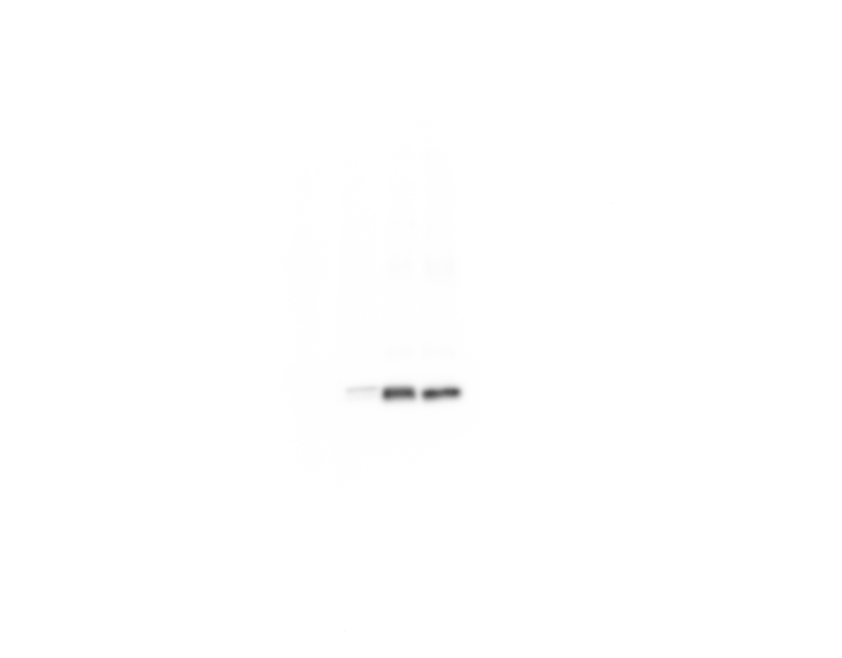

### Supplementary Fig 2c exo.tif

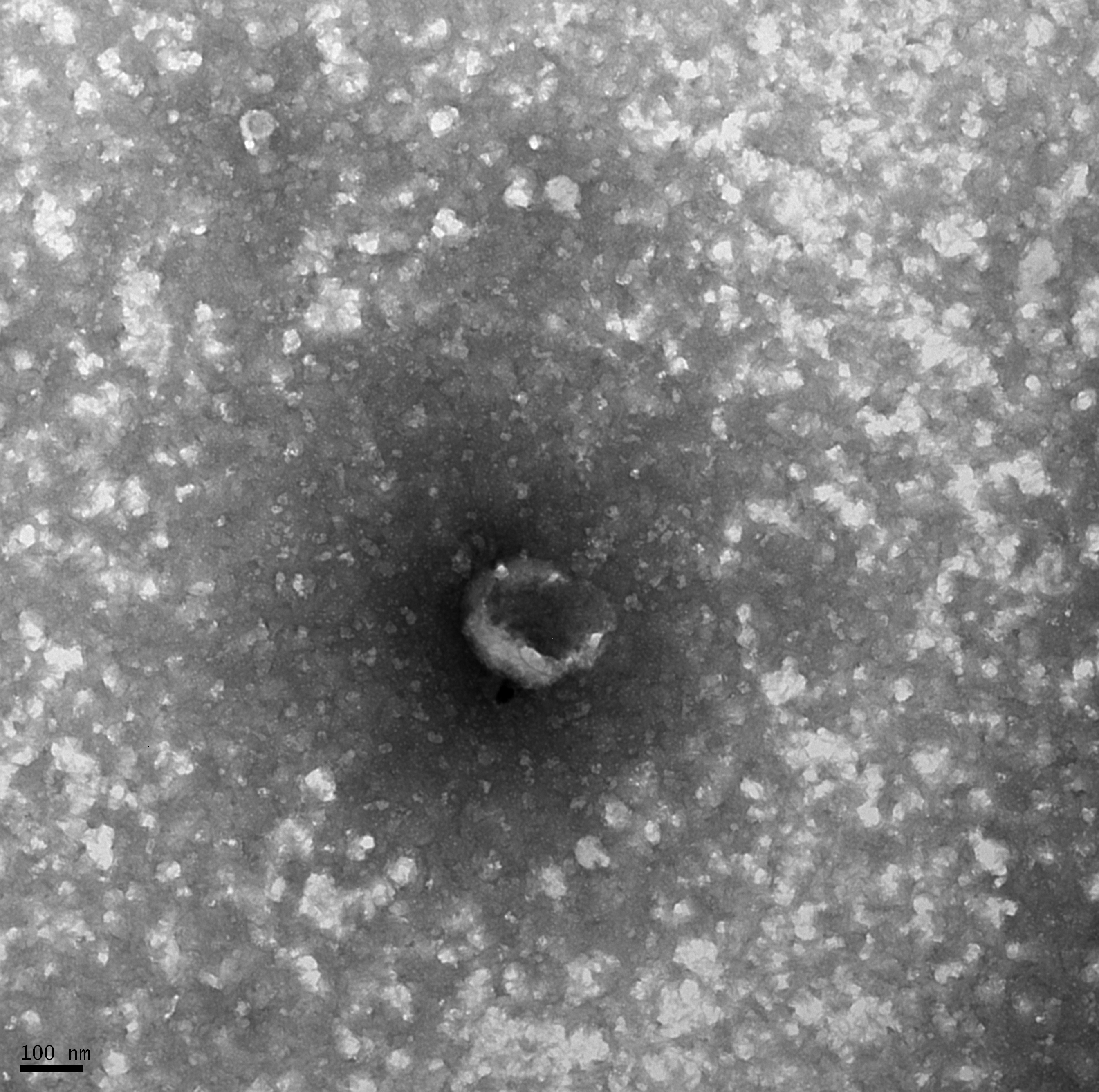

### Supplementary Fig 2c MV.tif

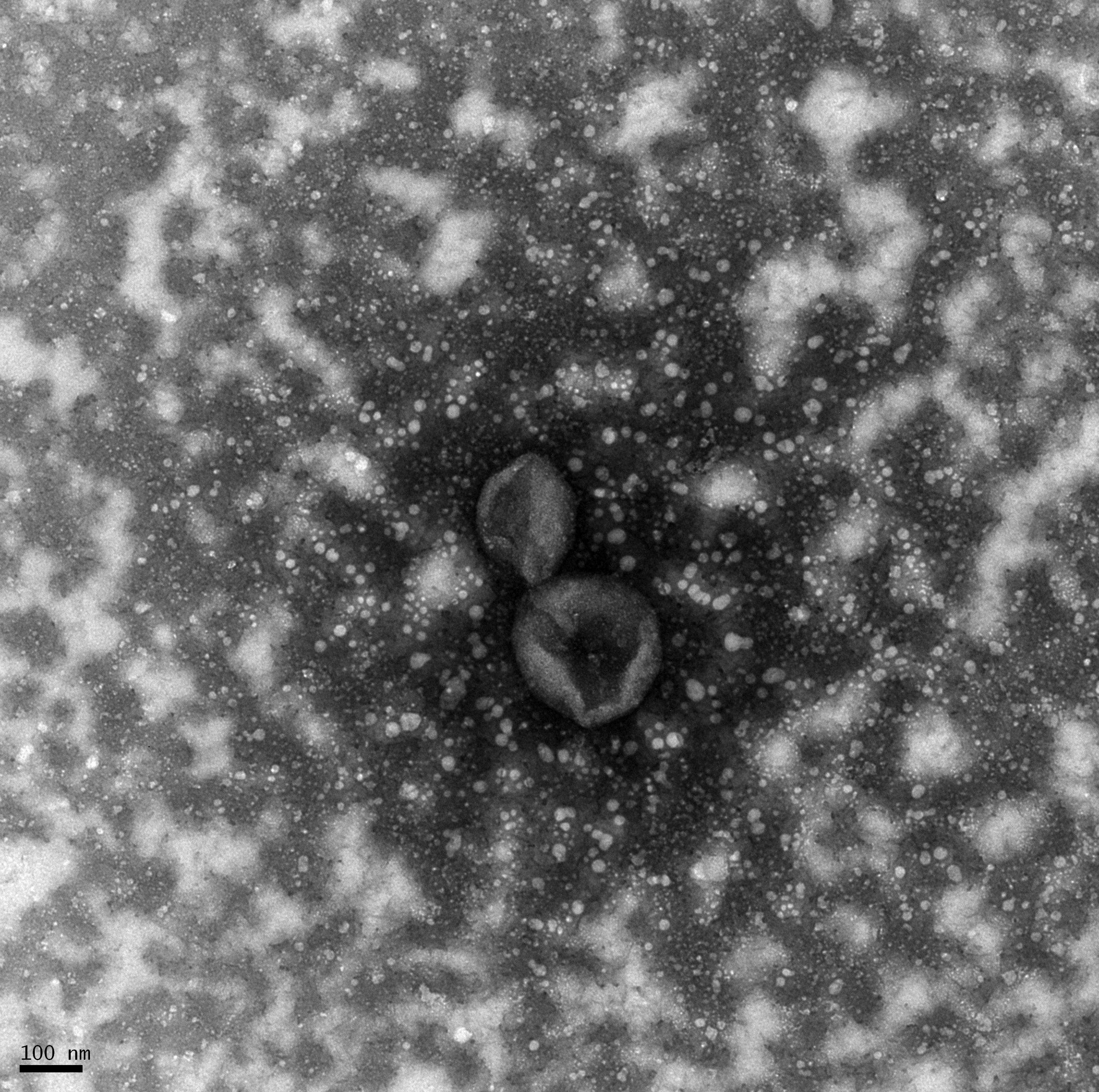

### Supplementary Fig 7a (right).jpg

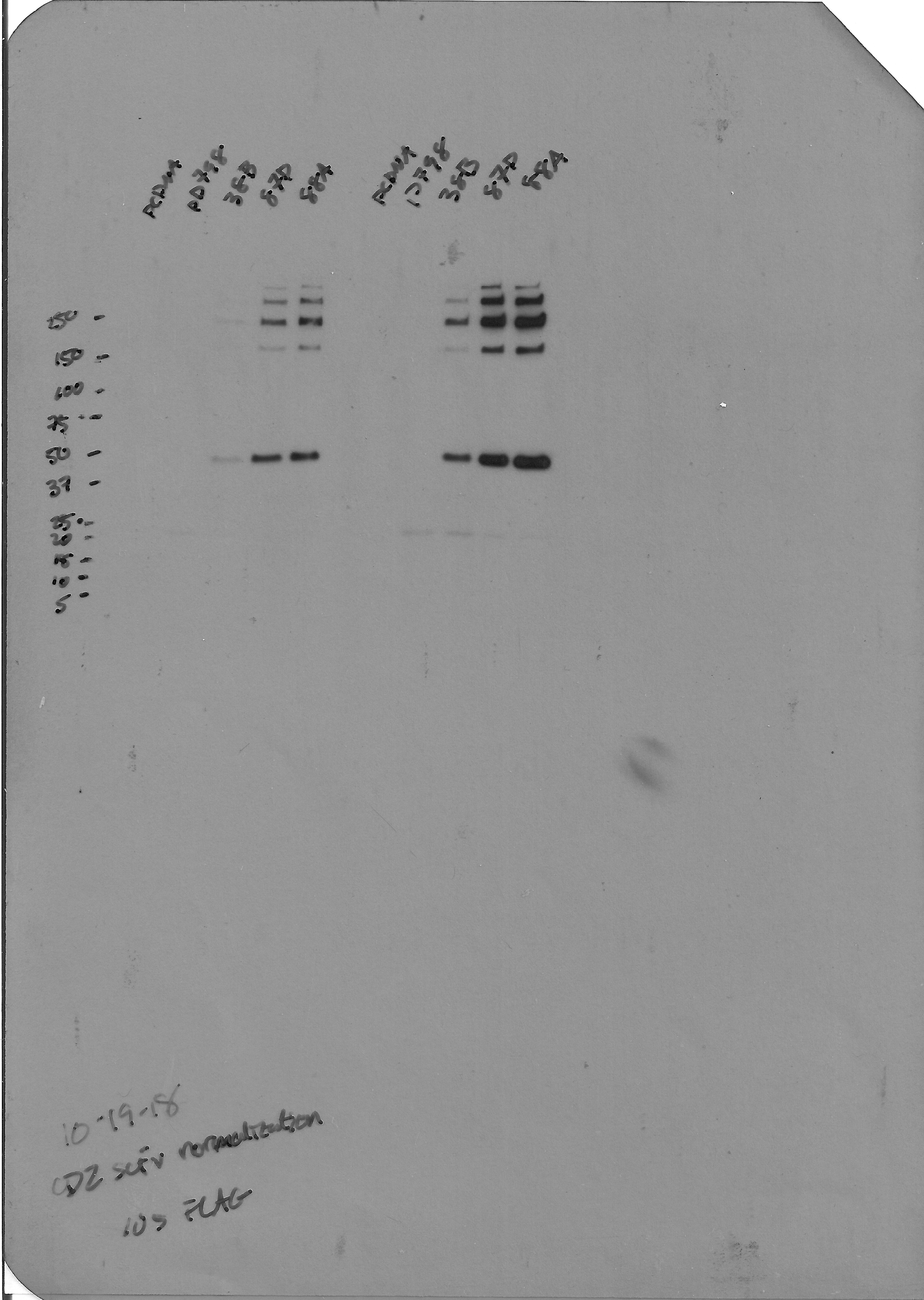

### Supplementary Fig 7b (left).tif

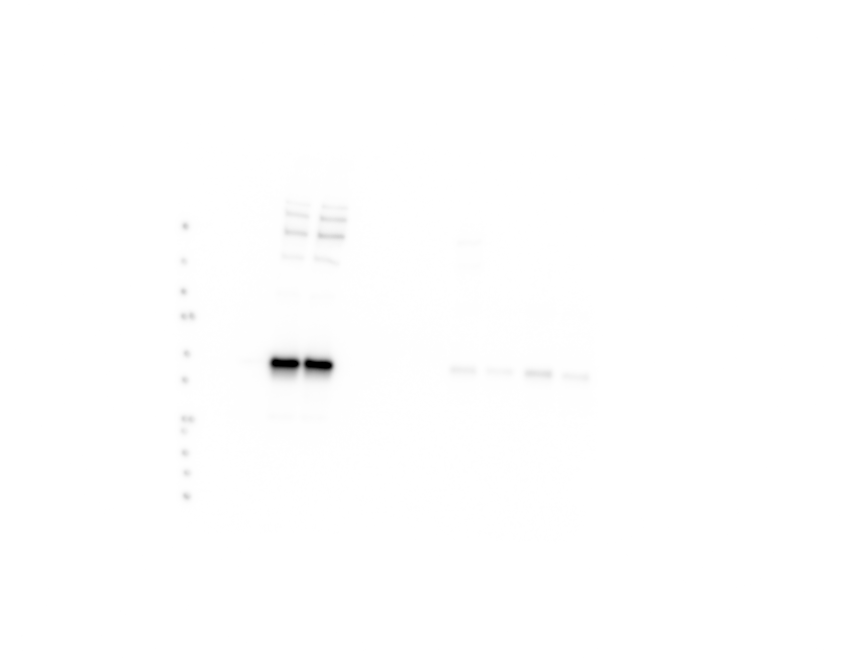

### Supplementary Fig 7c.tif

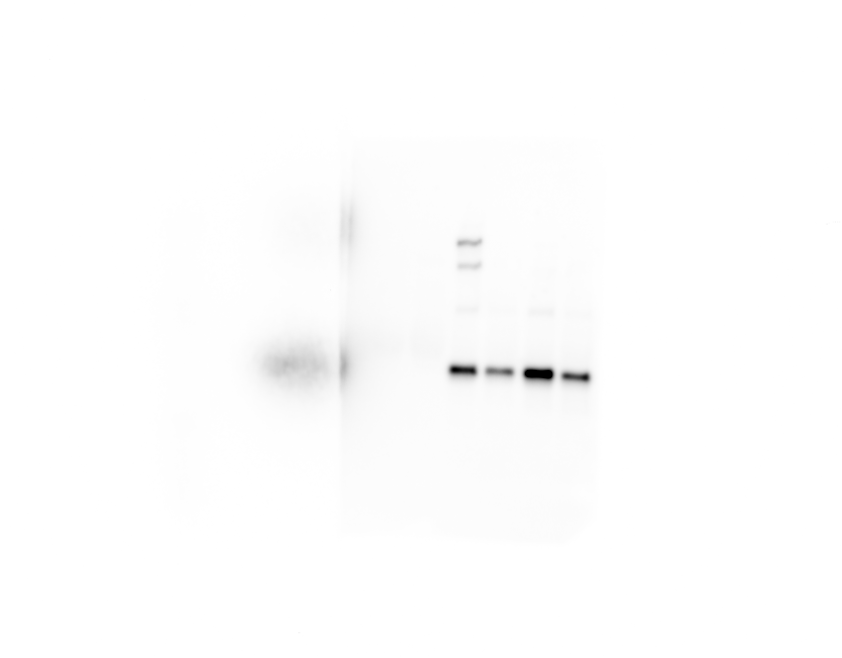

### Supplementary Fig 7e exo.tif

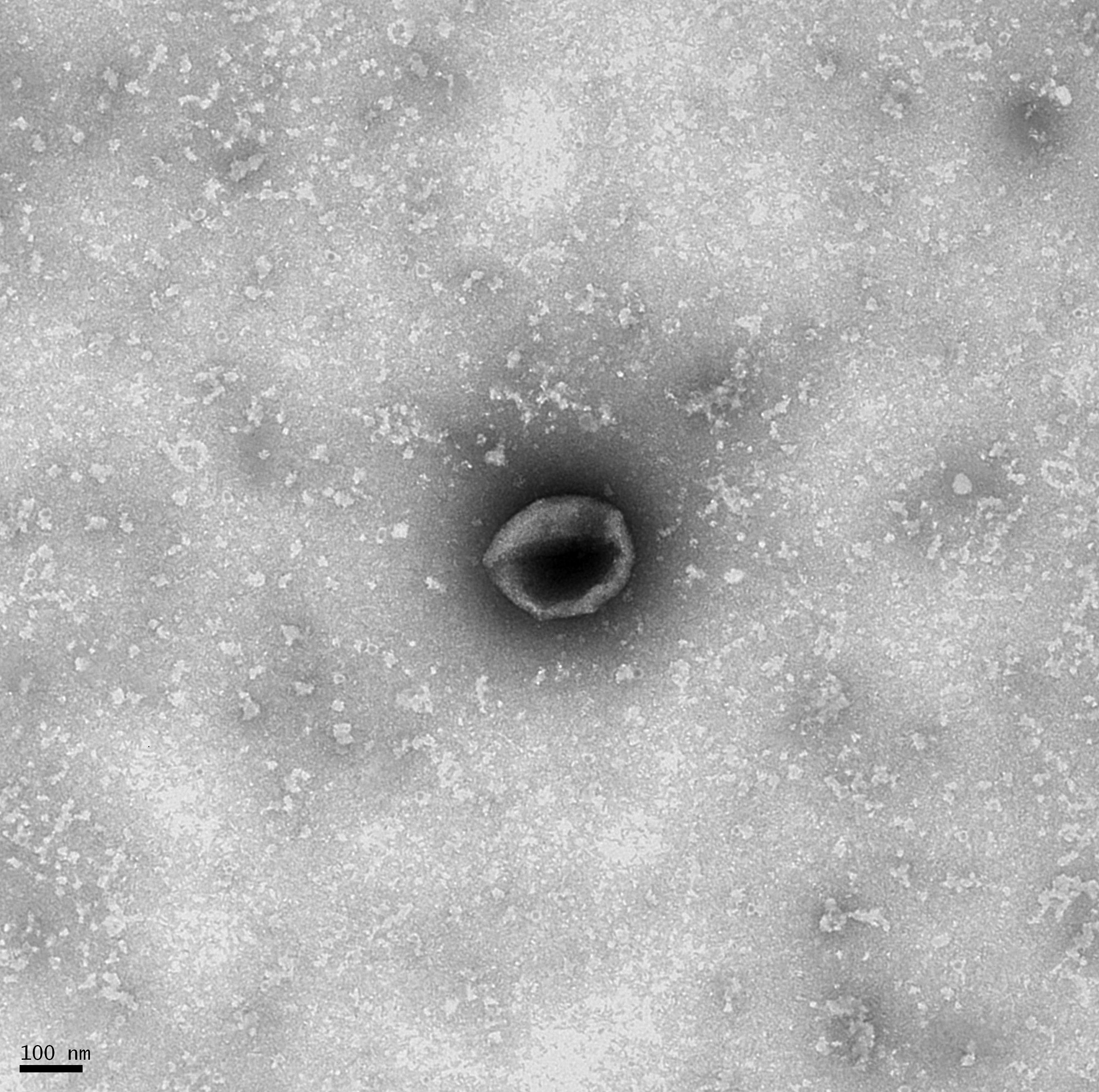

### Supplementary Fig 7e MV.tif

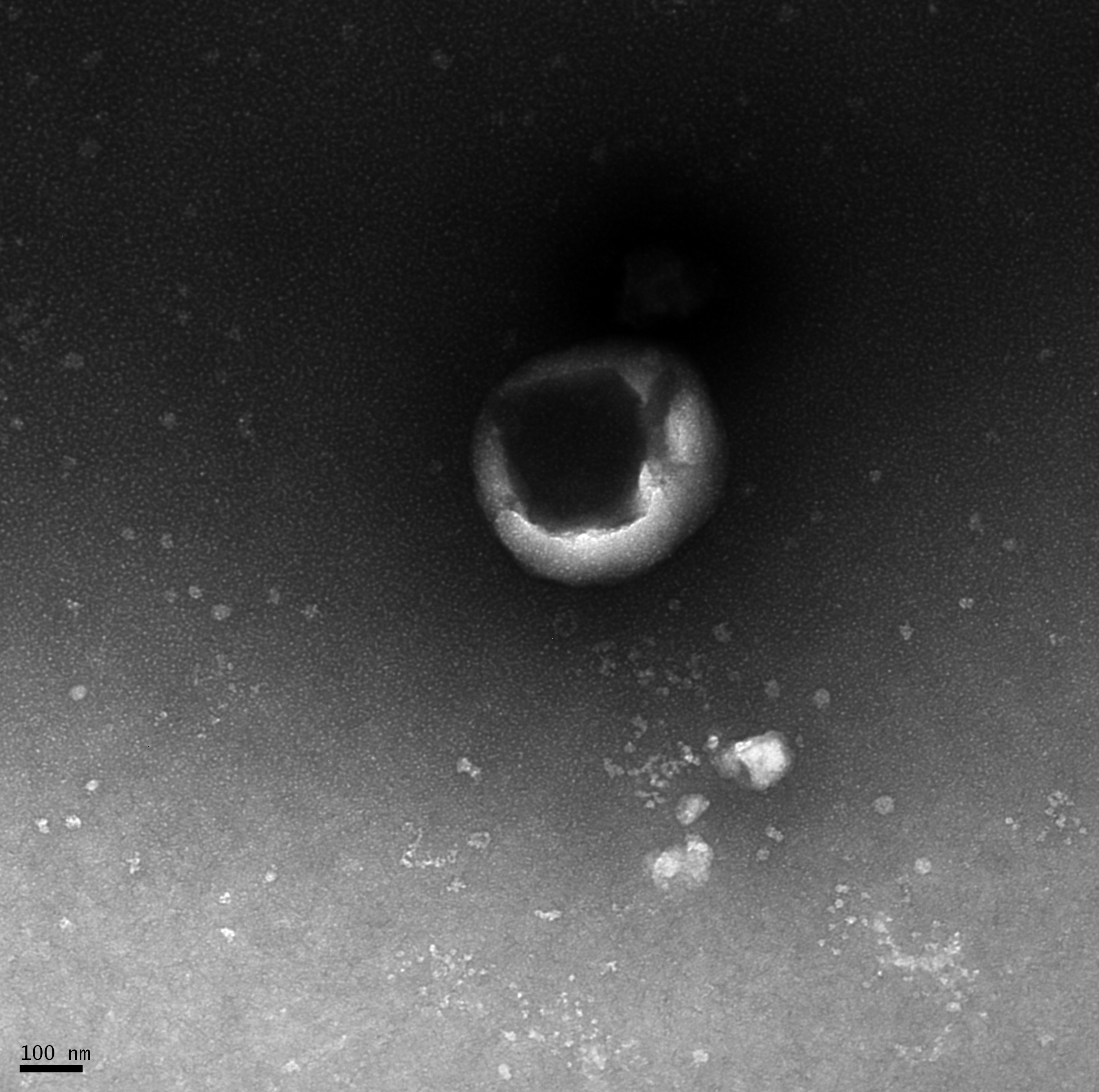

### Supplementary Fig 9b (left).jpg

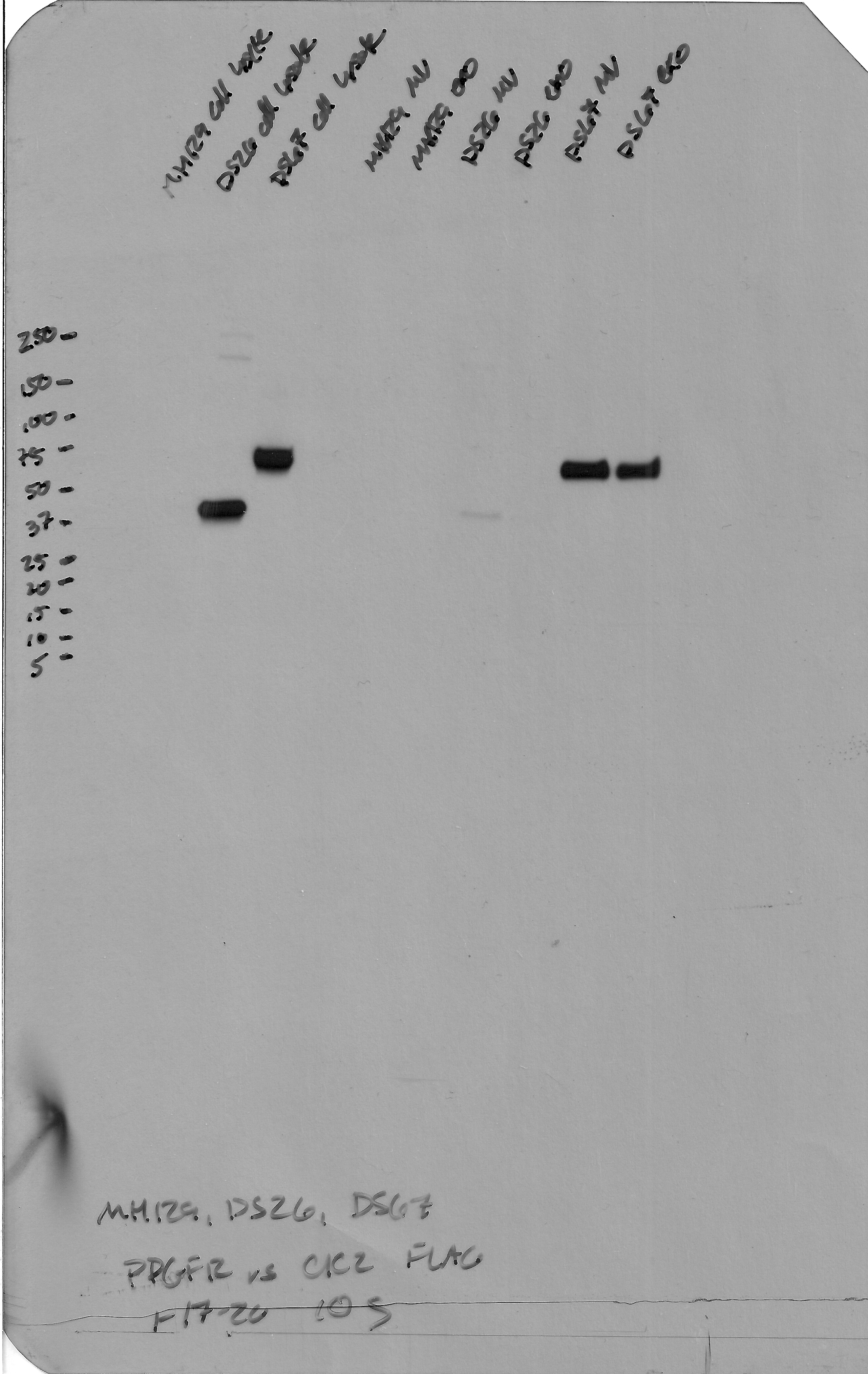

### Supplementary Fig 9c (right).jpg

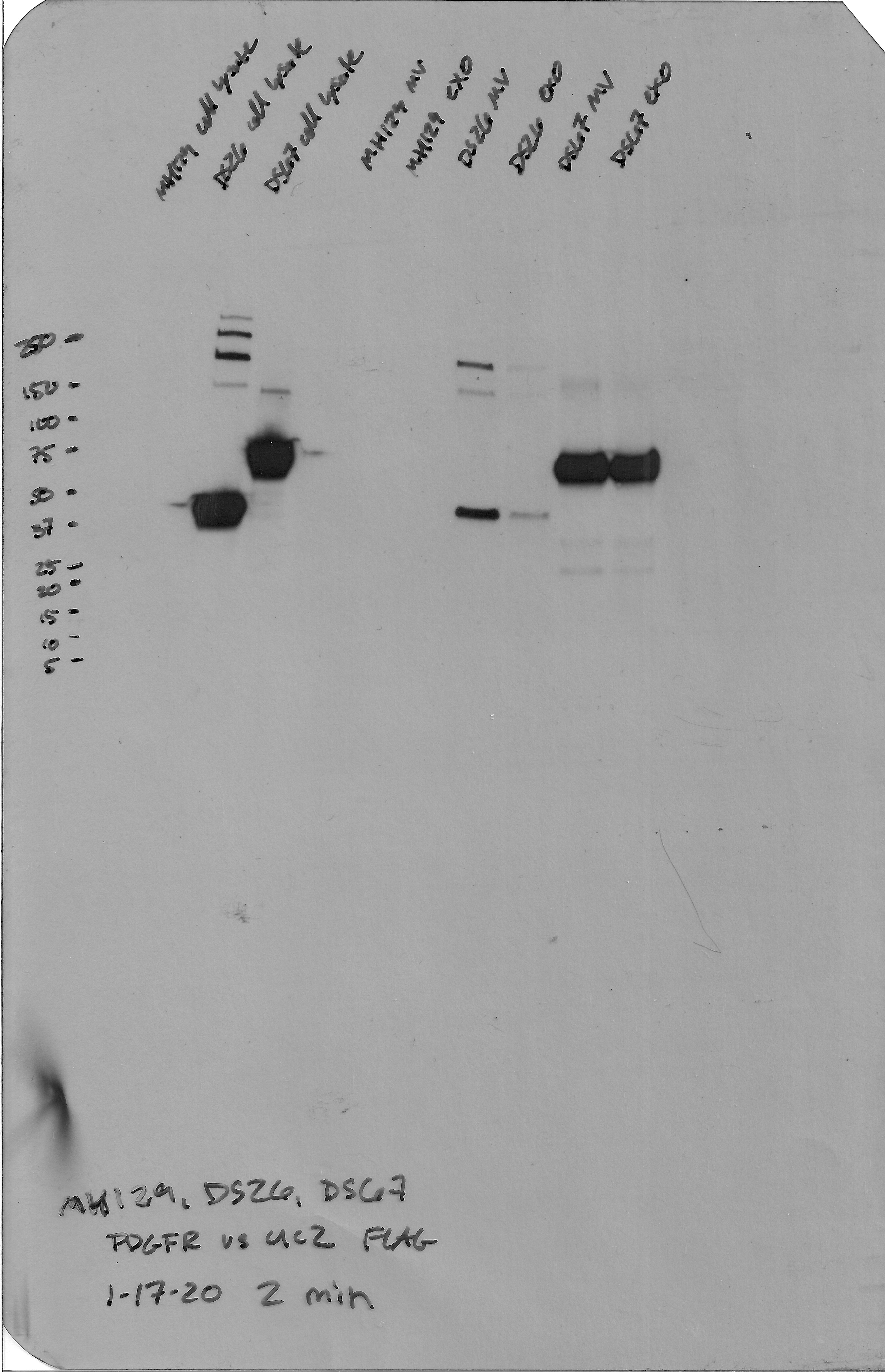

### Supplementary Fig 10c (top).jpg

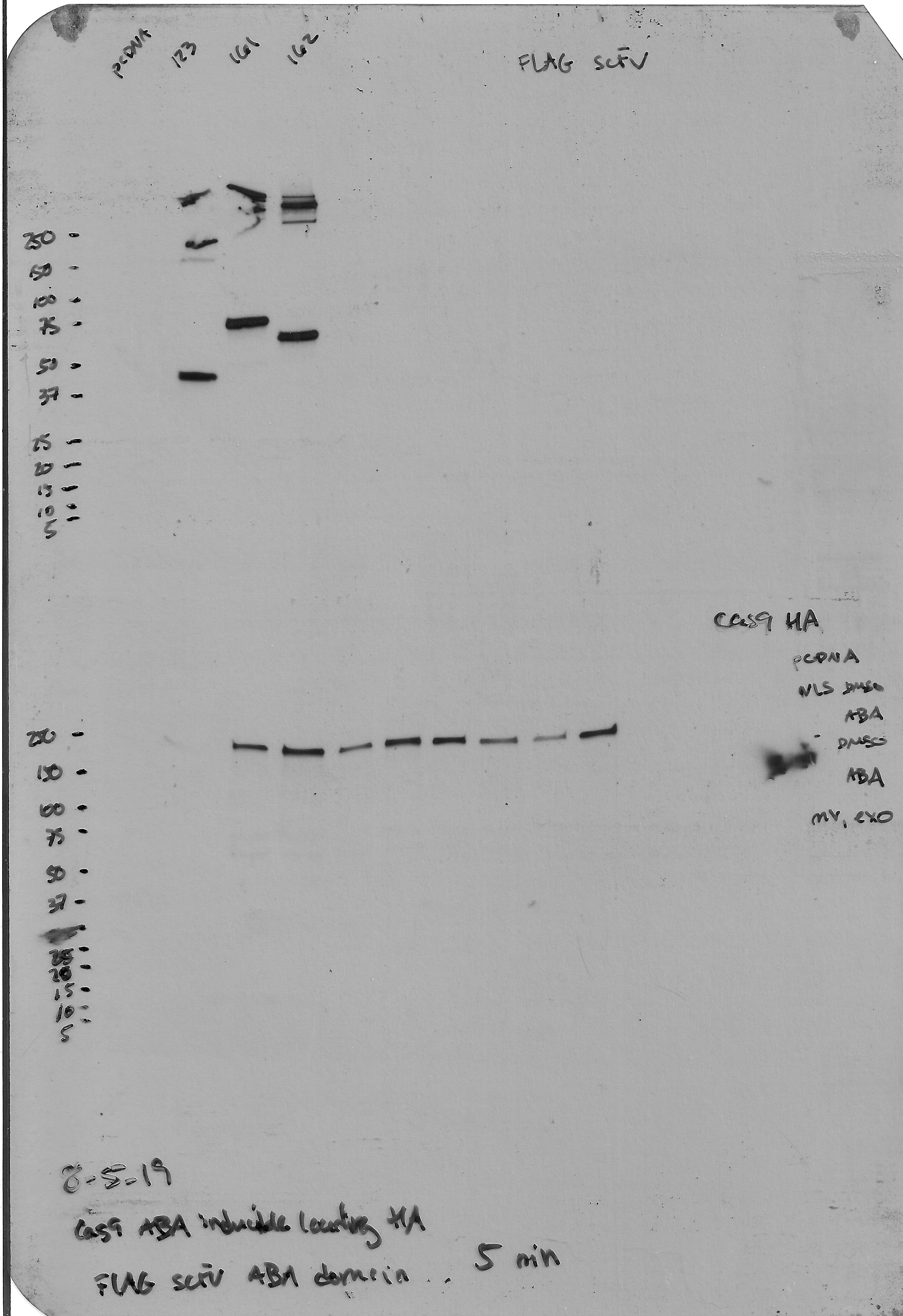

### Supplementary Fig 11a.tif

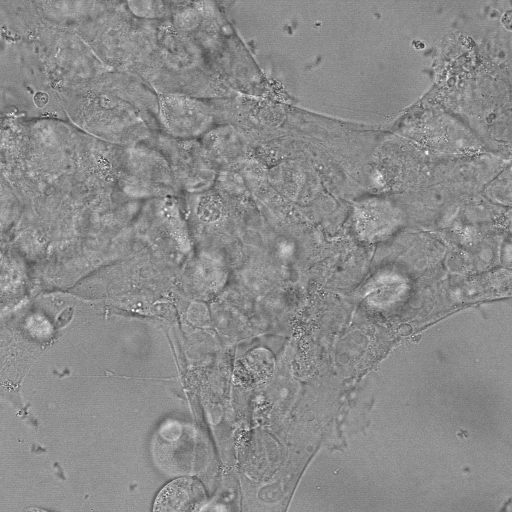

### Supplementary Fig 11b.tif

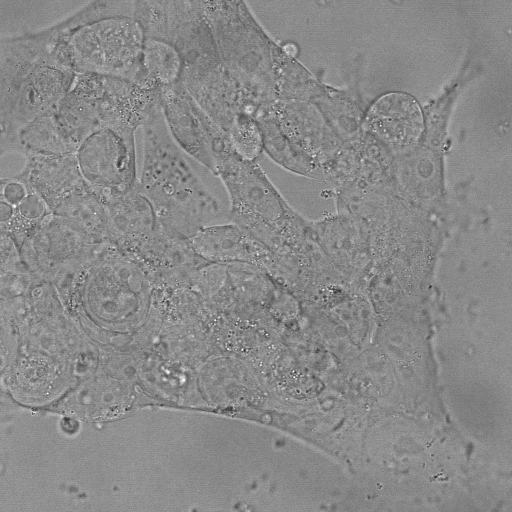

### Supplementary Fig 13a (right).jpg

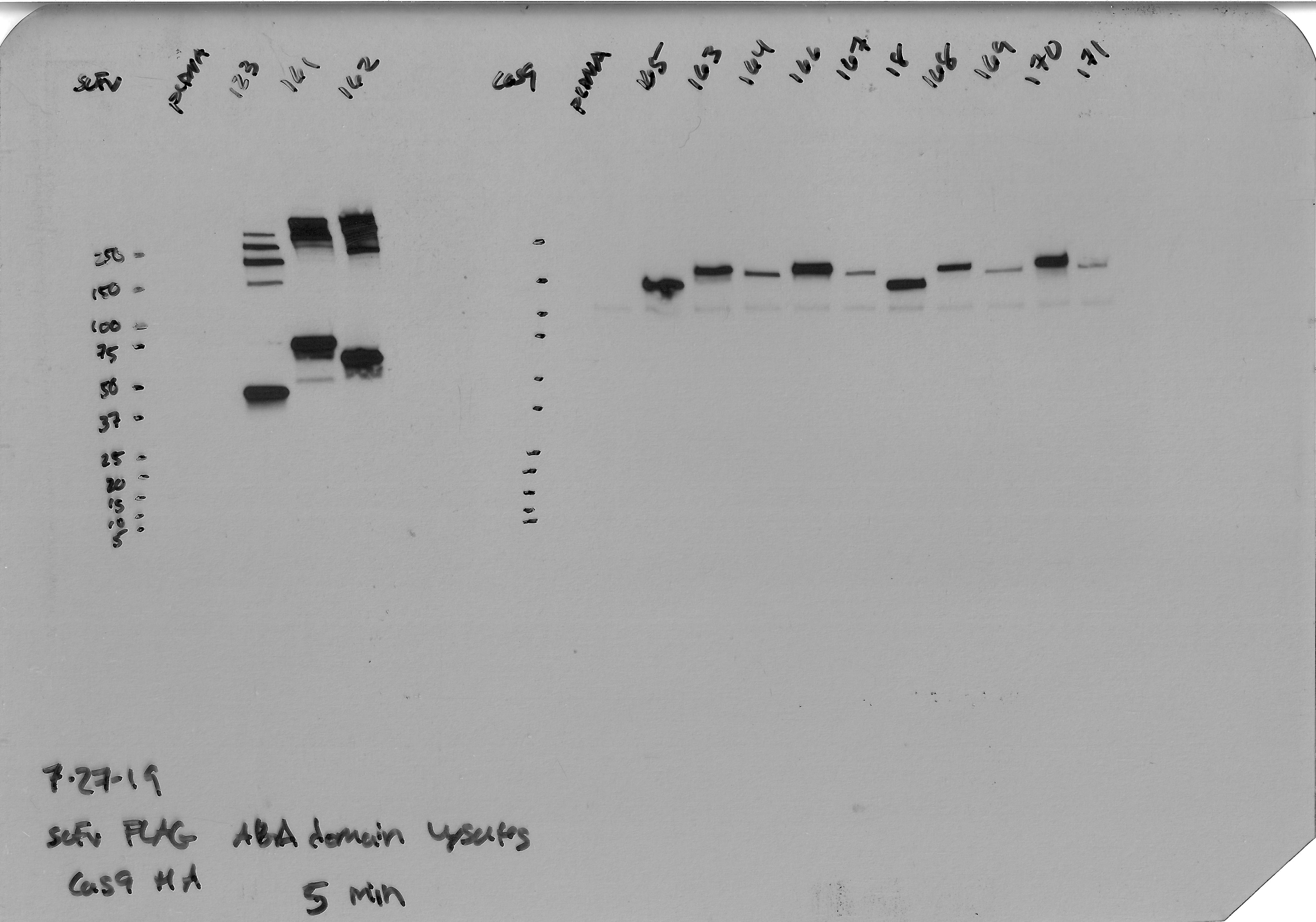

### Supplementary Fig 13e.tif

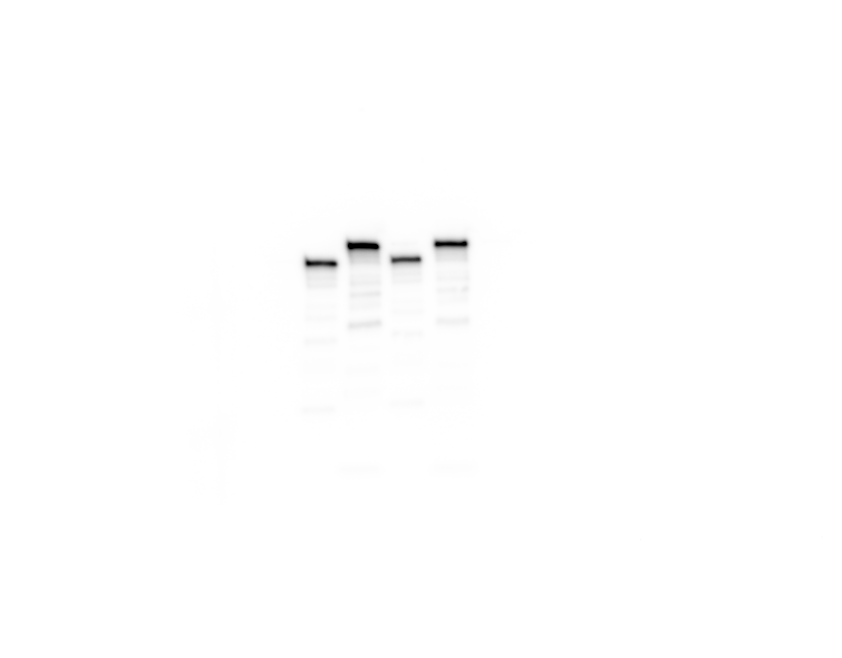

### Supplementary Fig 14a (bottom left).jpg

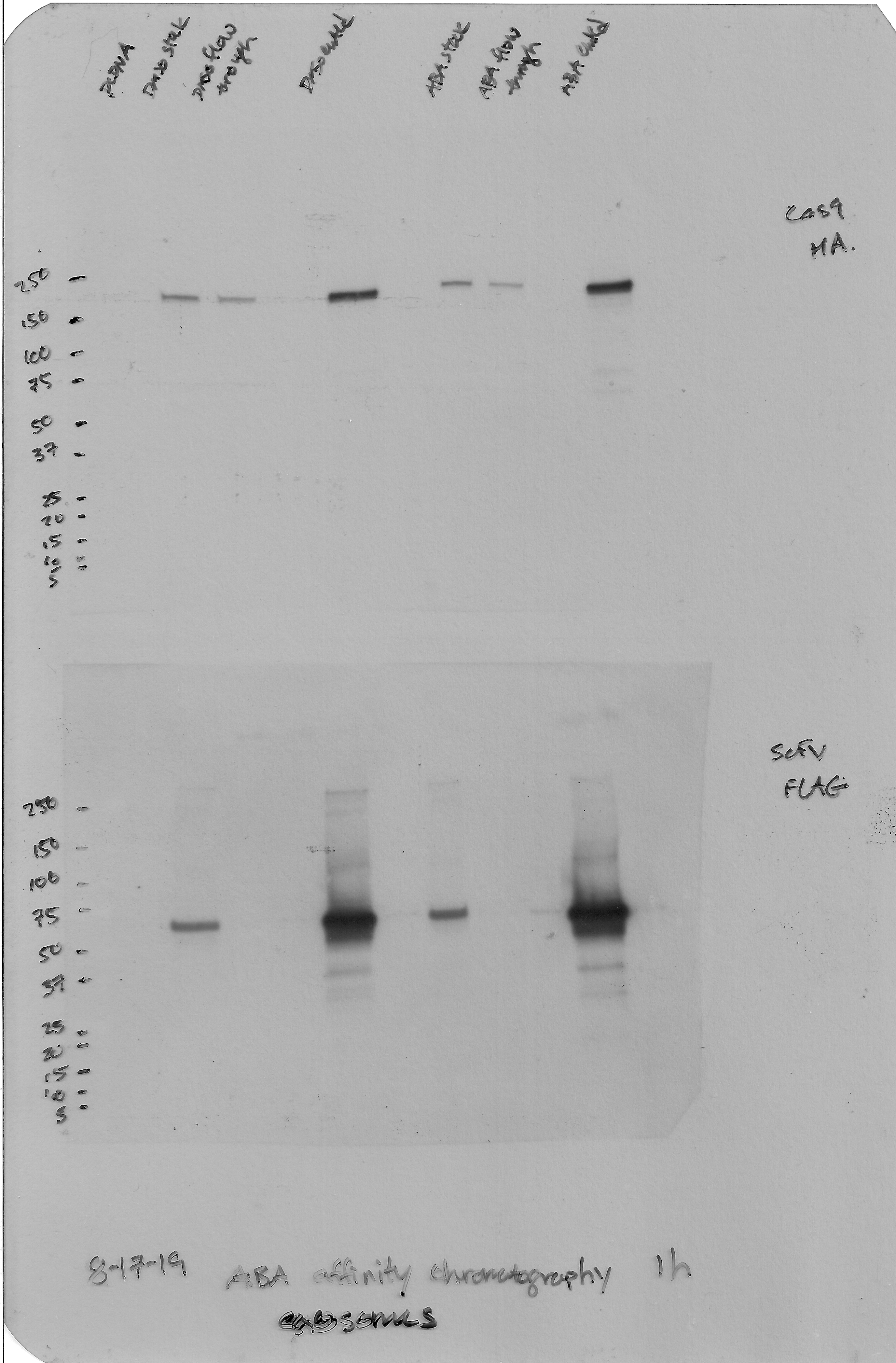
